# Two distinct SWI/SNF complexes direct chromatin-linked transcriptional programs in *Toxoplasma*

**DOI:** 10.1101/2025.07.16.665172

**Authors:** Dominic Schwarz, Benicio Tapia, Sebastian Lourido

## Abstract

Chromatin remodeling complexes dynamically modify DNA accessibility to mediate changes in gene expression during eukaryotic cell cycle progression, developmental transitions, and environmental adaptation. Higher eukaryotes have multiple remodeler subtypes based on the incorporation of different ATPases; however, the coordination and functional specificity of these diverse complexes is not well understood. Apicomplexan parasites such as *Toxoplasma gondii* have a limited set of chromatin remodelers offering a divergent setting in which to explore the function of homologous complexes. These parasites have selectively retained the Myb domain–containing proteins with homology to chromatin-associated regulators like SNF2H and SWI3. Here, a comprehensive analysis of the Myb protein family in *Toxoplasma* defines the composition of two SWI3 complexes defined by mutually exclusive ATPases with homology to the widely conserved BRG1 and BRM. Integrating transcriptomics with a custom chromatin-profiling strategy, we show that BRG1 is essential for the timely transcription of genes during mitosis and cytokinesis, while BRM ensures global transcriptional competency and fidelity throughout the cell cycle and developmental transitions. Our findings demonstrate that BRG1 and BRM perform distinct yet interdependent regulatory roles shaped by their chromatin context. This work uncovers ancestral principles of chromatin regulation and offers new insight into the functional diversification of SWI/SNF complexes across eukaryotes.

## INTRODUCTION

Apicomplexan parasites rely on intricate gene-expression programs, orchestrated by DNA-binding proteins and chromatin regulators, to colonize hosts and evade immune responses^1–8^. In *Toxoplasma gondii*, a model apicomplexan parasite, broad transcriptional changes shape key developmental transitions. During the lytic cycle, *Toxoplasma* tachyzoites undergo a tightly regulated cell cycle consisting of G1a, G1b and overlapping S, M and C phases. G1a and G1b constitute roughly half of the cell cycle, dedicated to cellular growth and preparation for DNA replication. S phase initiates chromosome duplication, organelle biogenesis, and centriole replication, while the formation of daughter cells begins late in S phase and proceeds through closed mitosis. These sequential events are driven by transcriptional programs, with successive waves of gene expression ensuring the timely production of factors essential for daughter cell formation^9–11^. In addition to cell cycle-linked changes, large-scale transcriptional shifts define developmental stage transitions as parasites differentiate to chronic bradyzoites, sexual stages, or environmentally resilient oocysts necessary for transmission and recombination^12^. Together, these dynamic expression programs enable *Toxoplasma* to balance rapid proliferation with long-term persistence across diverse host environments.

DNA-binding transcription factors have been extensively studied as key regulators of gene expression. In particular, AP2 (Apetala2/ethylene response factor) transcription factors have received considerable attention due to their lineage-specific expansion in apicomplexans and their critical roles in controlling developmental transitions and cell cycle progression^13^. However, chromatin structure constrains access to regulatory DNA elements and is shaped by nucleosome positioning, histone modifications, and higher-order chromatin organization^14^. Chromatin transitions frequently precede transcriptional changes, with certain regulatory boundaries becoming detectable at the chromatin level before transcription is initiated^6^. Both conserved and parasite-specific histone modifications have been reported in apicomplexan parasites which are tightly regulated by histone-modifying enzymes^7,15,16^. Yet, the functional landscape of chromatin readers—proteins that recognize and interpret histone modifications to mediate downstream regulatory processes—remains largely unexplored in apicomplexans, as current models of chromatin regulation primarily emphasize the recruitment of remodeling complexes through interactions with DNA-binding AP2 transcription factors^7,8,17^. The mechanisms by which these factors access regulatory DNA regions remain poorly understood, highlighting a major gap in our understanding of gene regulation in these parasites.

Our recent work pointed to Myb domain–containing proteins as compelling candidates for alternative chromatin regulatory pathways in *T. gondii*. Although some Myb proteins, such as *Toxoplasma* BFD1, *Cryptosporidium* Myb-M, and *Plasmodium* Myb1, retain classical DNA-binding functions, apicomplexans have predominantly retained Myb proteins of the SANT type, which lack DNA-binding capacity due to their negatively charged electrostatic surface^3,18–20^. SANT domains are commonly found in chromatin remodeling complexes, where they mediate interactions with histone tails and regulatory cofactors^18^. This suggests that apicomplexans may rely more heavily on chromatin-based regulatory mechanisms, with SANT-type Mybs potentially playing a central role in chromatin remodeling and gene regulation.

In the present study, we comprehensively analyze the Myb domain–containing proteins of *T. gondii* and identify nine as essential with diverse phenotypes and transcriptional signatures. Among those, the SWI3 ortholog emerged as a central regulator that assembles into two distinct chromatin-associated complexes with broad genomic distribution and profound impacts on gene expression. Both complexes are distinguished by incorporation of either BRG1 or BRM, SWI/SNF helicases that confer catalytic activity and establish regulatory potential. In canonical models, SWI/SNF complexes are thought to regulate chromatin accessibility by coupling ATP hydrolysis to nucleosome repositioning or eviction, thereby modulating gene expression programs that control fundamental processes such as differentiation, proliferation, and DNA repair^21^. In *Toxoplasma*, these remodelers similarly contribute to key biological transitions with *Tg*BRG1- and *Tg*BRM-containing complexes regulating cell cycle progression and developmental stage transitions, establishing an alternative layer of gene regulation critical for parasite development.

## RESULTS

### A diverse array of Myb domain–containing proteins is essential for the *Toxoplasma* lytic cycle

*Toxoplasma gondii* encodes 15 Myb domain–containing proteins, of which five contain canonical DNA-binding Myb-type domains and ten contain SANT-type domains typically associated with chromatin-related functions^18^. While Myb-type proteins include the master regulator of chronic differentiation BFD1, the SANT type group contains orthologs of chromatin-associated factors such as Ada2 (Ada2a: TGME49_217050, Ada2b: TGME49_262420), SWI3 (TGME49_286920), RCOR1 (TGME49_321450), DMAP1 (TGME49_213890) or the recently characterized SNF2h (TGME49_321440) (**Fig. 1A**)^3,8,18^. To systematically characterize their role during the *Toxoplasma* lytic cycle, we utilized a high-throughput tagging strategy, paired with a minimal auxin-inducible degron (mAID) to profile subcellular localization and growth defects following auxin-induced conditional knockdowns (cKD; **Fig. 1B**)^22^. BFD1 was excluded as it has been extensively characterized^3^.

**Figure 1.**
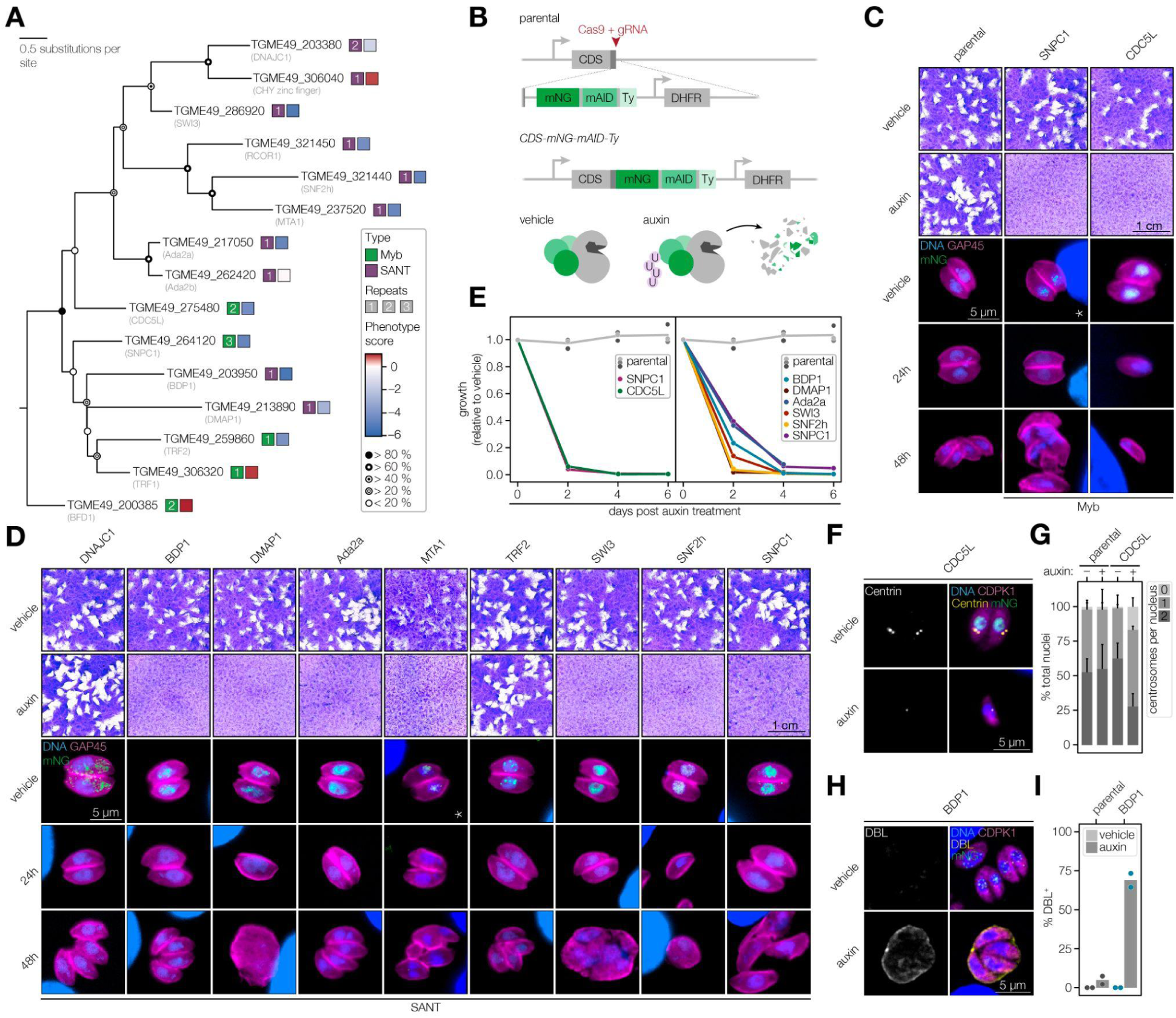
A high-throughput tagging strategy reveals functional diversity of *Toxoplasma* Myb domain–containing proteins. **A.** Phylogeny of *Toxoplasma* Myb domain–containing proteins^18^; DNA-binding Myb-type proteins (green squares) and SANT-type Myb proteins (purple squares). Numerals within each square denote the number of Myb domain repeats. The right square indicates the phenotype score derived from a genome-wide CRISPR screen, reflecting the relative fitness contribution of each gene^23^. Bootstrap values of 1000 replicates are represented by circles. **B.** Generation of conditional knockdown strains by C-terminal tagging of the coding sequence (CDS) with high-throughput tagging vectors^22^. Tagging with mNG-mAID-Ty targets proteins for degradation upon auxin treatment. DHFR denotes a pyrimethamine-resistance cassette to enable mutant selection. **C–D.** Plaque assays and representative vacuoles after 24 hr and 48 hr of knockdown of parental and Myb proteins (C) or SANT proteins (D). Fixed parasites were probed with antibodies against GAP45 (magenta) and mNG or ALFA-tag (green) as well as Hoechst dye to label DNA (blue). Asterisk denotes a strain tagged with 2×ALFA due to insufficient signal from mNG constructs. **E.** Competition assays of fitness-conferring Myb proteins (left) or SANT proteins (right) comparing the relative growth of mAID-tagged strains against the parental strain expressing IMC1-tdTomato^22^ in auxin-containing media. At the indicated time points, the proportion of fluorescent parasites within each competing population was measured by flow cytometry and normalized to the vehicle control. **F.** Representative images of cell cycle arrest following TGME49_275480 (CDC5L) knockdown. Parasites were fixed and stained with antibodies against Centrin (yellow), CDPK1 (magenta), and mNG (green), and DNA was visualized with Hoechst (blue). **G.** Quantification of centrosomes per parasite nucleus of CDC5L knockdown parasites. Centrosomes were quantified based on immunofluorescence analysis (F). Data represent the mean ± s.d. from two independent experiments. **H.** Representative images of the BDP1 differentiation phenotype following knockdown. Parasites were fixed and stained with antibodies against CDPK1 (magenta), and mNG (green), DNA was visualized with Hoechst (blue); the cyst wall was stained with DBL (yellow). **I.** Quantification of DBL-positive vacuoles of BDP1 knockdown parasites. Vacuoles were quantified based on immunofluorescence analysis (H). Data represent the mean from two independent experiments.

We examined all Myb domain–containing proteins that were found to be fitness conferring in a genome-wide CRISPR screen^23^. Out of the eleven fitness-conferring Myb proteins, nine displayed clear growth defects by plaque assay, suggesting essential roles during the lytic cycle (**Fig. 1C–D**). Notably, TGME49_203380, a homolog of the co-chaperone DNAJC, did not localize to the nucleus, while TGME49_259860, a homolog of Telomeric Repeat-binding Factor 2 (TRF2), exhibited a distinct sub-nuclear localization—patterns that are consistent with their predicted functional homology (**Fig. 1D**)^18,24,25^. The absence of a plaquing defect for TGME49_203380 and TGME49_259860 indicates that these proteins may confer only a subtle fitness defect, detectable primarily under competitive growth conditions. To further assess the impact of Myb protein depletion on parasite fitness, we performed competition assays by co-culturing each cKD line with the parental strain expressing IMC1-tdTomato^22^ in the presence of auxin. The resulting population dynamics closely mirrored the severity of the phenotypes observed by microscopy. Mutants exhibiting pronounced morphological defects or cell cycle arrest dropped out rapidly, whereas those with milder or delayed phenotypes displayed more gradual declines (**Fig. 1E; Supplementary Table 1**). Knockdown of nuclear and essential Myb proteins caused aberrant cell morphologies with two exceptions. Depletion of TGME49_275480—a homolog of the spliceosome component and cell cycle regulator CDC5L—resulted in a G1 phase arrest (**Fig. 1F–G; Supplementary Table 2**)^26–28^. By contrast, depletion of TGME49_203950—a homolog of Transcription factor TFIIIB component B (BDP1) and *Plasmodium* spp. High Mobility Group Protein B3 (HMGB3)^29^ led to the formation of tissue cyst-like structures under standard growth conditions (**Fig. 1H–I; Supplementary Table 3**).

The Myb proteins lacking fitness defects were assessed for their involvement in stress-induced differentiation. Quantifying cyst wall staining with *Dolichos biflorus* lectin (DBL), demonstrated that the three dispensable Myb proteins, along with TGME49_203380 and TGME49_259860—which showed no defects in plaque formation—showed no defects in differentiation (**Extended Data Fig. 1A–B; Supplementary Table 4**).

### INO80 and ISWI chromatin-regulator components share cell cycle signatures distinct from SWI/SNF

To place the Myb proteins into the hierarchy of a transcriptional network, we performed transcriptomic profiling following knockdown for 6 hr or 12 hr, focusing on those that were nuclear-localized and fitness-conferring. A clear separation between knockdown and control samples could be observed by principal component analysis (PCA), which captures the global transcriptional effects of Myb depletion (**Fig. 2A, Extended Data Fig. 2A–C; Supplementary Table 5**). Knockdown of Myb proteins with homology to transcription factors, such as SNPC4 and BDP1 caused only modest transcriptional changes. By contrast, depletion of Myb proteins related to chromatin regulation and the highly conserved spliceosome-associated factor CDC5L resulted in widespread transcriptional dysregulation (**Fig. 2A–C**)^27^.

**Figure 2.**
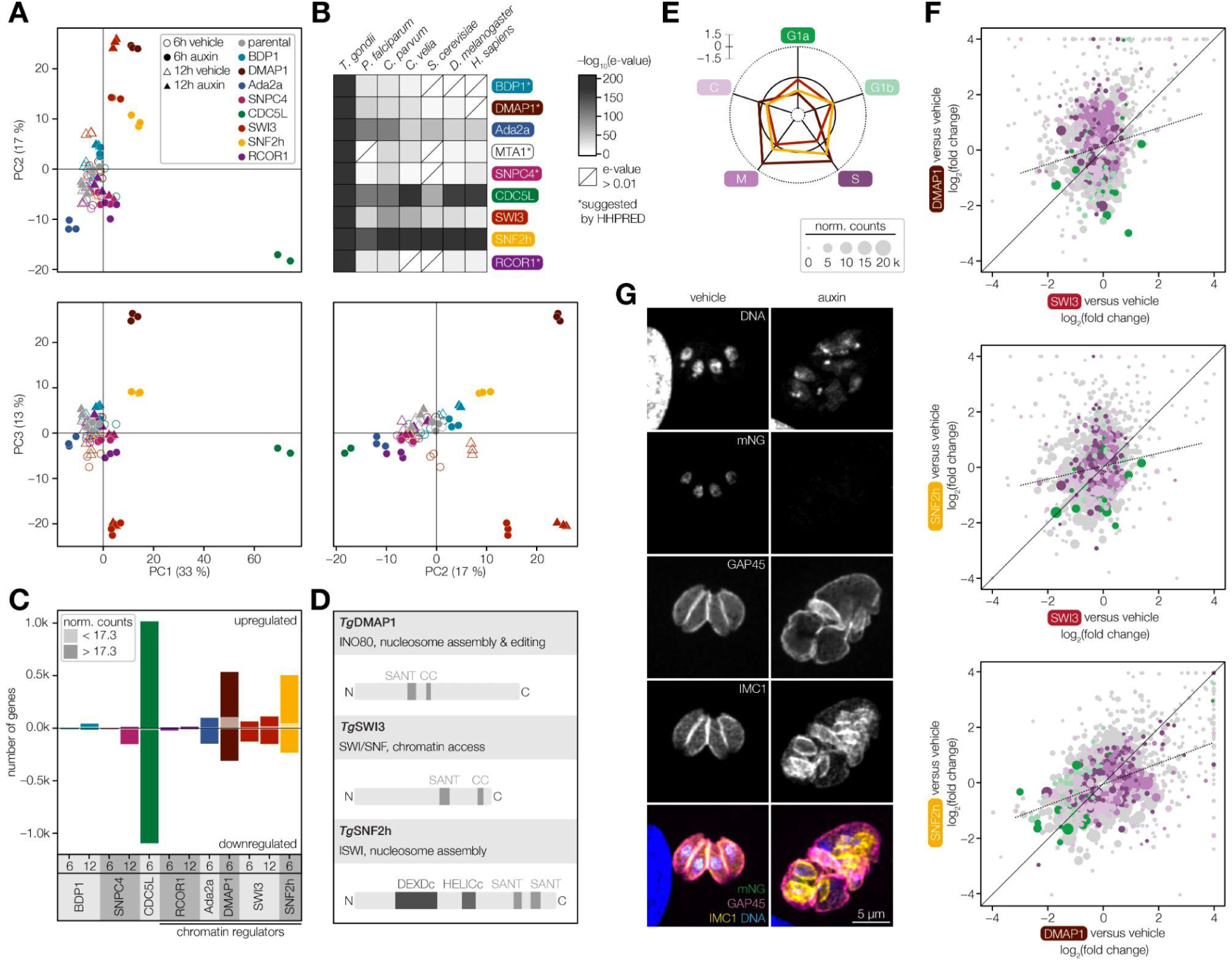
Myb proteins associated with chromatin remodeling differentially impact the *Toxoplasma* cell cycle. **A.** PCA was performed on the 500 most variable genes across all samples, based on DESeq2^77^ variance-stabilized expression values from biological triplicates for each condition and time point. The first three principal components (PC1–PC3) are shown, representing the major axes of transcriptional variation. **B.** Conservation analysis of full-length protein sequences for Myb proteins in apicomplexans (*T. gondii*, *Plasmodium falciparum*, and *Cryptosporidium parvum*), the chromerid *Chromera velia*, and the reference eukaryotes *Saccharomyces cerevisiae*, *Drosophila melanogaster*, and *Homo sapiens*. For each protein and species, the lowest e-value hit was reported to indicate sequence conservation; hits exceeding the 0.01 threshold were visually marked to reflect lack of significant homology between homologs. **C.** Bar plot displaying the number of significantly up-regulated and down-regulated genes following knockdown of individual Myb proteins, as determined by DESeq2^77^. Differentially expressed genes were defined by an adjusted *p*-value below 0.005 and an absolute fold change greater than 2. The proportion of genes with low read counts and Myb proteins with predicted roles in chromatin regulation are annotated. **D.** Schematic representation of the domain organization for three Myb proteins associated with ATP-dependent chromatin remodeling. Domains are annotated according to the SMART database^81^ and include SANT domains, ATP-dependent helicase domains (DEXDc & HELICc), and coiled coil (CC) regions. **E.** Transcriptomic impact of ATP-dependent chromatin remodeler components on cell cycle–regulated gene expression. Mean log₂ fold change in RNA abundance of gene sets associated with distinct cell cycle phases significantly affected following 6 hr of knockdown. Cell cycle gene sets were based on Xue et al.^37^. **F.** Pairwise comparisons of transcriptomic changes following 6 hr knockdown of proteins belonging to three distinct ATP-dependent chromatin remodeling complexes. Scatter plots display log₂ fold changes from RNA-seq data of genes with an adjusted *p*-value < 0.005. Genes are color-coded according to their cell cycle–regulated classification as shown in E. Dot size corresponds to the mean RNA-seq counts in vehicle-treated samples of the corresponding strains. **G.** Representative immunofluorescence image of *Tg*SWI3 after 48 hr of knockdown. Parasites were stained with antibodies against GAP45 (magenta), mNG (green), and IMC1 (yellow) as well as Hoechst for DNA (blue).

We next compared Myb proteins with homology to chromatin regulatory components. Comparative analysis showed SNF2h is well conserved among apicomplexans, yeast, fruit fly, and humans, while Ada2a shows moderate conservation, and DMAP1 and SWI3 are more divergent, with limited sequence similarity (**Fig. 2B**). PCA analysis showed that knockdown of *Tg*SNF2h and *Tg*DMAP1 induced comparable transcriptional responses, whereas depletion of *Tg*SWI3 led to a distinct expression profile (**Fig. 2A**). *Tg*SNF2h or *Tg*DMAP1 knockdown triggered mostly transcriptional upregulation, while *Tg*SWI3 knockdown predominantly resulted in downregulation, underscoring differences in their regulatory impact (**Fig. 2C**). Structural features further indicate functional specialization. *Tg*SNF2h contains the conserved ATPase domain characteristic of the catalytic core of the ISWI complex^8^. By contrast, *Tg*DMAP1 and *Tg*SWI3 lack ATPase domains and are presumed to function as cofactors within the INO80 and SWI/SNF complexes, respectively (**Fig. 2D**)^30^.

Despite shared involvement in chromatin remodeling, the transcriptional signatures of *Tg*SNF2h, *Tg*DMAP1, and *Tg*SWI3 diverged substantially, particularly on genes expressed during specific stages of the parasite’s cell cycle. Knockdown of *Tg*SNF2h or *Tg*DMAP1 primarily caused upregulation of S/M-specific transcripts and downregulation of G1a-specific transcripts, while *Tg*SWI3 depletion led to global downregulation of cell cycle–specific transcripts (**Fig. 2E–F**). Consistent with their transcriptional impacts, depletion of *Tg*DMAP1 and *Tg*SNF2h resulted in similar phenotypes, characterized by cell cycle arrest and disrupted nuclear morphology (**Fig. 1D**). By contrast, parasites depleted of *Tg*SWI3 retained the ability to undergo nuclear division but failed to complete cytokinesis. Chromatin in the *Tg*SWI3-depleted multinucleate cells appeared less dense, based on reduced Hoechst staining, suggesting compromised chromatin organization (**Fig. 2G**).

To investigate whether *Tg*SNF2h, *Tg*DMAP1, or *Tg*SWI3 functionally cooperate with known transcriptional repressors, we explored their relationship with the *Tg*MORC complex, a well-established repressive chromatin modifier directed by AP2 transcription factors^7^. Reanalysis of published *Tg*MORC ChIP-seq data revealed distinct correlations between *Tg*MORC occupancy at promoter regions and the transcriptional impact of different chromatin regulators. Specifically, genes upregulated upon *Tg*DMAP1 knockdown exhibited higher *Tg*MORC enrichment at promoter regions. By contrast, *Tg*SNF2h displayed the opposite trend: downregulated genes were associated with elevated *Tg*MORC occupancy, while upregulated transcripts corresponded to loci with reduced *Tg*MORC binding (**Extended Data Fig. 3A–D; Supplementary Table 6**). Together, these findings point to a functional divergence among ATP-dependent chromatin remodelers in *T. gondii*. While ISWI- and INO80-associated factors (*Tg*SNF2h and *Tg*DMAP1) display opposing dependencies on the repressive *Tg*MORC complex, *Tg*SWI3 exhibits a transcriptional profile distinct from both and consequently a more complex relationship with *Tg*MORC occupancy.

### *Tg*SWI3 defines two distinct SWI/SNF complexes distinguished by two different ATPases

SWI3 acts as a central scaffold in canonical SWI/SNF chromatin remodeling complexes, which regulate DNA accessibility for transcription, replication, and DNA repair. These remodelers function through the coordinated action of distinct modules—including ATPase and substrate recognition domains, that couple ATP hydrolysis to nucleosome repositioning or eviction^21,31^. To understand how SWI/SNF chromatin regulators operate in apicomplexan parasites, we set out to define the composition of the *Toxoplasma* SWI/SNF complex by co-immunoprecipitation mass spectrometry (co-IP MS) of *Tg*SWI3 endogenously tagged with mNG. Comparison of pulldowns with an untagged control revealed a network of *Tg*SWI3-associated proteins including several factors homologous to conserved components of the SWI/SNF complex such as SS18 (TGME49_226900), SMARCD1 (TGME49_248120), BRD4 (TGME49_306460), ALP2a (TGME49_258050), SMARCB1 (TGME49_226350), and ARID (TGME49_246170). Notably, the co-IP included two DEAD-box helicases: TGME49_278440 and TGME49_320300. Reciprocal IPs with either helicase confirmed their interaction with *Tg*SWI3 and several other SWI/SNF complex homologs (**Fig. 3A; Supplementary Table 7**).

**Figure 3.**
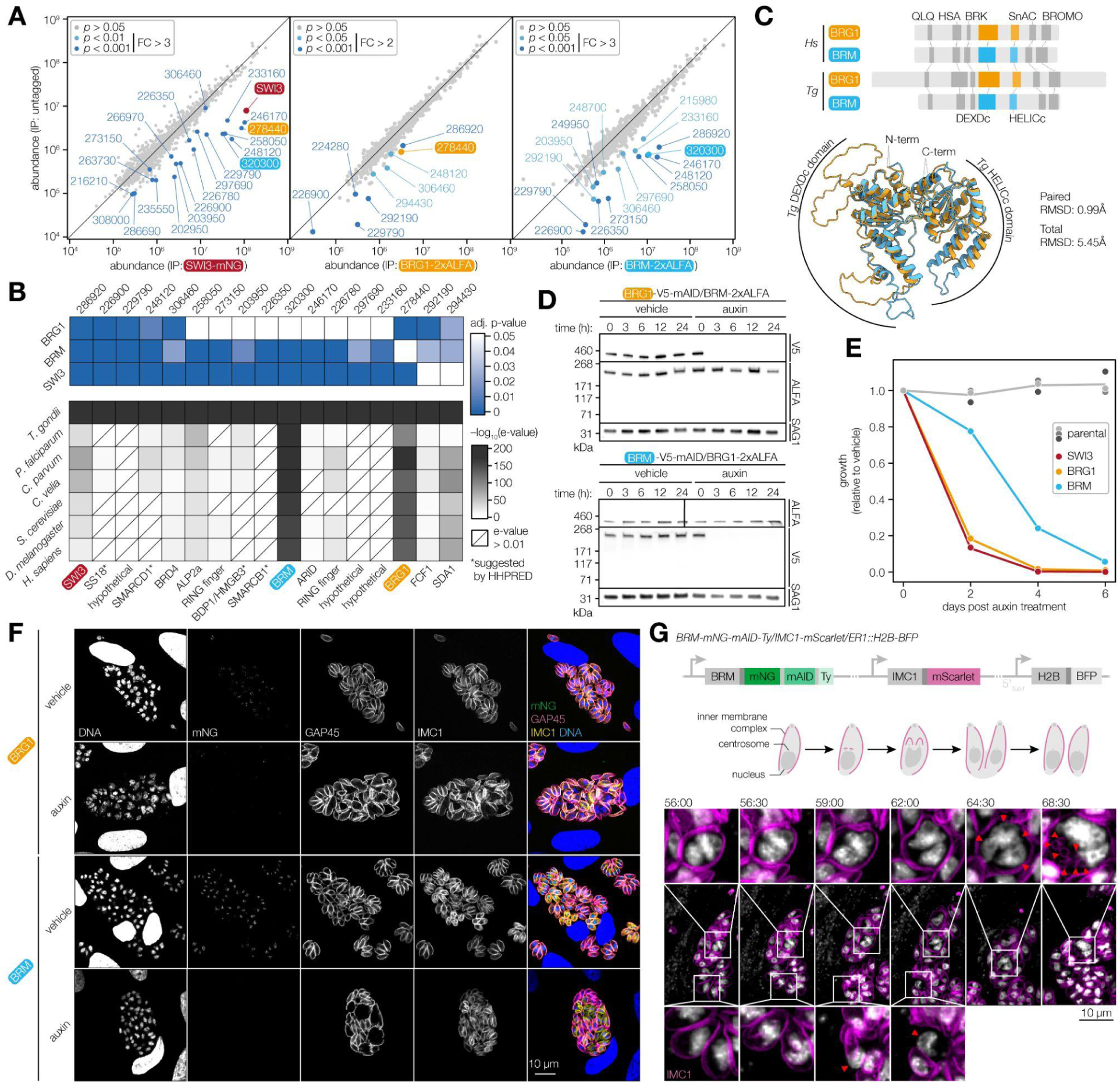
Two SWI/SNF chromatin remodelling complexes are defined by their different helicases *Tg*BRG1 or *Tg*BRM. **A.** Co-IP MS of endogenously tagged *Tg*SWI3 with mNG and reciprocal co-IP MS of endogenously tagged *Tg*BRG1-2×ALFA and *Tg*BRM-2×ALFA. Abundances of proteins identified with > 2 unique peptides are shown for *n* = 3 biological replicates. Significantly enriched proteins are colored in blue according to their *p*-value by Benjamini-Hochberg corrected ANOVA. **B.** Heatmap of proteins co-purified with *Tg*SWI3, *Tg*BRG1 or *Tg*BRM as in A, showing only proteins that were significantly enriched in two of the three pulldowns. Tiles are colored by their adjusted *p*-value in blue or by their conservation in representative genomes. For each protein and species, the lowest e-value hit after homology searches is reported. **C.** Schematic representation of domain architectures identified by MSA based on **Extended Data Fig. 5B**, alongside superimposed AlphaFold3^33^ structural predictions of the DEXDc and HELICc domain regions from *Tg*BRG1 (orange) and *Tg*BRM (blue). The overall structural similarity yields a RMSD of 5.45 Å, while the individual DEXDc and HELICc domains exhibit lower RMSDs of 2.42 Å and 0.54 Å, respectively. **D.** Immunoblot analysis of *Tg*BRG1 and *Tg*BRM stability during reciprocal knockdown. Parasite lines expressing either *Tg*BRG1-mAID with *Tg*BRM-2×ALFA (top) or *Tg*BRM-mAID with *Tg*BRG1-2×ALFA (bottom) were treated with vehicle or auxin for 3, 6, 12, or 24 hr to induce conditional knockdown. Blots were probed with anti-ALFA to monitor the abundance of the non-degraded helicase, anti-V5 to confirm the knockdown, and anti-SAG1 as loading control. **E.** Competition assays of *Tg*SWI3, *Tg*BRG1, and *Tg*BRM mAID-tagged strains comparing the relative growth against the parental strain expressing IMC1-tdTomato^22^ in auxin-containing media. At each time point, the proportion of fluorescent parasites within each competing population was measured by flow cytometry and normalized to the vehicle control. **F.** Representative immunofluorescence images of *Tg*BRG1 and *Tg*BRM knockdown parasites following 72 hr of auxin or vehicle treatment. Fixed parasites were stained for GAP45 (magenta), IMC1 (yellow), and mNG (green) as well as Hoechst to label DNA (blue). **G.** Cell cycle reporter strain engineered for live-cell imaging. IMC1 was endogenously tagged at the C terminus with mScarlet-I to visualize daughter cell formation and H2B fused to BFP was ectopically expressed to visualize nuclear dynamics. A schematic of wildtype *T. gondii* endodyogeny is shown for reference. Live-cell imaging of *Tg*BRM knockdown parasites imaged over a 24-hr period beginning 48 hr after auxin-induced knockdown. Selected frames from 56 to 68.5 hr post-knockdown highlight distinct phenotypes by red triangles: nuclear division without subsequent cytokinesis (top) and mitosis and cytokinesis followed by failure of nuclear retention in daughter cells (bottom). See also **Video S1**.

Several proteins identified in the co-IPs were unexpected constituents of the putative SWI/SNF complexes. TGME49_229790, TGME49_297690, and TGME49_233160 lacked homologs beyond the closest relatives of *Toxoplasma*, suggesting parasite-specific adaptations of the SWI/SNF complex (**Fig. 3B**). TGME49_320300 and *Tg*SWI3 co-IP’d with the Myb protein TGME49_203950 (*Tg*BDP1). However, interactions between SWI/SNF complexes and BDP1 have not been previously reported and BDP1 is not typically associated with HMG domains^21,32^. Closer inspection revealed that all peptides identified in the *Tg*SWI3 pulldown mapped exclusively to the HMG domain–containing N-terminus of the gene model, with no coverage of the annotated C-terminal Myb domain. The uneven distribution of RNA-seq reads across the locus, peaking in the middle of the annotated gene model, further suggest the current annotation is inaccurate and the locus produces two distinct transcripts (**Extended Data Fig. 4A**). Experimental validation confirmed that TGGT1_203950 indeed gives rise to two distinct transcripts, encoding separate proteins that we will refer to as *Tg*HMGB3 for the N-terminal portion and *Tg*BDP1 for the C-terminal portion (**Extended Data Fig. 4B**-D), and only *Tg*HMGB3 interacts with the *Tg*SWI3 complex.

The two helicases (TGME49_278440 and TGME49_320300) stand out as the best-conserved elements across the analyzed eukaryotic lineages (**Fig. 3B**). Phylogenetic analysis of the 17 most closely related (e-value cutoff: 0.01) *Toxoplasma* proteins containing both DEXDc and HELICc domains revealed distinct clades with homology to canonical chromatin remodeler families, including CHD, ISWI, and INO80, as well as RAD-like ATPases involved in DNA repair (**Extended Data Fig. 5A**). The two helicases identified in the *Tg*SWI3 co-IPs were closely related members of the SWI/SNF clade and show highest sequence similarity to BRG1 (TGME49_278440) and BRM (TGME49_320300), respectively (**Extended Data Fig. 5B–C**). Others have also suggested these two proteins are candidate SWI/SNF helicases based on bioinformatic analyses^8,17^. Alignment with SWI/SNF helicases from other eukaryotes identified all of the hallmark domains including QLQ, HSA, BRK, SnAC and BROMO domains (**Fig. 3C; Extended Data Fig. 5B**). Furthermore, *Tg*BRG1 and *Tg*BRM are predicted to adopt similar overall folds within their ATPase cores, despite notable differences in their predicted structures (AlphaFold3^33^ RMSD: 5.45 Å) and primary sequence (42.18 % identity; **Fig. 3C**).

We considered whether depletion of one helicase might destabilize or co-deplete the other^31,34^. To test this, we generated dually-tagged parasite lines in which either *Tg*BRG1 or *Tg*BRM was endogenously fused to an mAID, while the reciprocal helicase was tagged with a 2×ALFA epitope, enabling depletion of one helicase and monitoring of the other. We assayed *Tg*BRG1 and *Tg*BRM abundance following different periods of knockdown. Immunoblot analysis showed that depletion of either *Tg*BRG1 or *Tg*BRM did not impact the stability or abundance of the other. These results indicate that the two SWI/SNF subcomplexes remain independently assembled and stable upon loss of the alternate helicase (**Fig. 3D**), consistent with the formation of discrete *Tg*SWI3-associated chromatin remodeling complexes incorporating either *Tg*BRG1 or *Tg*BRM as observed in mammals^31,34^.

### Functional divergence between *Tg*BRG1 and *Tg*BRM reveals non-redundant roles in *Toxoplasma*

We assessed the individual contributions of *Tg*BRG1 and *Tg*BRM to parasite fitness by performing competition assays in the presence of auxin. These experiments revealed that *Tg*BRG1-depleted parasites were rapidly outcompeted by the wildtype parental line with similar kinetics to *Tg*SWI3-depleted parasites^22^. By contrast, *Tg*BRM-depleted parasites displayed a comparatively milder phenotype, suggesting a more acute functional requirement for *Tg*BRG1 during the *Toxoplasma* lytic cycle (**Fig. 3E; Extended Data Fig. 6A; Supplementary Table 8**).

To further dissect the functional consequences of depleting the two SWI/SNF helicases, we performed immunofluorescence microscopy following 72 hr of *Tg*BRG1 or *Tg*BRM knockdown. Strikingly, depletion of *Tg*BRG1 resulted in severe cell division defects, characterized by aberrant nuclear morphology and pronounced changes in Hoechst staining intensity, indicative of altered chromatin structure and compaction^35^. By contrast, *Tg*BRM knockdown led to the emergence of parasites appearing to undergo endopolygeny, with more than two daughter cells forming within a single mother cell^36^. This phenotype was observed in ∼30 % of parasite vacuoles and often affected only a subset of parasites within the same vacuole, suggesting dysregulation of the parasite’s cell cycle (**Fig. 3F; Extended Data Fig. 6B; Supplementary Table 9**). These findings are consistent with non-redundant roles for *Tg*BRG1 and *Tg*BRM in orchestrating parasite division and chromatin architecture.

To observe the cell division phenotype associated with *Tg*BRM depletion by live-cell imaging, we generated a conditional *Tg*BRM knockdown strain expressing fluorescently tagged markers for the inner membrane complex (IMC1) and chromatin (H2B; **Fig. 3G**). Time-lapse microscopy showed that *Tg*BRM knockdown disrupts the coordination between mitosis and cytokinesis. In a subset of parasites, nuclear division proceeds but is not followed by cytokinesis, leading to the accumulation of multinucleated cells and subsequent formation of multiple daughters within a single mother cell. In other cases, daughter cells fail to retain their nucleus during cytokinesis, resulting in the presence of nuclei within the residual body (**Fig. 3G; Video S1**). These observations indicate that *Tg*BRM is critical for the faithful progression of nuclear segregation and cytokinesis during parasite replication.

### *Tg*BRG1 and *Tg*BRM depletion have distinct transcriptional outcomes

The distinct phenotypes resulting from *Tg*BRG1 and *Tg*BRM depletion suggest that the associated complexes contribute differentially to gene expression. To investigate their transcriptional impact, we performed transcriptional profiling following knockdown for 6, 12, or 24 hr. Loss of either helicase was predominantly associated with down-regulation of gene expression. Overall, *Tg*BRG1 knockdown led to a faster and more substantial transcriptional disruption compared to *Tg*BRM, underscoring the immediate dependence of gene expression on *Tg*BRG1 (**Fig. 4A; Supplementary Table 10**). Substantial transcriptional upregulation emerged after 12 hr of *Tg*BRG1 knockdown, although most of these genes were lowly expressed under baseline conditions and thus prone to high variance; accordingly, they were excluded from subsequent analyses (**Fig. 4A; Extended Data Fig. 7A**). PCA revealed that *Tg*SWI3 and *Tg*BRG1 knockdowns clustered closely on the first principal component (PC1, 41 % of variance), suggesting a shared transcriptional signature, while *Tg*SWI3 and *Tg*BRM knockdowns were more similar on the second component (PC2, 20 % of variance). Moreover, *Tg*BRM knockdown showed transcriptional progression over time, while *Tg*BRG1 and *Tg*SWI3 profiles remained relatively stable (**Fig. 4B, Extended Data Fig. 7B**). Together, these findings suggest that *Tg*BRG1 and *Tg*BRM operate in distinct temporal regimes, while *Tg*SWI3 integrates features of both complexes, reflecting its role as scaffold in the two SWI/SNF complexes.

**Figure 4.**
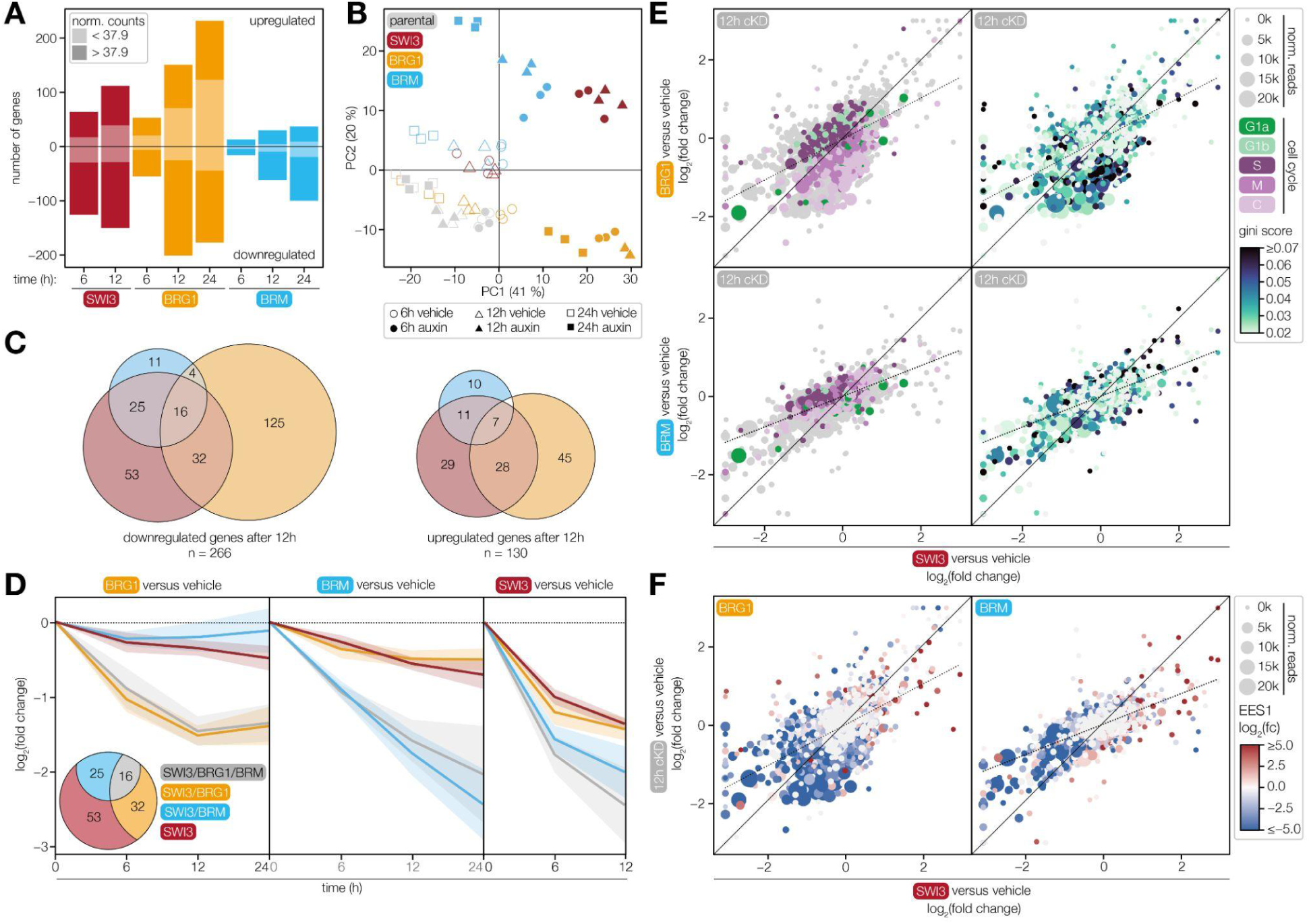
Time-resolved transcriptomic analysis reveals distinct regulatory roles for *Tg*BRG1 and *Tg*BRM. **A.** Bar plot displaying the number of significantly up-regulated and down-regulated genes following knockdown of *Tg*BRG1, *Tg*BRM, or *Tg*SWI3, as determined by DESeq2^77^. Differentially expressed genes were defined by an adjusted *p*-value below 0.005 and an absolute fold change greater than 2. The proportion of genes with low read counts (< 37.9) is annotated (**Extended Data Fig. 7A**). **B.** PCA was performed on DESeq2^77^ variance-stabilized expression values generated in biological triplicates for each condition and time point, using genes with a variance > 0.1 across all samples. Genes with low read counts (< 37.9) across all samples were excluded to minimize noise. The first two principal components (PC1–PC2) are shown, representing the major axes of transcriptional variation. **C.** Venn diagrams of the overlap of up- and down-regulated genes following a 12-hr knockdown of *Tg*BRG1 (orange), *Tg*BRM (blue), and *Tg*SWI3 (red). **D.** The expression trajectories of genes classified as downregulated after 12-hr *Tg*SWI3 knockdown were analyzed in each of the knockdowns. Gene expression changes were tracked at 6, 12 and 24 hr timepoints for *Tg*BRG1 and *Tg*BRM, and for 6 and 12 hr timepoints for *Tg*SWI3. Lines represent the mean log_2_ fold change at each timepoint; shaded areas indicate s.e.m. **E–F.** Pairwise comparisons of transcriptomic changes following 12-hr knockdown of *Tg*BRG1 or *Tg*BRM, compared to *Tg*SWI3. Scatter plots display log₂ fold changes from RNA-seq data. Genes are color-coded by Gini score based on Behnke et al.^39^, by their cell cycle–regulated classification based on Xue et al.^37^ (E), or by their log₂ fold change in EES1-stage parasites relative to tachyzoites based on Ramakrishnan et al.^60^ (F). Dot size corresponds to the mean RNA-seq counts in vehicle-treated samples of the corresponding strains.

We further investigated the temporal dynamics and specificity of transcriptional responses to disrupting the different SWI/SNF complexes. Venn diagrams of genes that were up- or down-regulated following 12 hr of knockdown for *Tg*BRG1, *Tg*BRM, or *Tg*SWI3 indicated that most differentially expressed genes were associated with the depletion of individual complex subunits (**Fig. 4C**). This pattern may result from overly conservative thresholds in differential-expression analysis (adjusted *p*-value < 0.005 and absolute fold change > 2), which might not capture weaker trends in gene expression. As an alternative analysis, we examined the behavior of all genes that were downregulated in the *Tg*SWI3 dataset across the various knockdowns (**Fig. 4D**). In *Tg*BRG1-depleted parasites, genes associated with *Tg*BRM-dependent down-regulation remained largely unaffected, while the *Tg*SWI3-associated subset showed moderate down-regulation. By contrast, depletion of *Tg*BRM resulted in the down-regulation of genes associated with both *Tg*BRG1 and *Tg*SWI3, although the magnitude of repression was less pronounced than for genes associated with *Tg*BRM based on magnitude and significance cutoffs. As expected, *Tg*SWI3 depletion caused widespread down-regulation across all three subsets. This pattern suggests an asymmetric codependence, with *Tg*BRG1-dependent genes requiring *Tg*BRM while *Tg*BRM-dependent genes are less reliant on *Tg*BRG1. Together, these findings support a model in which *Tg*BRM acts as a general transcriptional activator, while *Tg*BRG1 is selectively required for the activation of a distinct subset of genes that depend more acutely on its presence.

Prompted by the distinct cell division defects observed upon *Tg*BRG1 and *Tg*BRM depletion, we investigated how cell cycle–regulated genes respond to the loss of these remodelers. We compared transcriptional profiles at 12 hr post-knockdown relative to *Tg*SWI3 depletion. Consistent with our earlier observations, *Tg*BRM and *Tg*SWI3 knockdowns predominantly affected the same set of genes, though with distinct kinetics, with *Tg*SWI3 depletion triggering faster transcriptional downregulation (**Fig. 4E**). By contrast, *Tg*BRG1-sensitive genes were enriched for genes expressed during short-lived cell cycle stages such as mitosis and cytokinesis^37^. These genes strongly correlated with high Gini scores—a measure of distribution inequality that quantifies variation across samples ranging from genes expressed equally across all stages (Gini score = 0) to genes expressed uniquely at a single stage (Gini score = 1)^38^. Gini scores were calculated from published transcriptome data obtained from synchronized parasites^39^ (**Fig. 4E; Extended Data Fig. 7C**). This pattern suggests that *Tg*BRG1 is more critically required for the transcription of genes with narrow temporal expression windows, such as those activated during mitosis and cytokinesis.

Despite their distinct transcriptional profiles, *Tg*BRG1 and *Tg*BRM knockdowns share several upregulated genes, many of which are typically repressed in tachyzoites but activated in bradyzoites and early sexual stages (EES1, **Fig. 4F**; **Extended Data Fig. 7C**). Similar patterns of de-repression have been observed upon disruption of other ATP-dependent chromatin remodelers, such as *Tg*DMAP1, *Tg*SNF2h, and *Tg*MORC (**Extended Data Fig. 7D–E**)^7,8^. These findings suggest that while *Tg*BRG1 is required for the transcriptional activation of genes with narrow, phase-restricted expression windows, disruption of chromatin remodeling can lead to de-repression of developmentally silenced transcripts in tachyzoites.

### Chromatin profiling using nanobody-directed CUT&RUN

We sought to define the genome-wide binding profiles of the SWI/SNF complexes. Although chromatin profiling approaches such as ChIP-seq and CUT&RUN have been successfully applied in *Toxoplasma*, these methods typically require large numbers of parasites and are limited in their ability to capture transient or low-abundance interactions^3,6,8,40,41^. To overcome these limitations, we developed ALFA-CUT&RUN, a rapid and sensitive chromatin profiling approach optimized for *Toxoplasma,* extending the original CUT&RUN^42,43^. Our method leverages the strong and specific interaction between the ALFA epitope tag and its nanobody, NbALFA^44^. By directly fusing micrococcal nuclease (MNase) to NbALFA, the system eliminates the need for antibodies and enables targeted cleavage of chromatin in proximity to ALFA-tagged proteins.

To specifically map the binding sites of the various complexes, parasites expressing C-terminally 2×ALFA-tagged *Tg*SWI3, *Tg*BRG1, or *Tg*BRM were permeabilized, incubated with the NbALFA-MNase fusion protein, and subjected to calcium-dependent nuclease activation. Released DNA fragments were then purified and sequenced to generate high-resolution binding profiles of *Tg*SWI3, *Tg*BRG1 and *Tg*BRM across the genome for two biological replicates (**Fig. 5A; Extended Data Fig. 8**). To ensure specificity and assess background signal, we included two controls: the untagged parental strain, and a 2×ALFA-tagged *Tg*BDP1 line, which served as a positive control based on its well-characterized genomic distribution at short and non-coding RNAs in other systems^45^. This allowed us to distinguish specific enrichment patterns associated with *Tg*SWI3, *Tg*BRG1 and *Tg*BRM from background cleavage or nonspecific interactions. Although we found that all SWI/SNF subunits bound several *Tg*BDP1 target sites—highlighting their broad role in transcriptional regulation—these sites were excluded from downstream analyses so as to take full advantage of this control (**Extended Data Fig. 8D–F**). Following stringent differential-binding analysis, the signal intensities from biological replicates for *Tg*SWI3, *Tg*BRG1 and *Tg*BRM were strongly correlated (**Fig. 5B–C; Extended Data Fig. 8B–F**). These results underscore the robustness and specificity of the ALFA-CUT&RUN approach for profiling chromatin-associated proteins in *Toxoplasma*.

**Figure 5.**
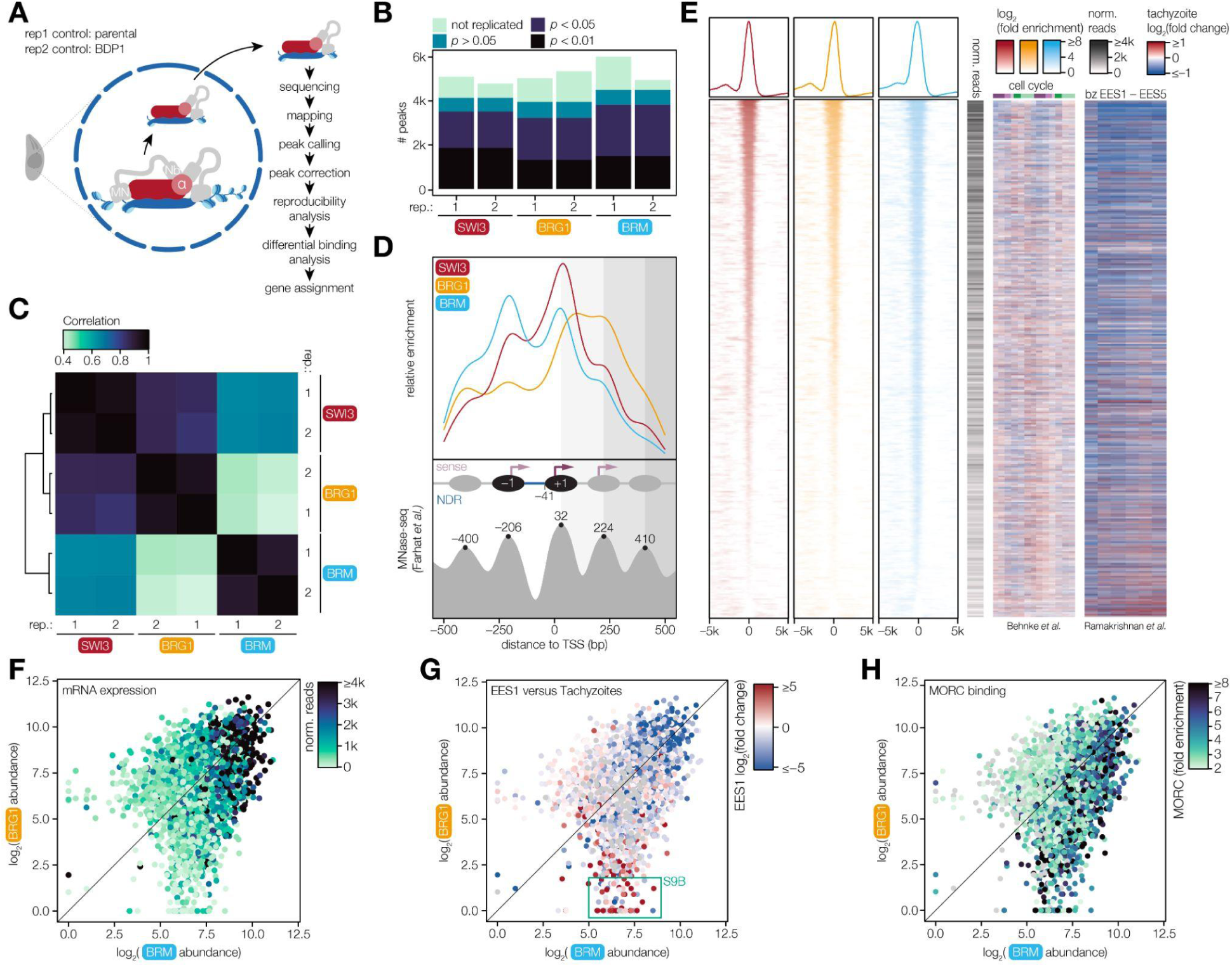
Genome-wide profiling of *Tg*BRG1 and *Tg*BRM occupancy reveals distinct promoter-binding patterns associated with developmental gene regulation. A. Overview of the ALFA-CUT&RUN protocol. The NbALFA-directed MNase is targeted to ALFA-tagged (α) proteins where it is cleaving the surrounding DNA, generating short fragments centered on the DNA-binding site of the protein of interest (red). Excised fragments diffuse out of the nucleus, allowing for selective enrichment, sequencing, and mapping. **B.** Number of peaks identified per ALFA-CUT&RUN replicate for *Tg*SWI3, *Tg*BRG1, or *Tg*BRM. Bars represent the total number of peaks called in each replicate, with replicated peaks indicated in a darker shade according to IDR-calculated *p*-values. **C.** Correlation heatmap of the ALFA-CUT&RUN signal across all samples and replicates for 4692 consensus loci between *Tg*SWI3, *Tg*BRG1 and *Tg*BRM samples. Pairwise Pearson correlation coefficients were calculated using normalized and background-subtracted read counts. **D**. Relative enrichment of *Tg*SWI3, *Tg*BRG1, or *Tg*BRM within ±500 bp of annotated TSSs. MNase-seq–defined nucleosome positions from Farhat et al.^7,82^ are highlighted to indicate phased nucleosome occupancy relative to the nearest TSS. **E.** Distribution of *Tg*SWI3 (red), *Tg*BRG1 (orange), and *Tg*BRM (blue) peaks ordered according to their average peak intensities, relative to the nearest TSS. Each row represents a single gene that could be confidently assigned to a specific peak based on proximity (4433 total peaks). Heatmaps display associated transcriptomic data: mean RNA expression in tachyzoites (greyscale); log₂ fold changes across cell cycle phases relative to tachyzoite mean expression, based on Behnke et al.^39^ (cell cycle stages indicated above with S, M, and C in shades of purple, and G1a and G1b in shades of green); and log_2_ fold changes in gene expression for tissue cysts and enteroepithelial stages (EES1–5) relative to tachyzoites, based on Ramakrishnan et al.^60^. **F–H.** Comparison of *Tg*BRG1 and *Tg*BRM log₂ abundance based on normalized read counts after background signal subtraction. Each point represents a single genomic consensus locus of the 4433 high confidence peak-gene pairs. Points are colored based on mean gene expression level in tachyzoites (F), by the log₂ fold change in expression of EES1 stages relative to tachyzoites based on Ramakrishnan et al.^60^ (G), or by *Tg*MORC fold enrichment based on Farhat et al.^7^ (H).

### *Tg*BRG1 and *Tg*BRM display binding patterns with distinct cell cycle and chromatin signatures

We applied the Irreproducible Discovery Rate (IDR) framework to identify loci consistently enriched across biological replicates^46^. Using an IDR *p*-value < 0.05, we identified 3,516 reproducible peaks for *Tg*SWI3, 3,229 for *Tg*BRG1, and 3,823 for *Tg*BRM (**Fig. 5B; Extended Data Fig. 8A**, D**–F; Supplementary Table 11**). Analysis of the genomic distribution of replicated peaks revealed that binding sites for *Tg*SWI3, *Tg*BRG1, and *Tg*BRM are evenly distributed across all chromosomes (**Extended Data Fig. 8G**), with specific enrichment at transcription start sites (TSSs; **Fig. 5D–E**). Nucleosome-level mapping further uncovered distinct binding patterns: *Tg*BRG1 was predominantly centered around the +1 and +2 nucleosomes downstream of the TSS, whereas *Tg*BRM was enriched at the –1 and +1 nucleosomes, flanking the nucleosome-depleted region (NDR; **Fig. 5D**). These findings suggest that the two SWI/SNF complexes engage with promoter architecture in distinct ways, potentially reflecting their specialized functions in gene regulation.

To enable the direct comparison of *Tg*BRG1 and *Tg*BRM binding across the genome, we defined a set of 4,692 consensus regions, representing all loci identified by an IDR *p*-value < 0.05 in any of the samples. Of these, 4,433 could be confidently assigned to a single promoter, providing a unified set of loci for quantitative analysis of differential binding (**Fig. 5E; Extended Data Fig. 8A**). Correlating *Tg*BRG1 and *Tg*BRM binding profiles with transcriptomic data across the cell cycle and life cycle stages revealed distinct regulatory patterns. Genes with the highest *Tg*BRG1 and *Tg*BRM occupancy were predominantly associated with G1a and G1b phases, whereas loci bound at lower intensity were more frequently linked to S, M, or C phase genes, which are often transiently expressed and require tight temporal control. Moreover, peak intensity was positively correlated with mRNA abundance, suggesting that SWI/SNF occupancy is linked to transcriptional output (**Fig. 5E–F; Supplementary Table 12**); a conclusion further supported by the trend of down-regulation following knockdown of the factors. *Tg*BRM bound several loci with reduced or absent *Tg*BRG1 binding (**Fig. 5E; Extended Data Fig. 8H**). These *Tg*BRM-enriched peaks were linked to genes that are transcriptionally silent during the lytic cycle but become strongly induced during chronic- and early sexual-stage differentiation, as well as in extracellular parasites (**Fig. 5E and G; Extended Data Fig. 9A–B**). One such differentiation-associated gene is *CST10*, which has been identified as an early chronic stage marker that is expressed before other chronic stage markers such as BAG1 or CST1^3^. This suggests that while *Tg*BRM is not sufficient on its own to drive transcriptional activation, it maintains these loci in a poised state for rapid activation during developmental transitions.

Given the association of *Tg*BRM-enriched loci with genes activated during developmental transitions, we compared *Tg*BRM and *Tg*BRG1 binding patterns to *Tg*MORC, a transcriptional repressor involved in silencing stage-specific genes (**Fig. 5H; Extended Data Fig. 9C–D**)^7^. *Tg*BRM was strongly correlated with *Tg*MORC occupancy, whereas *Tg*BRG1 exhibited no such correlation. Genomic profiling further revealed that *Tg*MORC, like *Tg*BRM, is predominantly localized at the –1 nucleosome upstream of the transcription start site, consistent with a role in modulating promoter accessibility (**Extended Data Fig. 9D**). Notably, a subset of genomic regions displayed high levels of *Tg*BRG1, *Tg*BRM and *Tg*MORC co-occupancy (**Fig. 5H**). These loci were associated with genes that are highly expressed in tachyzoites at the population level, indicating that the functional interplay between *Tg*BRG1, *Tg*BRM and *Tg*MORC extends beyond developmental regulation and also contributes to transcriptional control in tachyzoites.

### Late cell cycle stages exhibit an increased dependence on *Tg*BRG1

As noted above, *Tg*BRG1 and *Tg*BRM occupancy was higher for genes expressed during G1 than for genes expressed during the S, M, or C phases of the cell cycle. If *Tg*BRG1 and *Tg*BRM occupancy changes dynamically across the cell cycle, the observed pattern would be dominated by the profile of the abundant G1 stages, which represent 50 % of parasites in an asynchronous population^37^. Therefore, we examined how *Tg*BRG1- and *Tg*BRM-binding patterns correlate with the timing and precision of cell cycle–regulated transcription. *Tg*BRM was significantly more abundant than *Tg*BRG1 at genes associated with shorter cell cycle phases (**Fig. 6A**), such as S, M, and C phases, which represent 28 %, 15 %, and 7 % of asynchronous populations, respectively^37^. A similar pattern was observed for *Tg*MORC (**Extended Data Fig. 10A**). By contrast, *Tg*BRG1 abundance closely matched that of *Tg*BRM at genes expressed during G1a and G1b. However, starting in S phase and continuing through C phase, *Tg*BRM levels were significantly higher than *Tg*BRG1, suggesting a broader or more prolonged association of *Tg*BRM with these loci across the parasite population. This differential enrichment may indicate that *Tg*BRM plays a more sustained role in maintaining chromatin accessibility at these loci, whereas *Tg*BRG1 may be required only transiently (**Fig. 6A–B**).

**Figure 6.**
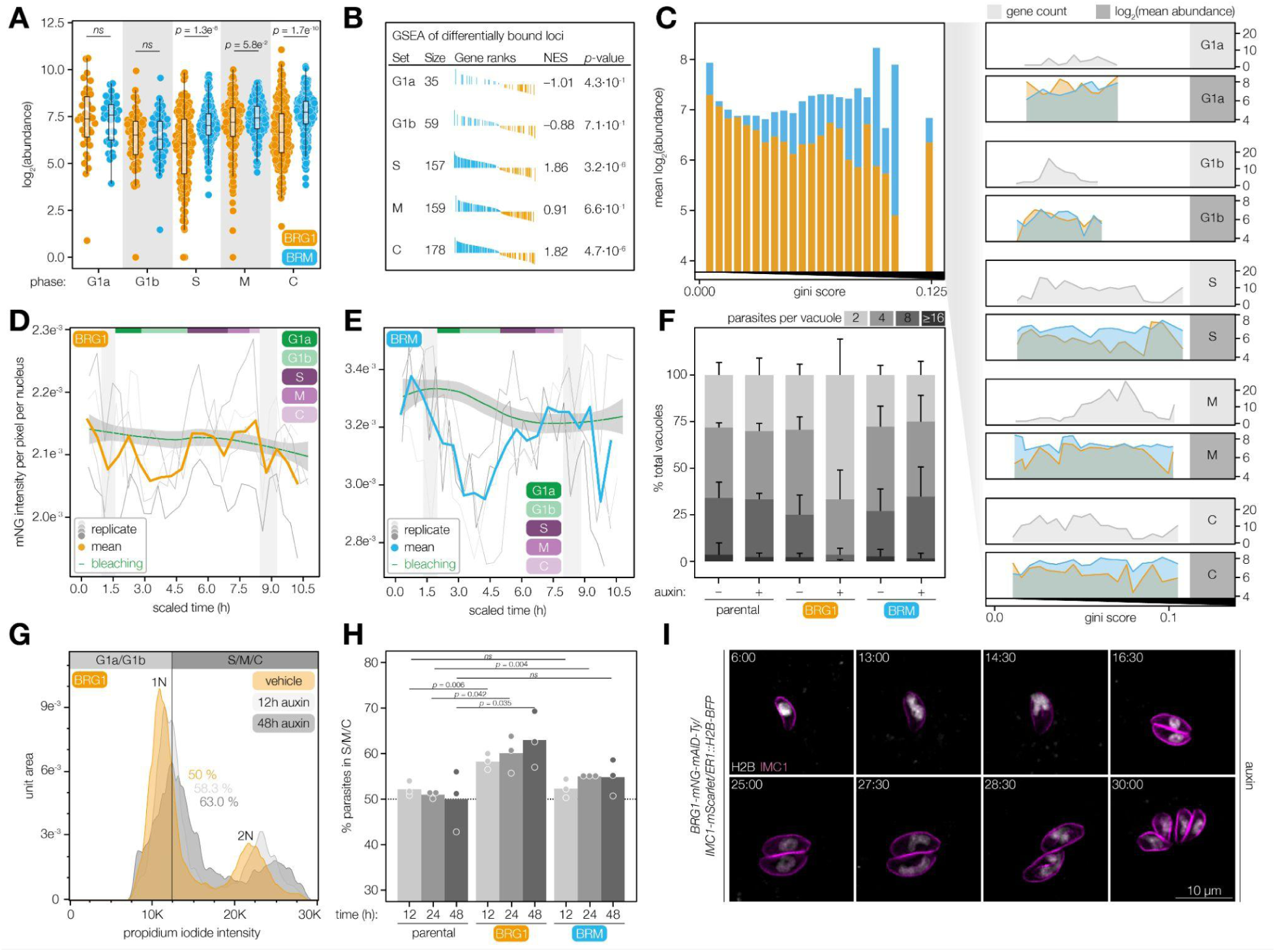
*Tg*BRG1 and *Tg*BRM remodel chromatin with distinct temporal roles during the *Toxoplasma* cell cycle. **A.** *Tg*BRG1 (orange) and *Tg*BRM (blue) abundance at promoters of genes exhibiting phase-specific expression patterns, classified by single-cell RNA-sequencing^37^. Differences between *Tg*BRG1 and *Tg*BRM abundance within each cell cycle phase were assessed using the Wilcoxon rank-sum test. **B.** Gene Set Enrichment Analysis (GSEA) of cell cycle–associated gene sets ranked by the log₂ fold change in occupancy between *Tg*BRG1 and *Tg*BRM at each promoter. Normalized Enrichment Scores (NES) indicate whether *Tg*BRG1 or *Tg*BRM are preferentially associated with genes expressed during specific cell cycle phases. The size of each gene set reflects the number of genes in each gene set^37^ that is also targeted by the SWI/SNF complex while *p*-values were calculated based on an adaptive multi-level split Monte-Carlo scheme^83^. **C.** Mean log₂ abundance of *Tg*BRG1 and *Tg*BRM across genes grouped into Gini score bins (bin size = 0.005), derived from synchronized parasite transcriptomes^39^. A zoom-in panel displays the data split by cell cycle–associated gene sets; the number of genes per bin for each module is shown in light grey plots, and the mean log₂ abundance of *Tg*BRG1 or *Tg*BRM for each module is shown in dark grey plots. **D–E.** Quantification of nuclear mNG fluorescence intensity per pixel from live-cell video microscopy of endogenously tagged *Tg*BRG1 (D) and *Tg*BRM (E). Cell cycle stages are annotated above the plot, with G1a and G1b indicated in shades of green and S, M, and C phases in shades of purple. Grey bars mark periods of ongoing cytokinesis, corresponding to transitions from one-to-two and two-to-four parasites, and account for minor variability in division timing between vacuoles. The green line represents the expected fluorescence decrease due to photobleaching. Fluorescence intensity of *n* = 4 biological replicates was quantified using CellProfiler^84^. **F.** Replication assays of *Tg*BRG1- and *Tg*BRM-mAID–tagged strains and the parental line in the presence or absence of auxin. Parasites per vacuole were quantified from fixed intracellular parasites after 24 hr of growth, based on immunofluorescence images across *n* = 3 biological replicates. Black bars indicate the standard deviation of the mean. **G.** Flow cytometry analysis of DNA content of *Tg*BRG1 knockdown strains 12, 24, and 48 hr after auxin treatment. A representative result of three independent experiments is shown. The solid line shows the gate used to segregate G1 parasites (1N) and replicating (S, M, C: >1–2N) parasites. The indicated percentages represent the mean proportion of parasites with S, M, and C phase DNA content profiles across *n* = 3 biological replicates. **H.** Quantification of DNA content staining of parental as well as *Tg*BRG1 and *Tg*BRM knockdown strains 12, 24, and 48 hr after auxin treatment as shown in (G). The dotted line indicates the expected value for wildtype parasites (50%)^37^. Statistical significance was assessed using Welch’s one-tailed *t-*tests. **I.** Live-cell imaging of *Tg*BRG1 knockdown parasites expressing fluorescent markers for the inner membrane complex (IMC1-mScarlet, magenta) and chromatin (H2B-BFP, white). Parasites were imaged over a 24 hr period beginning 6 hr after auxin-induced knockdown. Selected frames shown from 6 to 30 hr post-knockdown. See also **Video S2**.

The classification of genes to a specific cell cycle phase relies on peak expression; however, it does not consider how much of the total expression is stage specific. To capture the stage specificity of expression, we again used the Gini score where higher values indicate transcripts that are predominantly expressed in one cell cycle phase and likely repressed in others. *Tg*BRM occupancy increases with higher Gini scores, indicating a possible link with genes that remain poised for most of the cell cycle (**Fig. 6C; Extended Data Fig. 10B–C**)^39^. These results reinforce the notion that *Tg*BRG1 and *Tg*BRM are associated with different patterns of gene regulation, and *Tg*BRM is particularly associated with poised or repressed loci not only in preparation for differentiation but also throughout the *Toxoplasma* cell cycle.

The association of *Tg*BRG1 and *Tg*BRM with promoters of cell cycle–regulated genes prompted us to investigate whether their abundance changes across the cell cycle. We measured the intensity of mNG endogenously fused to either *Tg*BRG1 or *Tg*BRM, using live video microscopy. *Tg*BRG1 levels changed modestly throughout the cell cycle. However, *Tg*BRM levels declined following cytokinesis and remained low during G1a and G1b followed by a sharp increase during S phase that coincided with the onset of DNA replication (**Fig. 6D–E; Supplementary Table 13–14**). This S phase–specific surge in *Tg*BRM abundance suggests a role in preserving promoter accessibility during replication-coupled chromatin assembly, in contrast to the stable levels of *Tg*BRG1, which are consistent with the proposed transient role during all transcriptional activation events across the cell cycle.

Knockdown of *Tg*BRG1, but not *Tg*BRM, resulted in a significant reduction in parasite numbers per vacuole, consistent with an immediate slowdown of the parasites cell cycle (**Fig. 6F; Supplementary Table 15**). To determine whether this defect was linked to a specific cell cycle phase, we performed propidium iodide staining to assess DNA content following 12, 24, and 48 hr of knockdown (**Fig. 6G–H; Supplementary Table 16**). *Tg*BRG1-depleted parasites exhibited an increased proportion of cells in S, M, and C phases, a shift not observed after *Tg*BRM knockdown. These results indicate that *Tg*BRG1 is required for timely progression through DNA replication and mitosis. Consistent with this, live-cell imaging revealed prolonged cell cycle phases in *Tg*BRG1-depleted parasites, during which mitosis frequently preceded the formation of daughter cells by ∼1 hr (**Fig. 6I; Video S2**)—a phenotype that was not observed in the vehicle-treated control, where daughter cell budding occurred concurrently with mitosis (**Extended Data Fig. 10D, Video S3**). Together, these findings implicate *Tg*BRG1-dependent chromatin remodeling in the regulation of late cell cycle transitions and the maintenance of proper division timing, highlighting its importance for rapid transcriptional activation during short, transcriptionally dynamic phases of the cell cycle.

### SWI/SNF-dependent genes are defined by their chromatin context

To better understand the transcriptional response to SWI/SNF disruption, we examined the chromatin context of bound promoters and the associated gene bodies. We integrated chromatin profiles of published histone modifications, histone variants, and chromatin-associated factors computing Pearson correlations between the occupancy profiles of *Tg*BRG1, *Tg*BRM, and *Tg*SWI3 and the various chromatin features^7,40,47^. Consistent with our earlier observations of co-occupancy between *Tg*BRM and *Tg*MORC (**Fig. 5H**), *Tg*BRM binding was strongly correlated with binding of *Tg*HDAC3, a known *Tg*MORC-interacting histone deacetylase (**Fig. 7A; Extended Data Fig. 11A**)^7^. By contrast, *Tg*BRG1 was strongly correlated with H4K31ac and H3K14ac, which are typically associated with increased chromatin accessibility^47^. Although *Tg*BRM was also positively correlated with H4K31ac, its presence was negatively correlated with H3K14ac, suggesting divergent interactions with acetylated chromatin. Notably, the active promoter mark H3K4me3 was positively associated with *Tg*BRG1 but negatively correlated with *Tg*BRM. The opposite trend was observed for the repressive H3K9me3 gene body mark^47^, which was positively correlated with *Tg*BRM but negatively with *Tg*BRG1. A weak positive correlation was observed between the SWI/SNF factors and gene body H3.3, a histone variant that marks active and poised promoters. No correlation was found with H2AX, which is associated with gene silencing during the DNA damage response and with silenced promoters activated during chronic stage differentiation (**Fig. 7A; Extended Data Fig. 11A; Supplementary Table 17**)^48–51^. These patterns highlight the distinct chromatin environments associated with *Tg*BRG1 and *Tg*BRM, where *Tg*BRG1 preferentially associates with open chromatin marks such as histone acetylation, while *Tg*BRM is predominantly colocalized with repressive chromatin features such as reduced histone acetylation and enrichment of silencing complexes.

**Figure 7.**
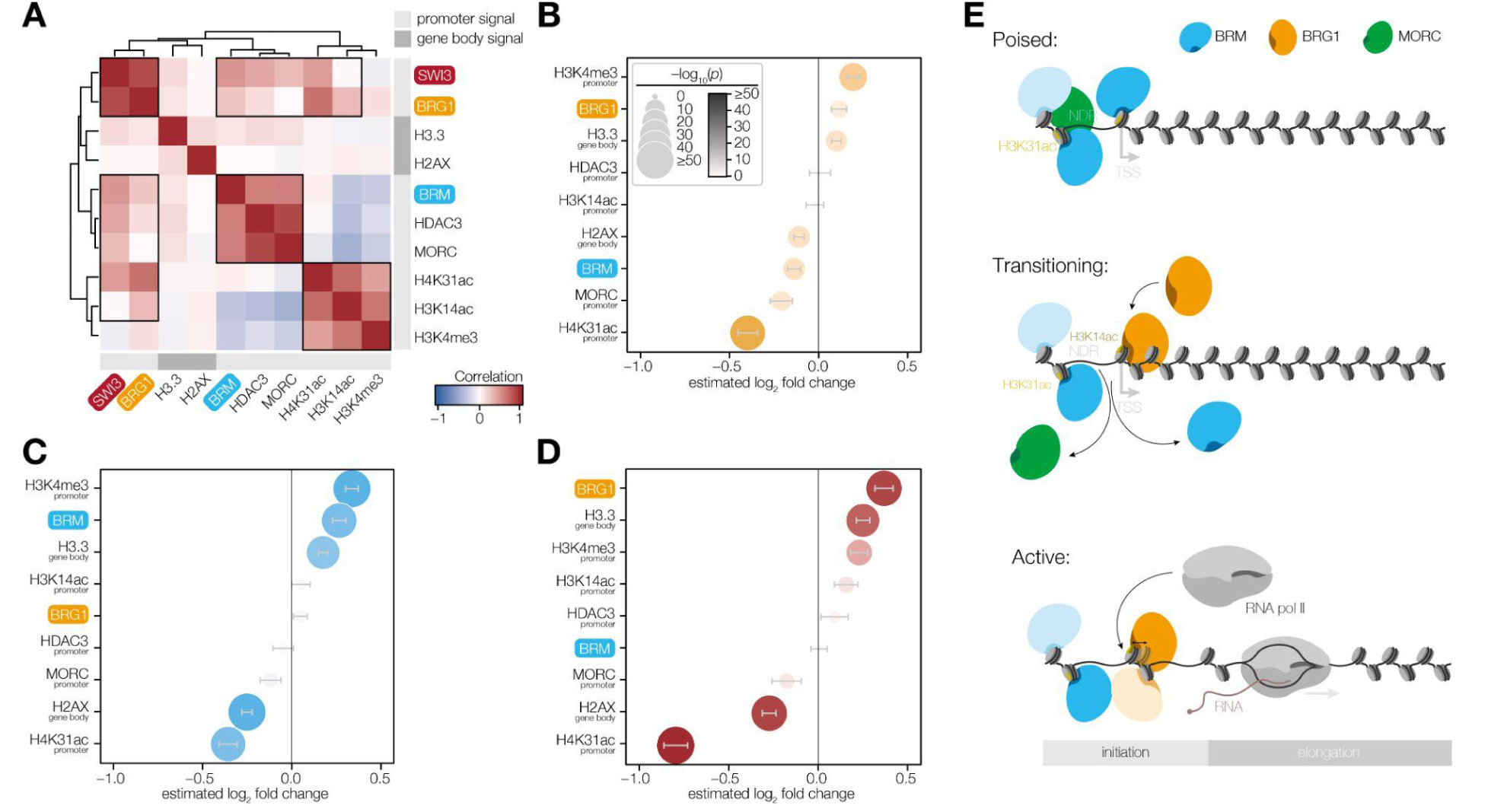
Distinct chromatin environments shape transcriptional responses to SWI/SNF complex disruption. **A.** Correlation heatmap of indicated chromatin-associated features across all genes associated with a consensus locus between *Tg*SWI3, *Tg*BRG1 and *Tg*BRM samples. Pairwise Pearson correlation coefficients were calculated using log_2_ transformed mean-centered values. Light grey bars indicate that the signal was measured over the promoter region within the SWI/SNF consensus locus, whereas dark grey bars denote that the signal was assessed across the gene body. Chromatin-associated features were derived from Farhat et al.^7^, Nardelli et al.^40^ and Sindikubwabo et al.^47^. **B–D.** Multiple linear regression models were fit to assess the contribution of chromatin-associated features to transcriptional changes following *Tg*BRG1 (B), *Tg*BRM (C) or *Tg*SWI3 (D) depletion. Plots show regression coefficients (estimated log_2_ fold change) for each chromatin feature, with 95 % confidence intervals (gray bars). Point size and shade reflect the statistical significance of each predictor, indicating the relative strength of association. **E.** Schematic model illustrating a proposed mechanism of transcriptional regulation mediated by *Tg*BRM and *Tg*BRG1 associated SWI/SNF complexes. Genes poised in a repressed chromatin state are characterized by H3K31ac enrichment, *Tg*BRM, and *Tg*MORC occupancy. Transition to an active state involves loss of *Tg*MORC, acquisition of H3K14ac, and recruitment of *Tg*BRG1. This chromatin remodeling cascade facilitates RNA polymerase II or RNA polymerase III based transcriptional activation.

We turned to multiple linear regression modeling to determine if these chromatin features can explain transcriptional outcomes following SWI/SNF disruption. The modeling estimates the change in gene expression following knockdown of *Tg*BRG1, *Tg*BRM, or *Tg*SWI3 as a function of increasing a given chromatin feature by one standard deviation, while holding all other variables constant. For example, increased abundance of H3K4me3 is associated with up-regulation upon *Tg*BRG1 knockdown, whereas the presence of *Tg*BRG1 was more weakly associated with up-regulation (**Fig. 7B**). By contrast, high levels of the euchromatin mark H4K31ac were predictive of strong down-regulation following *Tg*BRG1 knockdown (**Fig. 7B**). A similar pattern was observed for predictors of the impact of *Tg*BRM knockdown, where high H4K31ac levels were strongly associated with transcriptional repression following knockdown. Repression was additionally associated with elevated H2AX levels within the gene body. Therefore, despite the lack of correlation between *Tg*BRM and H2AX binding, genes highly bound by H2AX are more likely to be downregulated following *Tg*BRM knockdown. This may reflect the role of SWI/SNF complexes in resolving H2AX-associated gene silencing during DNA double-strand break repair^52,53^. Upregulation in the *Tg*BRM knockdown context was again predicted by H3K4me3 and *Tg*BRM itself, potentially reflecting the loss of repressive chromatin features such as *Tg*MORC in the absence of *Tg*BRM (**Fig. 7C**). The effect of *Tg*SWI3 knockdown was best represented by a composite of the predictors for *Tg*BRG1 and *Tg*BRM effects. Repression following *Tg*SWI3 knockdown was most strongly associated with high levels of H4K31ac and H2AX, with the magnitude of the H4K31ac effect closely matching the summed effects of the two helicase knockdowns, suggesting a shared requirement for both remodelers at these loci. Upregulation following *Tg*SWI3 knockdown was correlated with elevated *Tg*BRG1 levels and enrichment of H3.3 in the gene body. (**Fig. 7D**). These results suggest that the complex pattern observed following knockdown of the two SWI/SNF complexes is shaped by the chromatin context of each locus.

## DISCUSSION

Transcriptional regulation coordinates proliferation and developmental transitions in apicomplexan parasites^1–8^. Numerous sequence-specific transcription factors have been identified as critical regulators of cell cycle progression and differentiation^13,18^. However, while these factors have well-defined roles in gene expression control, the mechanisms by which chromatin is restructured for transcriptional activation or repression remain largely unexplored. In this study, we systematically examined the family of Myb domain–containing proteins, since apicomplexans have preferentially retained SANT-type Myb proteins that are commonly found in chromatin remodeling complexes. Using high-throughput tagging^22^ we comprehensively analyzed eight essential factors exhibiting diverse phenotypic and transcriptional effects. We reveal distinct roles during the *Toxoplasma* cell cycle for factors representing three types of ATP-dependent chromatin remodeling complexes: ISWI, INO80, and SWI/SNF. We focused on the SWI/SNF complex, revealing two mutually exclusive SNF2-type ATPase subunits: *Tg*BRG1 and *Tg*BRM. Functional characterization demonstrated that *Tg*BRG1 and *Tg*BRM fulfill distinct yet cooperative regulatory roles. *Tg*BRG1 knockdown preferentially impaired expression of genes active during short cell cycle phases, such as mitosis and cytokinesis, resulting in slowed or stalled progression through these stages. By contrast, *Tg*BRM knockdown induced progressive transcriptional repression and disrupted mitotic coordination, leading to multinucleated parasites and abnormal daughter cell formation. Notably, we also found that *Tg*BRM co-localizes with the repressor *Tg*MORC at poised promoters and contributes to transcriptional priming of genes activated during chronic and sexual development. Together, these findings suggest a division of labor between the SWI/SNF complexes, with *Tg*BRM broadly supporting transcriptional competency, while *Tg*BRG1 mediates transitions in gene expression.

By profiling nuclear-localized Myb proteins required for parasite fitness, we identified five candidates whose knockdown led to widespread transcriptional dysregulation and pronounced cell cycle–related phenotypes. Four of these were predicted to function as chromatin regulators based on domain composition and homology, and one corresponded to the conserved spliceosome-associated factor CDC5L. Knockdown of *Tg*CDC5L resulted in cell cycle arrest in G1. While this manuscript was under preparation a complementary study found that intron retention and the associated formation of protein aggregates likely underlies the cell cycle arrest observed after *Tg*CDC5L knockdown^28^. We focused on chromatin regulators that exhibit varying degrees of conservation among apicomplexans, which included *Tg*SNF2h, a conserved ATPase characteristic of ISWI-family remodelers, and *Tg*DMAP1, an INO80 cofactor. Knockdown of *Tg*SNF2h and *Tg*DMAP1 induced broadly comparable transcriptional responses, characterized by preferential downregulation of G1-expressed genes and modest upregulation of transcripts expressed during S, M, and C phases. Both knockdowns also caused transcriptional de-repression, though with contrasting dependencies on *Tg*MORC. *Tg*DMAP1 depletion led to upregulation of transcripts with high *Tg*MORC occupancy, suggesting that *Tg*DMAP1 may cooperate with *Tg*MORC for gene silencing. In canonical systems, DMAP1 is involved in chromatin-regulatory complexes with opposing functions. As part of a silencing complex, DMAP1 recruits DNMT1 and HDAC2 to newly replicated chromatin, facilitating transcriptional repression via DNA methylation and histone deacetylation^54^. Although *Toxoplasma* was initially thought to lack DNA methylation, later studies confirmed its presence and identified two functional *Tg*DNMTs^55,56^. Conversely, within the INO80 SRCAP complex, DMAP1 contributes to transcriptional activation by promoting histone exchange of canonical H2A–H2B dimers with H2A.Z–H2B dimers^57^. These findings imply that the repressive functions of *Tg*DMAP1 represent the dominant function in tachyzoites.

Genes upregulated upon *Tg*SNF2h depletion were generally marked by lower *Tg*MORC levels. In mammals, SNF2h and its paralog SNF2L maintain transcription factor binding by organizing phased nucleosome arrays immediately adjacent to their binding sites. In *Toxoplasma*, *Tg*SNF2h appears to play an analogous role. It has recently been shown that loss of *Tg*SNF2h triggers widespread chromatin condensation, blocking both *Tg*MORC and transcription factor access at many loci. Yet, its depletion also induces local accessibility shifts that expose new regulatory elements and drive *Tg*MORC-independent gene activation^8^. SNF2h and SNF2L are thought to be primarily recruited via interactions with transcription factors and, to some extent, by specific histone marks^58^. *Tg*SNF2h has similarly been shown to interact with *Tg*AP2VIII-2 and *Tg*RFTS, supporting a conserved recruitment mechanism^8^. Interestingly, in mammalian systems, SNF2h has been found to directly interact with DNMT1 to enhance the deposition of repressive chromatin marks^59^. While this specific interaction may not be conserved in *Toxoplasma*, the similar transcriptional and phenotypic consequences of *Tg*SNF2h and *Tg*DMAP1 knockdown suggest that these factors may exert complementary functions.

In contrast to the widespread transcriptional dysregulation observed following *Tg*DMAP1 or *Tg*SNF2h depletion, *Tg*SWI3 knockdown resulted in comparatively modest changes in gene expression but pronounced morphological defects that included multinucleation and altered chromatin condensation. These distinct phenotypes suggested a unique role for *Tg*SWI3 in coordinating nuclear division and chromatin organization. In animals, plants, and yeast, SWI3 serves as a core structural component of SWI/SNF remodeling complexes, which generally consist of three functional modules: (1) the base module formed by structural components like SWI3 and subunits involved in binding the nucleosome surface, histone tails and DNA through sequence-agnostic contacts; (2) the motor module composed of a SNF2-type ATP-dependent helicase (BRG1; SMARCA4 or BRM; SMARCA2); and (3) the actin-related protein (ARP) module which connects the base module with the motor module^21,31^. Through IP and reciprocal co-IP’s, we identified 17 components of the *Toxoplasma* SWI/SNF complex. These include two mutually exclusive SNF2-type helicases—*Tg*BRG1 and *Tg*BRM—as well as five shared core components present in both complexes. All of the shared components have been found to be fitness-conferring in a genome-wide CRISPR screen, with *Tg*SS18 showing the mildest phenotype score at –2.14^23^. Notably, while both helicases were well conserved across a broad range of representative eukaryotic lineages, many of the remaining complex members showed limited sequence conservation. Among these were canonical core subunits such as SWI3 (SMARCC1/2), SMARCD1 and SMARCB1, which are typically shared between different SWI/SNF subcomplexes or SS18 which is found in specific SWI/SNF subtypes^21,31^. Additional identified proteins include components that may contribute to the ARP module, such as *Tg*ALP2a, or control SWI/SNF stability through ubiquitination, such as RING finger–containing proteins (TGME49_273150 and TGME49_226780)^21^. Substrate recognition may be mediated by DNA-binding proteins such as *Tg*ARID, *Tg*HMGB3 or *Tg*BRG4, a bromodomain-containing protein likely involved in binding acetylated histones^31^. Together, these findings highlight both conserved and parasite-specific components of the *Toxoplasma* SWI/SNF complex and underscore substantial adaptations that may reflect unique regulatory demands imposed by the parasite’s divergent chromatin landscape and complex life cycle^7,15,16^. Additional work will be needed to comprehensively define functional SWI/SNF subcomplexes and to elucidate the specific contributions of novel, apicomplexan-adapted components.

Combining transcriptomics and chromatin profiling, we reveal a division of labor between *Tg*BRG1- and *Tg*BRM-containing complexes, which function collaboratively to regulate gene expression in *Toxoplasma*. Both complexes are widely distributed across the genome and collectively associated with 4,692 loci. Promoters characterized by high levels of both *Tg*BRG1 and *Tg*BRM were associated with high gene expression suggesting that both factors are important for efficient transcription. Transcriptomic analysis revealed that *Tg*BRM knockdown leads to progressive transcriptional repression that mirrors the profile observed upon *Tg*SWI3 depletion, albeit with delayed kinetics for *Tg*BRM. This repression coincided with striking morphological defects: *Tg*BRM depletion disrupted the coordination between mitosis and cytokinesis, resulting in the accumulation of multinucleated parasites and the aberrant formation of multiple daughter cells within a single mother cell. By contrast, *Tg*BRG1 knockdown disproportionately affected genes expressed during short phases of the cell cycle like mitosis and cytokinesis, and led to reduced chromatin condensation, resulting in an immediate slowdown of parasite growth and ultimately parasite death. Although it may seem counterintuitive, genes with a short phase of expression showed greater *Tg*BRM occupancy compared to *Tg*BRG1, which is explained by the probability of finding each factor on its target locus in an asynchronous parasite population, since, based on our model, *Tg*BRM would mark such promoters throughout the cell cycle, while *Tg*BRG1 would only associate during active transcription. Notably, we also identified a subset of loci that were bound by *Tg*BRM and *Tg*MORC but not by *Tg*BRG1; accordingly, these genes were not expressed during the lytic cycle^7^. They were enriched for developmentally regulated genes expressed during parasite differentiation^7,60^. While this pattern has not been appreciated in other systems, these findings suggest distinct temporal roles for *Tg*BRG1 and *Tg*BRM throughout the *Toxoplasma* cell cycle with transcription being more acutely dependent on the presence of *Tg*BRG1 while *Tg*BRM has a more general function in maintaining chromatin in a poised state for rapid activation throughout the cell cycle and developmental transitions^61,62^. Consistent with the role of *Tg*BRM in developmental regulation, chromatin remodeling events involving BRM-associated factors have also been implicated in *Plasmodium* sexual development^63^. Specifically, disruption of *Pf*ARID resulted in a complete loss of male gametocyte formation, despite normal asexual blood-stage replication^63,64^.

The preferential association of *Tg*BRM with genes displaying brief expression windows was less pronounced for M-phase genes. This sustained promoter association may reflect a *Toxoplasma*-specific adaptation of the SWI/SNF-dependent mitotic bookmarking mechanism observed in mammalian systems, whereby subunits SMARCE1 and SMARCB1 remain attached to the chromatin during mitosis to bookmark genes for reactivation while BRG1 is evicted^65^. However, in *Toxoplasma*, mitosis largely overlaps with DNA synthesis and cytokinesis and is accompanied by a single major chromatin remodeling event starting in late S phase and ending in late cytokinesis^6^. This results in a more continuous cell cycle compared to higher eukaryotes and may require *Tg*BRG1 to remain attached to ensure continuous gene expression^36^. This hypothesis is further supported by a study showing that components of both SWI/SNF complexes are differentially phosphorylated in response to *Tg*Crk4 knockdown, a kinase essential for mitotic entry^66^.

We correlated the distinct genome-wide binding patterns of *Tg*BRG1 and *Tg*BRM with published chromatin datasets. This analysis showed that *Tg*BRG1 is enriched at active promoters and correlates with classical markers of euchromatin, such as H3K4me3, H4K31ac, and H3K14ac^47^. Accordingly, *Tg*BRG1 occupancy is elevated at constitutively expressed genes but diminished at genes that are predominantly transcriptionally silent, reinforcing the notion that *Tg*BRG1 associates with genes as they become transcriptionally active. By contrast, *Tg*BRM co-localized with the repressive marks *Tg*MORC and *Tg*HDAC3, but also marks of activation like H4K31ac^7,40,47^. This pattern of chromatin association highlights a dual role for *Tg*BRM in both gene repression and activation. Our regression analysis of factors associated with transcriptional changes following SWI/SNF perturbations revealed that chromatin context, rather than remodeler occupancy alone, best predicts transcriptional sensitivity. Genes marked by high levels of H4K31ac were more likely to be repressed following *Tg*BRG1, *Tg*BRM or *Tg*SWI3 depletion. Conversely, genes upregulated upon *Tg*BRM knockdown were enriched for H3K4me3 and *Tg*BRM occupancy, and likely represent genes maintained in a poised state through a *Tg*BRM-dependent mechanism. Genes upregulated upon *Tg*BRG1 knockdown lacked strong associations with individual chromatin marks. This may indicate that the relevant chromatin features are either heterogeneous or that the correct chromatin context for *Tg*BRG1-dependent de-repression remains unidentified. A compelling candidate is the transcriptional repressor REST, whose DNA binding depends on BRG1 in mammalian systems^67^. While a clear REST ortholog has not been identified in *Toxoplasma*, the presence of structural homologs to key REST cofactors—such as RCOR1—suggests that a functionally analogous repressive complex may exist.

Together, these findings support a model in which *Tg*BRG1 and *Tg*BRM function in a coordinated but non-redundant manner to regulate gene expression (**Fig. 7E**). *Tg*BRM functions primarily to establish and maintain promoter accessibility, acting in concert with repressive factors such as *Tg*MORC and *Tg*HDAC3 to poise chromatin at developmentally regulated or cell cycle-dependent genes. *Tg*BRG1, by contrast, facilitates transcriptional reactivation at these accessible promoters, a function that becomes critical for genes with high temporal specificity, expressed during mitosis and cytokinesis. This division of labor ensures that promoter accessibility is maintained by *Tg*BRM, while *Tg*BRG1 reinstates or sustains transcription, ensuring coordinated gene expression and developmental timing.

## MATERIALS & METHODS

### Plasmids and primers

Oligos were ordered from IDT. All cloning was performed with Q5 2 × master mix (NEB). All oligos used or generated in this study can be found in **Supplementary Table 18**.

### Parasite transfection

*T. gondii* parasites were collected by centrifugation at 1000 × g for 10 minutes and resuspended in Cytomix buffer (10 mM KPO_4_, 120 mM KCl, 0.15 mM CaCl_2_, 5 mM MgCl_2_, 25 mM HEPES, 2 mM EDTA, 2 mM ATP, and 5 mM glutathione). The cell suspension was mixed with transfection constructs to a final volume of 400 µl and electroporation was performed using an ECM 830 Square Wave electroporator (BTX) with 4 mm cuvettes, applying two 1.7 kV pulses for 150 µs each at an interval of 100 ms.

### Strain generation

All Myb domain–containing proteins as well as *Tg*BRG1 and *Tg*BRM were endogenously tagged using a previously described high-throughput tagging (HiT) strategy^22^. Cutting units consisting of homology regions and sgRNA specific to each gene were purchased as gBlocks from IDT and integrated into the HiT vector backbone pGL015 (GenBank: OM640005) or pALH086 (GenBank ON312869) for TGGT1_237520 via Gibson assembly. Vectors were linearized using BsaI-HF V2 (NEB) and co-transfected with the Cas9 expression plasmid pSS014 (GenBank: OM640002) into Type I RH/TIR1/ΔKU80/ΔHXGPRT or Type II ME49/TIR1/ΔKU80/ΔHXGPRT parasites. Clones were selected with 3 µM pyrimethamine or 25 µg ml^−1^ mycophenolic acid and 50 µg ml^−1^ xanthine, isolated via limiting dilution and verified by PCR and immunofluorescence.

*Tg*BRG1, *Tg*BRM and *Tg*BDP1 were endogenously tagged with 2×ALFA^44^ using the HiT vector strategy by introducing gene specific gBlocks into pDS033 empty HiT vector which is based on pGL015 (GenBank: OM640005) but possesses a XTEN-2×ALFA tagging unit. The vector was linearized with BsaI-HF V2 (NEB) and co-transfected with the Cas9 expression plasmid pSS014 (GenBank: OM640002) into Type I RH/ΔKU80/ΔHXGPRT parasites. Clones were selected with 3 µM pyrimethamine.

Cell cycle reporter strains expressing IMC1 endogenously tagged with mScarlet-I^68^ as well as H2B tagged with BFP^69^ ectopically expressed from empty region 1 (ER1, Chr. IV)^70^ were generated by co-transfecting the following constructs into Type I RH/TIR1/ΔKU80/ΔHXGPRT: For IMC1-mScarlet-I, a sgRNA targeting the IMC1 locus was introduced into the Cas9 expression plasmid pSS013 (GenBank: [OM640003](https://www.ncbi.nlm.nih.gov/nuccore/2233454012)) which was co-transfected with an XTEN-mScarlet-I amplicon generated from pDS131 empty HiT vector which is based on pALH193^71^ but possesses mScarlet-I derived from Addgene plasmid #125138 instead of mNG. The amplicon was generated using primers containing IMC1 homology arms. A copy of H2B-BFP was introduced to ER1 by co-transfection of pBM041 (GeneBank: MN019116) with an H2B-BFP amplicon containing the *Tg*Tub1 5’UTR as well as the *Tg*DHFR 3’UTR sequences and ER1 homology arms generated from pDS118 which was derived from pH2B-YFP^23^ by replacing the YFP sequence with the sequence of BFP derived from Addgene plasmid #171090. Clones were selected by flow cytometric sorting of double-positive parasites expressing both mScarlet-I and BFP.

*Tg*BDP1 endogenously tagged at the C-terminus using the HiT strategy was additionally tagged at the N-terminus with a 2×ALFA epitope tag by co-transfecting pSS013 containing a sgRNA targeting *Tg*BDP1 at the N-terminus as well as a 2×ALFA-XTEN amplicon generated with primers containing *Tg*BDP1 homology arms. Clones were selected by limiting dilution.

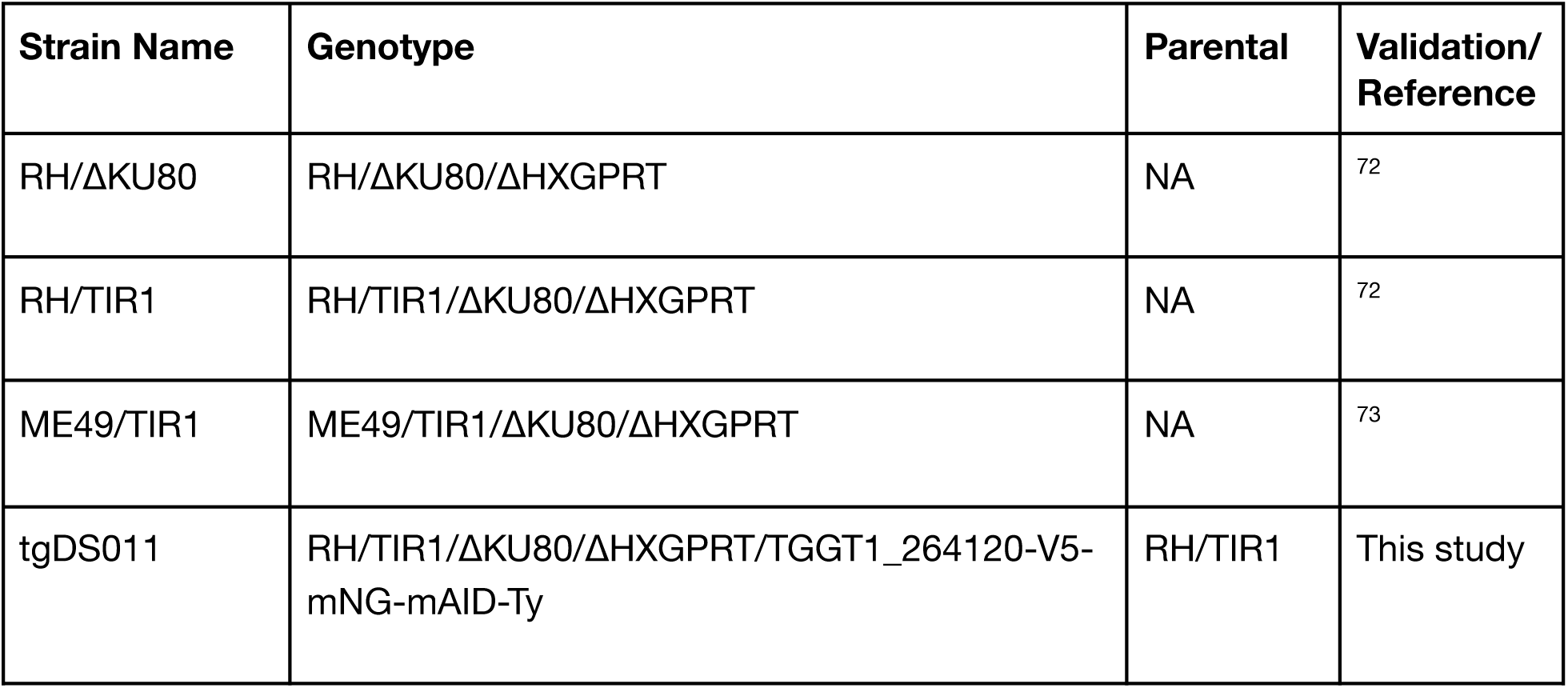
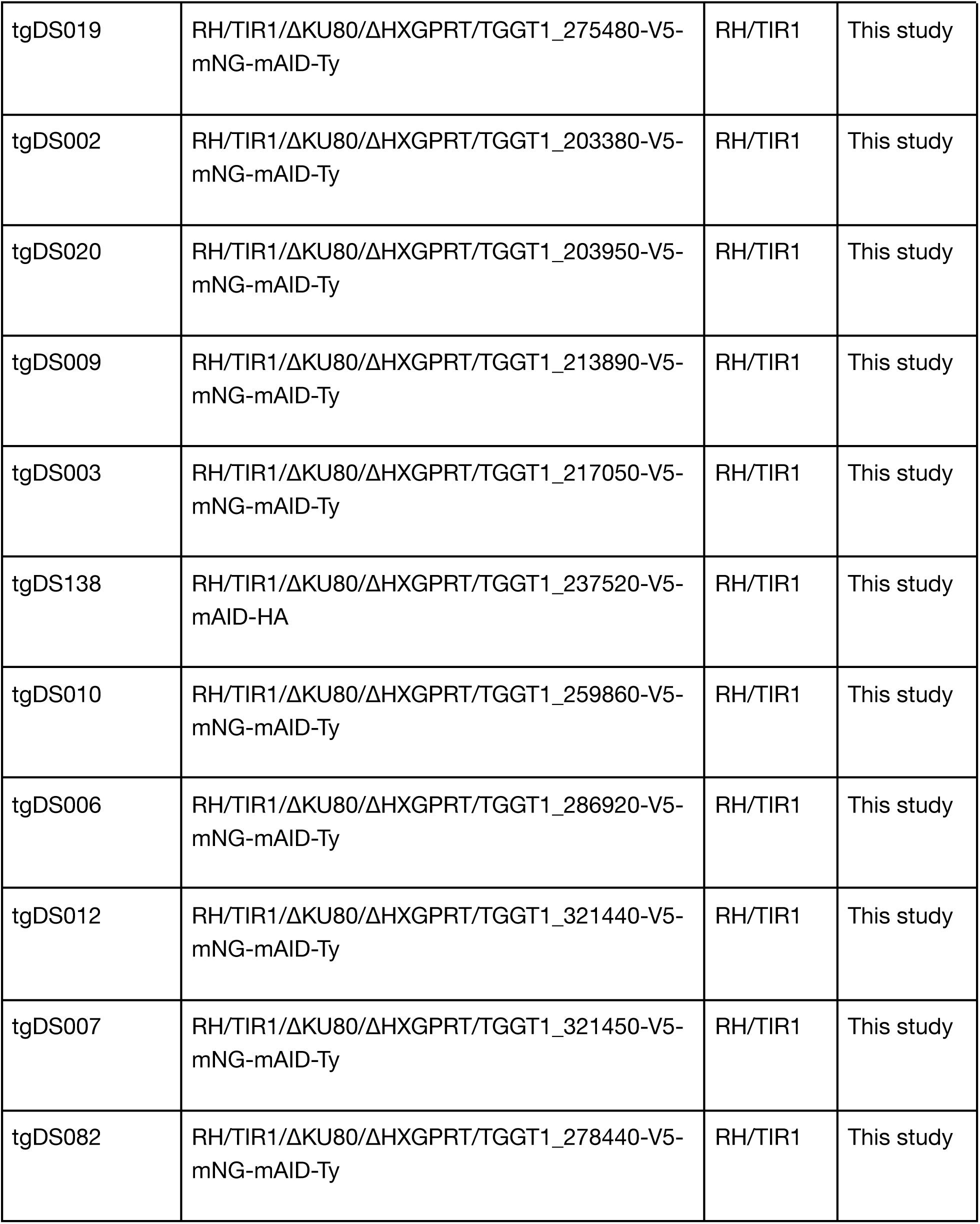
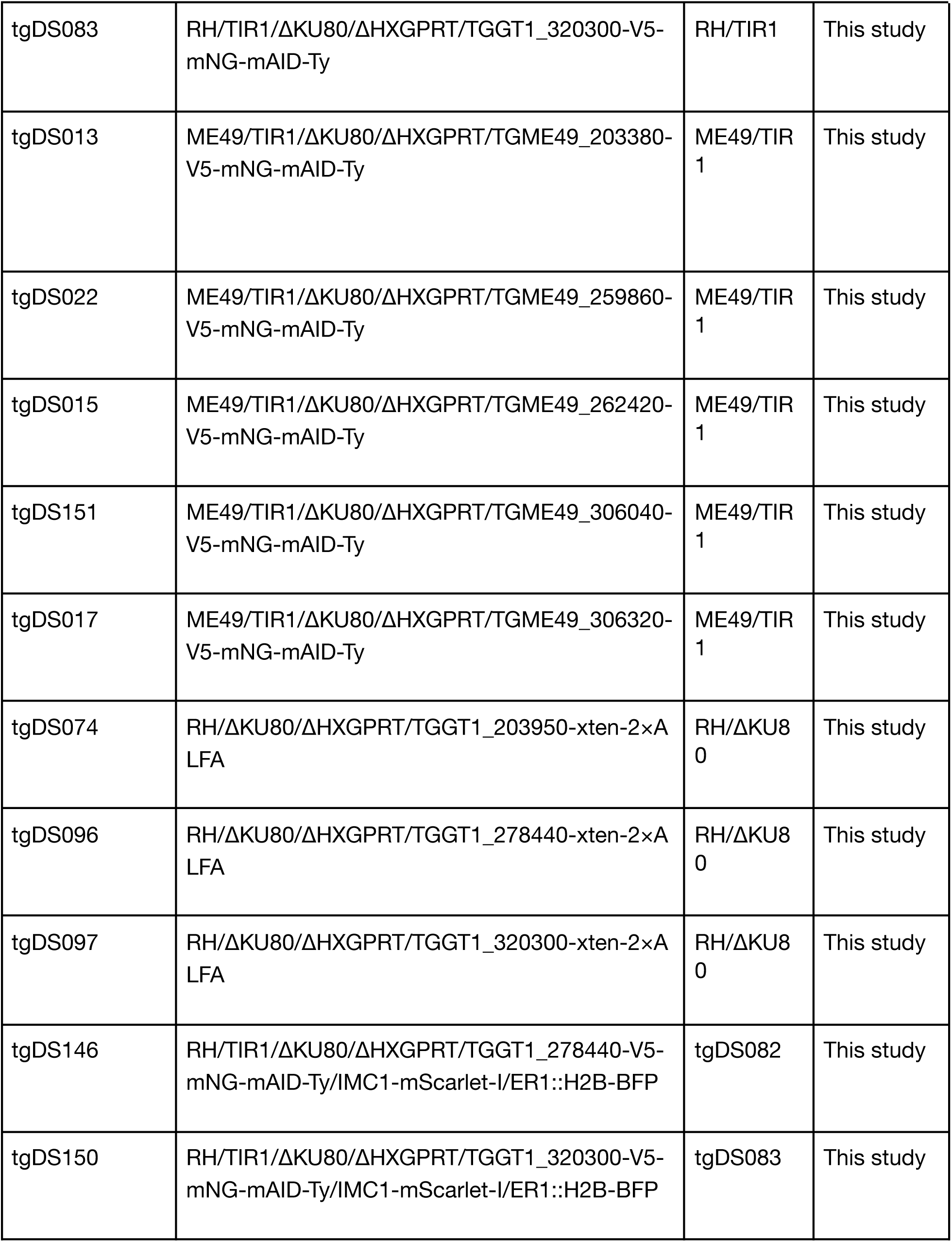
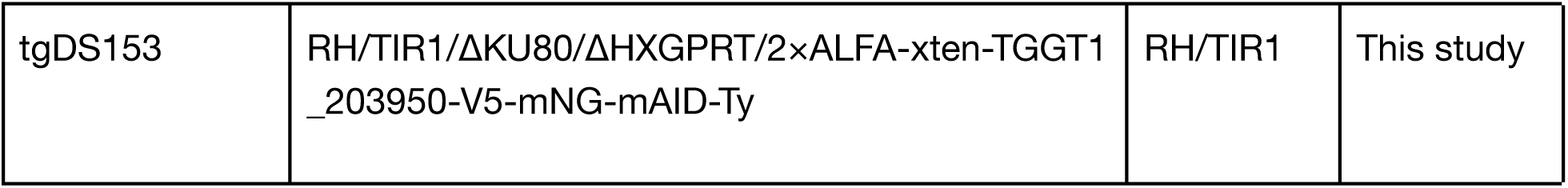

| Strain Name | Genotype | Parental | Validation/<br>Reference |
| --- | --- | --- | --- |
| RH/ΔKU80 | RH/ΔKU80/ΔHXGPRT | NA | <sup>72</sup> |
| RH/TIR1 | RH/TIR1/ΔKU80/ΔHXGPRT | NA | <sup>72</sup> |
| ME49/TIR1 | ME49/TIR1/ΔKU80/ΔHXGPRT | NA | <sup>73</sup> |
| tgDS011 | RH/TIR1/ΔKU80/ΔHXGPRT/TGGT1_264120-V5-mNG-mAID-Ty | RH/TIR1 | This study |
| tgDS019 | RH/TIR1/ΔKU80/ΔHXGPRT/TGGT1_275480-V5-mNG-mAID-Ty | RH/TIR1 | This study |
| tgDS002 | RH/TIR1/ΔKU80/ΔHXGPRT/TGGT1_203380-V5-mNG-mAID-Ty | RH/TIR1 | This study |
| tgDS020 | RH/TIR1/ΔKU80/ΔHXGPRT/TGGT1_203950-V5-mNG-mAID-Ty | RH/TIR1 | This study |
| tgDS009 | RH/TIR1/ΔKU80/ΔHXGPRT/TGGT1_213890-V5-mNG-mAID-Ty | RH/TIR1 | This study |
| tgDS003 | RH/TIR1/ΔKU80/ΔHXGPRT/TGGT1_217050-V5-mNG-mAID-Ty | RH/TIR1 | This study |
| tgDS138 | RH/TIR1/ΔKU80/ΔHXGPRT/TGGT1_237520-V5-mAID-HA | RH/TIR1 | This study |
| tgDS010 | RH/TIR1/ΔKU80/ΔHXGPRT/TGGT1_259860-V5-mNG-mAID-Ty | RH/TIR1 | This study |
| tgDS006 | RH/TIR1/ΔKU80/ΔHXGPRT/TGGT1_286920-V5-mNG-mAID-Ty | RH/TIR1 | This study |
| tgDS012 | RH/TIR1/ΔKU80/ΔHXGPRT/TGGT1_321440-V5-mNG-mAID-Ty | RH/TIR1 | This study |
| tgDS007 | RH/TIR1/ΔKU80/ΔHXGPRT/TGGT1_321450-V5-mNG-mAID-Ty | RH/TIR1 | This study |
| tgDS082 | RH/TIR1/ΔKU80/ΔHXGPRT/TGGT1_278440-V5-mNG-mAID-Ty | RH/TIR1 | This study |
| tgDS083 | RH/TIR1/ΔKU80/ΔHXGPRT/TGGT1_320300-V5-mNG-mAID-Ty | RH/TIR1 | This study |
| tgDS013 | ME49/TIR1/ΔKU80/ΔHXGPRT/TGME49_203380-V5-mNG-mAID-Ty | ME49/TIR1 | This study |
| tgDS022 | ME49/TIR1/ΔKU80/ΔHXGPRT/TGME49_259860-V5-mNG-mAID-Ty | ME49/TIR1 | This study |
| tgDS015 | ME49/TIR1/ΔKU80/ΔHXGPRT/TGME49_262420-V5-mNG-mAID-Ty | ME49/TIR1 | This study |
| tgDS151 | ME49/TIR1/ΔKU80/ΔHXGPRT/TGME49_306040-V5-mNG-mAID-Ty | ME49/TIR1 | This study |
| tgDS017 | ME49/TIR1/ΔKU80/ΔHXGPRT/TGME49_306320-V5-mNG-mAID-Ty | ME49/TIR1 | This study |
| tgDS074 | RH/ΔKU80/ΔHXGPRT/TGGT1_203950-xten-2xA LFA | RH/ΔKU80 | This study |
| tgDS096 | RH/ΔKU80/ΔHXGPRT/TGGT1_278440-xten-2xA LFA | RH/ΔKU80 | This study |
| tgDS097 | RH/ΔKU80/ΔHXGPRT/TGGT1_320300-xten-2xA LFA | RH/ΔKU80 | This study |
| tgDS146 | RH/TIR1/ΔKU80/ΔHXGPRT/TGGT1_278440-V5-mNG-mAID-Ty/IMC1-mScarlet-I/ER1::H2B-BFP | tgDS082 | This study |
| tgDS150 | RH/TIR1/ΔKU80/ΔHXGPRT/TGGT1_320300-V5-mNG-mAID-Ty/IMC1-mScarlet-I/ER1::H2B-BFP | tgDS083 | This study |
| tgDS153 | RH/TIR1/ΔKU80/ΔHXGPRT/2xALFA-xten-TGGT1_203950-V5-mNG-mAID-Ty | RH/TIR1 | This study |

### Cell culture

*Toxoplasma gondii* parasites and human foreskin fibroblasts (HFFs, ATCC SRC-1041) were cultured in Dulbecco’s Modified Eagle Medium (DMEM, GIBCO 11965118) supplemented 10 μg/ml gentamicin (Thermo Fisher Scientific), 2 mM L-glutamine (Gemini Bioproducts) and 3 % inactivated calf serum (ICS) for RH parasites or 3 % inactivated fetal bovine serum (IFS) for ME49 parasites. Tachyzoites and HFFs were grown at 37 °C and 5 % CO_2_. Stress conditions were induced using alkaline stress medium consisting of RPMI 1640 (Sigma) supplemented with 1 % IFS, 10 μg/ml gentamicin, and buffered with 50 mM HEPES, adjusted to pH 8.1 using 10N NaOH. HFFs and parasites were routinely tested for mycoplasma using the ATCC Universal Mycoplasma Detection Kit (30-1012K).

### Immunohistochemistry

HFFs were grown on coverslips for 2–6 days in DMEM (GIBCO 11965118) supplemented with 10 μg/ml gentamicin (Thermo Fisher Scientific), 2 mM L-glutamine (Gemini Bioproducts) and 10 % IFS to form a confluent monolayer before infection with *Toxoplasma*. Intracellular parasites were fixed for 20 min in ice-cold 4 % formaldehyde in PBS, permeabilized with 1 % Triton X-100 in PBS for 8 min, and blocked for 10 min in PBS with 5 % normal goat serum and 5 % IFS. Primary and secondary antibody incubations were performed for 1 hr each, with three intermediate PBS washes after each step. Before mounting, coverslips were washed by a brief immersion in dH₂O and mounted on Prolong Diamond (Thermo Fisher Scientific) at 37 °C for 30 min or overnight at room temperature. GAP45 was detected using the rabbit anti-GAP45 antibody (1:5,000 dilution) which was a gift from Dominique Soldati (University of Geneva)^74^. Mouse anti-mNG (ChromoTek) was used at 1:1,000. Mouse anti-IMC1 was a gift from Gary Ward (University of Vermont)^75^ and used at 1:2,000. Guinea pig anti-CDPK1 (Covance) was used at 1:10,000. Rabbit anti-Centrin1 (Kernfast) was used at 1:500. The ALFA tag was detected using either FluoTag-X2 anti-ALFA-647 (NanoTag Biotechnologies) at 1:1,000 or rabbit anti-ALFA (NanoTag Biotechnologies) at 1:1,000. Cyst walls were stained with *Dolichos biflorus* lectin (DBL-488 or DBL-575; 1:500 or 1:200, VectorLabs). Secondary antibodies conjugated to Alexa Fluor 488, 594 or 647 (Thermo Fisher Scientific) were used at 1:1,000, and DNA was counterstained with Hoechst 33258 (1:20,000, Life Technologies). Images were acquired using an Eclipse Ti-E epifluorescence microscope or a Yokogawa Spinning Disk Field Scanning Confocal microscope and processed in FIJI^76^.

### Differentiation assay

Confluent HFF monolayers grown on coverslips were infected with parasites and cultured in DMEM (GIBCO 11965118) supplemented with 10 μg/ml gentamicin (Thermo Fisher Scientific), 2 mM L-glutamine (Gemini Bioproducts) and 3 % IFS. At 24 hr post-infection, the media was exchanged for fresh media containing either 500 µM auxin (indole-3-acetic acid) or vehicle (PBS). After 4 hr of auxin or vehicle treatment, parasites were shifted to stress conditions (alkaline stress medium) supplemented with 500 µM auxin or vehicle. Stressed parasites were cultured at 37 °C and ambient CO_2_ for 48 hr before cells were fixed and stained as described under **Immunohistochemistry**. Parasites were probed with antibodies against mNG as well as DBL-575 and Hoechst dye to label the parasite’s cyst wall or DNA, respectively. DBL staining was quantified for ≥100 vacuoles per replicate.

### Replication assay

Confluent HFF monolayers grown on coverslips were infected with parasites and cultured in DMEM (GIBCO 11965118) supplemented with 10 μg/ml gentamicin (Thermo Fisher Scientific), 2 mM L-glutamine (Gemini Bioproducts), 3 % ICS and 50 µM auxin or vehicle (PBS), respectively. At 24 hr post-infection, cells were fixed and stained as described under **Immunohistochemistry**. Parasites were probed with antibodies against GAP45 and mNG as well as Hoechst dye to label DNA. Parasites per vacuole were quantified for ≥100 vacuoles per replicate.

### Video microscopy

Confluent HFF monolayers were grown in glass-bottom imaging dishes (VWR International) and infected with parasites ∼10 hr before imaging and grown in FluoroBrite™ DMEM (Thermo Fisher Scientific) supplemented with 10 μg/ml gentamicin (Thermo Fisher Scientific), 2 mM L-glutamine (Gemini Bioproducts), 10 % IFS and 50 µM auxin or vehicle (PBS), respectively. Parasites were imaged at 37 °C and 5 % CO_2_ for 24 hr at 1 frame every 20 min using a Dual-Fusion BT camera or at 1 frame every 30 min using a Fusion BT camera. Videos were cropped and annotated using NIS elements (Nikon) and FIJI^76^.

### Flow cytometry analysis

Parasite DNA content following *Tg*BRG1 and *Tg*BRM knockdown was evaluated by flow cytometry using a modified propidium iodide (PI, Sigma Aldrich) staining protocol described previously^66^. HFF monolayers were infected with parasites and cultured for 24 hr at 37 °C with 5 % CO₂ before they were treated with 50 µM auxin or vehicle (PBS) for 12, 24, or 48 hr. Following treatment, monolayers were aspirated, rinsed with PBS, and parasites were mechanically released by scraping and passage through a 27-gauge needle, followed by filtration through a 5 μm membrane. Parasite suspensions were centrifuged at 1,500 × g for 10 min and fixed by resuspension of the pellet in 70 % ice-cold ethanol. Samples were stored at –20 °C until flow cytometry analysis. On the day of analysis, parasites were pelleted at 4 °C at 800 × g for 5 min, the supernatant was aspirated, and pellets were resuspended in 1 ml ice-cold PBS. PI was added to a final concentration of 0.1 mg/ml, along with RNaseA/T1 mix (Thermo Fisher Scientific) to 0.08 mg/ml. Samples were incubated for 30 min in the dark at room temperature. DNA content was measured by flow cytometry using a MACSQuant VYB Flow Cytometer (Miltenyi Biotec). Flow cytometry data were analyzed using FlowJo (v10.10.0).

### Plaque assay

Plaque assays were performed by infecting a 6-well plate containing a confluent monolayer of HFFs with 500 parasites per well. Parasites were cultured in DMEM (GIBCO 11965118) supplemented 10 μg/ml gentamicin (Thermo Fisher Scientific), 2 mM L-glutamine (Gemini Bioproducts), 10 % IFS and 50 µM auxin or vehicle (PBS), respectively. Cultures were maintained undisturbed for 10 days, then washed with PBS and fixed in 100 % ethanol for 10 min at room temperature. Plates were air-dried, stained with crystal violet (12.5 g crystal violet, 125 ml 100 % ethanol and 500 ml 1 % ammonium oxalate) for 1 hr, washed three times with PBS and once with water, and air-dried prior to imaging.

### RNA-sequencing

Parasites were seeded onto confluent HFF monolayers in 10 cm dishes and allowed to replicate for 24 hr in DMEM (GIBCO 11965118) supplemented with 10 μg/ml gentamicin (Thermo Fisher Scientific), 2 mM L-glutamine (Gemini Bioproducts), 3 % ICS. Thereafter, the media was replaced with DMEM (GIBCO 11965118) supplemented with 10 μg/ml gentamicin (Thermo Fisher Scientific), 2 mM L-glutamine (Gemini Bioproducts), 3 % ICS and 50 µM auxin or vehicle (PBS), respectively. Parasites were cultured for an additional 6, 12, or 24 hr before harvest. Monolayers were aspirated, rinsed with PBS, and parasites mechanically released by scraping and passage through a 27-gauge needle, followed by filtration through a 5 μm

membrane. Parasite suspensions were centrifuged at 1,500 × g for 5 min, and RNA was extracted from parasite pellets by double extraction using TRIzol (Invitrogen). Therefore, the pellet was resuspended in 800 µl TRIzol Reagent (Invitrogen), vortexed, and split into two different tubes to retain one technical replicate. Phase separation was performed in phase lock heavy tubes (Quantabio) by addition of 100 μl chloroform:isoamyl alcohol (24:1, Sigma-Aldrich), vigorous vortexing (1 min), and centrifugation at 16,000 × g for 5 min at 4 °C. The aqueous phase was isolated, and TRIzol extracted again using 400 µl TRIzol. The resulting aqueous phase was supplemented with 1 μl Glycogen (20 mg/ml, Thermo Fisher Scientific) and 500 μl isopropanol, and RNA was precipitated at room temperature for 12 min. Pellets were recovered by centrifugation (12,000 × g, 12 min) at 4 °C, washed twice with 70 % ethanol, air-dried for 10 min at room temperature, and resuspended in 20 μl RNase-free dH_2_O. RNA quality was assessed using a Bioanalyzer (Agilent).

RNA-seq libraries were prepared with the Rapid Directional RNA-Seq Kit 2.0 (NEXTFLEX) using Poly(A) Beads 2.0 (NEXTFLEX). Paired-end sequencing (50/50 or 100/100) was performed on an Illumina NovaSeq SP platform. Reads were trimmed using Trim Galore (v0.6.7) and aligned to the Toxoplasma gondii ME49 reference genome (ToxoDB v68) with STAR (v2.7.1a) using the --quantMode GeneCounts option and the parameters --outFilterMismatchNoverLmax 0.04 and --outFilterMatchNminOverLread 0.4 to control alignment stringency. Differential expression analysis was performed using the DESeq2 R package (v. 1.40.2)^77^. Genes were classified as up- or down regulated using the Benjamini-Hochberg adjusted *p*-value. Genes with an adjusted *p*-value < 0.005 and a fold change > 2 were defined as upregulated, while genes with an adjusted *p*-value < 0.005 and a fold change < –2 were defined as downregulated. To minimize noise from low-abundance transcripts, only genes with a normalized mean read count greater than 17.3 or 37.9 were considered differentially regulated; genes below this threshold were excluded from subsequent analyses based on their expression distribution. Replicate 1 of the TGME49_275480 knockdown was excluded from further analysis due to underrepresentation in the sequencing libraries (**Extended Data Fig. 2A**).

### Immunoprecipitation

For anti-mNG immunoprecipitations, anti-mNG-Trap magnetic agarose resin (ChromoTek) was used while for anti-ALFA immunoprecipitations, anti-ALFA Selector PE magnetic agarose resin (NanoTag Biotechnologies) was used. Beads were pre-eluted using 50 mM HEPES-NaOH (pH 7.5), 0.4 % IGEPAL and 2 mM Dithiothreitol (DTT) in dH_2_O. Nuclear lysates of approximately 2.5×10^7^ intracellular parasites were generated using the NE-PER Nuclear and Cytoplasmic Extraction Kit (Thermo Fisher Scientific) according to the manufacturer’s protocol. The nuclear fraction was added to the pre-eluted beads and incubated at room temperature for 30 min at rotation followed by three washes with 50 mM HEPES-NaOH (pH 7.5), 0.04 % IGEPAL and 2 mM DTT in dH_2_O. Elution of bound proteins from beads was performed by incubation of beads in elution buffer (5 % SDS, 40 mM chloroacetamide, 10 mM Tris(2-carboxyethyl)phosphine) and 100 mM Triethylammonium bicarbonate (TEAB) in dH_2_O) for 10 min at 70 °C and slow mixing. The elution was transferred to a new tube, and the elution was repeated. Elutions were pooled, and the standard S-trap micro workup protocol (Profiti) was used to prepare samples for LC-MS/MS collection.

In brief, samples were acidified with 2.5 μl 27.5 % phosphoric acid and precipitated by adding 165 μl S-trap binding buffer (100 mM TEAB in 90 % ice-cold methanol). The total volume was applied to S-trap Micro columns and the precipitated proteins were separated from suspension via centrifugation for 1 min at 4000×g. Detergents and salts were removed by washing the S-trap columns six times with 150 μl S-trap binding buffer. Proteins caught by the S-trap fiber glass filters were enzymatically digested by adding 1 μg trypsin/LysC mix dissolved in 30 μl 100 mM TEAB and incubation overnight at 37 °C in a humidified incubator. Tryptic peptides were collected by centrifuging for 1 min at 4000×g followed by sequential washes with 40 μl of 50 mM TEAB pH 8.5, 40 μl 0.2 % formic acid in water, and 40 μl 50 % acetonitrile in water. The resulting peptides were pooled, lyophilized, and dissolved in 12 μl 0.2 % formic acid in MS-grade water.

LC-MS/MS data was generated using a Easy-nLC 1200 system coupled with an Orbitrap Exploris 480 mass spectrometer, both from Thermo Fisher Scientific (Waltham, MA, USA). The setup included a FAIMS Pro Interface and an Easy Spray ESI source. NanoLC separation utilized an Acclaim PepMap trap column (75 μm × 2 cm) (Thermo Fisher Scientific) combined with an Aurora Ultimate TS25 column (75 µm × 25 cm, 120 Å) from IonOpticks (Fitzroy, VIC, AUS). A 5 µl volume of peptide extracts was injected. Peptide separation was conducted with a mobile phase consisting of 0.1 % formic acid in water (solution A) and 0.1 % formic acid in 80 % acetonitrile (solution B), flowing at 400 nl/min, while the column temperature was held constant at 50 °C. Peptides were separated on a gradient of 3–25 % solution B for 68 min, 25–40 % solution B for 22 min, 40–95 % solution B for 3 min and 95 % solution B for 7 min. Using the MS in positive mode, the ion source temperature was set to 270 °C, and ionized peptides were passed through the FAIMS Pro at -45 and -65 V. Mass spectra were collected in MS1 mode with a resolution of 60,000, spanning the mass range of 350–1400 m/z using custom automatic gain control (AGC = 300) settings and an injection time of 25 ms. For MS2 data acquisition, the MS was operated in DDA mode with dynamic exclusion and a resolution of 15,000. The top precursors with charge states 2–6 were selected for MS2 analysis using a dynamic exclusion window of 45 sec. The following parameters were used for MS2 acquisition: automatic maximum injection time, standard AGC target, isolation window of 1.3 m/z, 30 % normalized collision energy, and an intensity threshold of 5 × 10^3^. Data analysis was performed using Proteome Discoverer (v2.4.1.15) for peptide identification and quantification. Downstream data processing was conducted using a custom R pipeline incorporating the packages dplyr (v1.1.4) and readxl (v1.4.3).

### CUT&RUN

#### NbALFA-MNase generation

The His-tagged NbALFA-MNase fusion protein construct was generated from the original pAG-MNase fusion protein construct generously provided by Mary Gehring^43^. The original vector was digested with AfeI (NEB) and mluI-HF (NEB) to replace the endogenous pAG sequence with the NbALFA sequence derived from the original ALFA-tag publication^44^. The DNA plasmid was transformed into chemically competent JM101 cells (Agilent) according to the manufacturer’s instructions and cultured in NZYM medium supplemented with 50 μg/ml kanamycin. At an OD₆₀₀ of ∼0.6, NbALFA-MNase expression was induced with 2 mM isopropyl-β-D-1-thiogalactopyranoside (IPTG; Thermo Fisher Scientific) for 2 hr at 37 °C. Cells were harvested by centrifugation (3,500 × g, 20 min), washed in ice-cold PBS, and lysed by resuspension in 10 ml lysis buffer (10 mM Tris-HCl pH 7.5, 300 mM NaCl, 10 mM imidazole, 5 mM β-mercaptoethanol) containing 10 μg/ml lysozyme (Sigma-Aldrich). After 15 min at 30 °C, cells were lysed using a microfluidizer at 4 °C. Lysates were clarified by centrifugation (12,000 × g, 20 min, 4 °C) and loaded onto a Ni-NTA agarose column (Qiagen). Binding was performed for 1 hr at 4 °C on a rotator, and columns were washed with wash buffer (10 mM Tris-HCl pH 7.5, 300 mM NaCl, 20 mM imidazole, 0.03 % Zwittergent 3-10). The NbALFA-MNase fusion protein was eluted with elution buffer (10 mM Tris-HCl pH 7.5, 300 mM NaCl, 250 mM imidazole) and dialyzed twice at 4 °C (2 hr and overnight) against dialysis buffer (10 mM Tris-HCl pH 7.5, 150 mM NaCl, 1 mM EDTA, 1 mM PMSF) using Slide-A-Lyzer cassettes (Thermo Fisher Scientific). Purified protein was supplemented with glycerol to 50 % final concentration and stored at –80 °C. The quality of the NbALFA-MNase purification was evaluated by Coomassie Brilliant Blue staining, and enzymatic activity was assessed by digestion of high-molecular-weight lambda DNA (NEB).

##### ALFA-CUT&RUN

Intracellular parasites were harvested by mechanical disruption through a 27-gauge needle and filtration through a 5 μm membrane. A total of ∼5 × 10⁷ parasites were pelleted (1500 × g, 10 min) and resuspended in wash buffer (20 mM HEPES-NaOH pH 7.5, 150 mM NaCl, 0.5 mM Spermidine, 1x EDTA-free Protease Inhibitor Cocktail) and attached to 100 μl BioMag Plus Concanavalin A beads (Polysciences) activated in bead activation buffer (20 mM HEPES-KOH pH 7.9, 10 mM KCl, 0.1 mM CaCl_2_, 0.1 mM MnCl_2_). Parasites were bound to the beads by incubation for 15 min at room temperature and rotation. Bead-bound parasites were blocked with wash buffer supplemented with 2 mM EDTA for 5 min at room temperature followed by binding of 1 µg NbALFA-MNase per sample in ice-cold wash buffer supplemented with 0.025 % Digitonin (Invitrogen) for 1 hr at 4 °C. After NbALFA-MNase binding, bead-bound parasites were washed three times with ice-cold wash buffer supplemented with 0.025 % Digitonin for 5 min at 4 °C. The cutting reaction was started by the addition of ice-cold wash buffer supplemented with 0.025 % Digitonin and 2 mM CaCl_2_ and incubation for 60 min on ice. The reaction was stopped by the addition of 4×STOP buffer (400 mM NaCl, 40 mM EDTA, 8 mM EGTA, 100 µg/ml RNaseA, 80 µg/ml Glycogen and 0.025 % Digitonin) to 1× and incubation at 37 °C for 20 min. The DNA was extracted using a modified organic extraction and ethanol precipitation protocol described previously^3^.

#### CUT&RUN library preparation

CUT&RUN libraries were prepared using the NEBNext Ultra II DNA Library Prep Kit for Illumina (NEB) and NEBNext Multiplex Oligos for Illumina (NEB) according to the manufacturer’s instructions with 11–12 PCR amplification cycles. Size selection and clean-up steps were performed with AMPure XP beads (Beckman Coulter). Library quality and fragment size distribution was assessed using a Bioanalyzer (Agilent). Paired-end sequencing (75/75) was performed on an Illumina MiSeq v3 platform, generating approximately 6 million reads per sample.

##### CUT&RUN data processing and analysis

Paired-end reads were trimmed using Trim Galore (v0.6.7) with a Phred quality cutoff of 20 and discarding reads shorter than 30 bp. Trimmed reads were aligned to the *Toxoplasma gondii* ME49 reference genome (ToxoDB v68) using Bowtie2 (v2.4.2) with the parameters: --end-to-end -D 15 -R 2 -N 1 -L 22 -i S,1,0.50 --no-mixed --no-discordant --phred33 -I 10 -X 700. Peak calling was performed with MACS2 (v2.2.7.1) using the following options: -f BAMPE -g 65.67e6 -B --max-gap 0 -p 0.05 --keep-dup all --nomodel --call-summits. Peak refinement and annotation was carried out using a custom R pipeline incorporating the packages utils (v4.3.1), tidyverse (v2.0.0), tidyr (v1.3.1), readxl (v1.4.3), and dplyr (v1.1.4). Peaks containing two summits separated by >1000 bp with a pileup difference <40 % were split into distinct peaks. Reproducibility was assessed using IDR2D (v1.14.0). Peaks reproducible at p < 0.05 and those also identified in the BDP1 control (fold enrichment >1.5 and p < 0.05) were assigned to gene IDs based on proximity to the nearest TSS, accounting for gene orientation, using plyr (v1.8.9), DescTools (v0.99.58), ape (v5.8.1), openxlsx (v4.2.7.1), GenomicRanges (v1.52.1) and stringr (v1.5.1). Peaks equidistant to two genes (<1.5-fold difference) were assigned to both. Differential binding analysis was performed using DiffBind (v3.10.1) with DESeq2 as the statistical framework and a minimum quality control threshold of 15. Regions were defined as windows centered on the peak summit with a total length corresponding to two mean fragment sizes. Differential enrichment between *Tg*BRM and *Tg*BRG1 was defined using the Benjamini-Hochberg adjusted *p*-value combined with a fold change cutoff: peaks with adjusted *p*-value < 0.05 and fold change > 2 were classified as *Tg*BRM enriched, while those with adjusted *p*-value < 0.05 and fold change < –2 were classified as *Tg*BRG1 enriched. For RNA-seq integration, if multiple peaks were associated with a gene, only the closest peak was retained.

### Phylogenetic analysis

Phylogenetic trees were generated based on the identification of Myb domains (SM00717) in *T. gondii* as described in our previous work^18^. Proteins containing a DEXDc (SM00487) and a HELICc (SM00490) domain were identified in a broadly representative group of apicomplexan species including *T. gondii, E. tenella, P. falciparum, P. berghei, T. annulata, B. microti, C. parvum and G. niphandrodes* and the related species *C. velia*, through a homology-based search with SMART 9.0 (e-value < 0.05). The initial set of 604 DEXDc+HELICc domain–containing proteins was further enhanced by 40 additional DEXDc+HELICc domain–containing proteins using BLASTP^78,79^ against all analyzed species and filtering by domain length of the initial protein set identified by SMART (min. length = 0.5x DEXDc: 81 AA, 0.5x HELICc: 35 AA; max. length = 1.5x DEXDc: 1848 AA, 1.5x HELICc: 398 AA). Reciprocal BLASTP using the newly identified DEXDc+HELICc domain–containing proteins resulted in the identification of 3 additional DEXDc+HELICc domain–containing proteins. An additional reciprocal BLASTP search did not result in any additional hits resulting in 647 DEXDc+HELICc domain–containing of which 85 were *Toxoplasma*. We performed phylogenetic analysis on the 17 *Toxoplasma* proteins that showed the highest sequence similarity to *Tg*BRG1 and *Tg*BRM (e-value cutoff: 0.01). To generate a maximum likelihood (ML) phylogenetic tree we concatenated all DEXDc and HELICc domains for each protein and aligned them using the MUSCLE implementation of MEGA11. The best model to construct the ML tree was calculated using ModelTest-NG (v0.2.0). The best scoring model available in MEGA11 (LG+G4+F) was used to construct the ML tree using MEGA11 with 1000 bootstrap replicates. The closest relative to *Tg*BRG1 and *Tg*BRM with an e-value > 0.01 (TGME49_231800) was used as outgroup.

### Homology based searches

Whole-proteome FASTA files were obtained from EuPathDB (v68) and NCBI RefSeq databases (release 225). Protein datasets were formatted for local sequence similarity searches using makeblastdb from the BLAST+ suite (v2.15.0)^78,79^. Homology searches were performed locally using BLASTP from the BLAST+ suite, with each protein used as a query. Searches were conducted with parameters optimized for sensitivity, using a word size of 5 and a maximum of 10,000 alignments. For each species, the best hit corresponding to the lowest e-value was retained for subsequent analysis to assess evolutionary conservation and lineage-specific retention.

### Immunoblotting

*Tg*BRG1-mAID/*Tg*BRM-2×ALFA and *Tg*BRG1-2×ALFA/*Tg*BRM-mAID parasites were grown in standard media for 24 hr followed by treatment with either vehicle or 50 µM auxin for the duration of the indicated time points (0, 3, 6, 12 and 24 hr). 2×ALFA-TGGT1_203950-mNG-V5-mAID-Ty parasites were grown in standard media for 24 hr. About 10^6^ intracellular parasites were scraped, passed through a 27-gauge needle, filtered through a 5 µm filter and pelleted. The parasite pellets were resuspended in Laemmli buffer (2 % SDS, 10 % glycerol, 60 mM Tris HCl pH 6.8, 0.01 % bromophenol blue and 1 % β-mercaptoethanol) and run on a 4–15 % SDS-PAGE gel (BioRad). Proteins were transferred onto a nitrocellulose membrane using the Trans-Blot® Turbo™ Transfer System and the high or mixed molecular weight protocol (high: 1.3A and up to 25V for 10 min; mixed: 1.3A and up to 25V for 7 min). Membrane blocking and subsequent antibody probing was performed with rocking for 1 hr at room temperature in TBST (Tris-buffered saline with 0.1 % Tween-20) containing 5 % milk. Between primary and secondary antibody probing and before imaging, membranes were washed three times with TBST. Horseradish peroxidase-conjugated secondary antibodies were imaged using the Radiance Plus Femtogram HRP substrate (Azure biosystems) and a ChemiDoc imaging system (BioRad).

### Chromatin feature extraction and analysis

Wig and bigwig files for chromatin analysis were obtained from Sindikubwabo et al., 2017^47^ (H3K9me3: GSM2612412, H4K20me3: GSM2612413, H3K4me1: GSM2612414, H3K4me3: GSM2612415, H3K14ac: GSM2612416, H4K31me1: GSM2612417, H4K31ac: GSM2612418) as well as Nardelli et al., 2022^40^ (H3.3: GSM2795864, H2AX: GSM2795867, H2AZ: GSM2795868, H2BV: GSM2795870) and Farhat et al., 2020^7^ (MORC: [GSM4040359](https://www.ncbi.nlm.nih.gov/geo/query/acc.cgi?acc=GSM2795870), HDAC3: GSM4040363). Wig and bigwig files were analyzed using a custom R pipeline relying on the R packages wig (v0.1.0), GenomicRanges (v1.52.1), ape (v5.8.1), scales (v1.3.0) and rtracklayer (v1.60.1). Signal intensities from wig and bigwig files were quantified over gene bodies as annotated in the *T. gondii* ME49 reference genome (ToxoDB v68) and *Tg*BRG1/*Tg*BRM consensus peaks and averaged per region. Gene body and SWI/SNF consensus promoter signals were log_2_ transformed, mean centered and filtered to retain chromatin features of interest, followed by calculation of Pearson correlations to assess relationships between chromatin marks and remodeler occupancy.

Chromatin features predictive of transcriptional changes following *Tg*BRG1, *Tg*BRM or *Tg*SWI3 knockdown were identified using a custom R pipeline relying on the R package dplyr (v1.1.4). We filtered genes for sufficient expression (normalized counts > 37.93) and retained those containing a SWI/SNF consensus promoter signal. We selected both promoter-associated and gene body chromatin features, including histone modifications and remodeler occupancy, as predictors. A weighted multiple linear regression model was then fitted, using the respective log_2_ normalized fold changes as the response and −log_10_(adjusted *p*-values) as weights. Model coefficients were extracted and visualized with confidence intervals to assess the direction and strength of each feature’s contribution.

### Data analysis and visualization

All data analyses were performed using RStudio (v2023.06.0+421) and R (v4.3.1). Data manipulation and visualization were conducted using standard R packages, including ggplot2 (v3.5.2), ggbeeswarm (v0.7.2), ggpubr (v0.6.0), ggradar (v0.2), VennDiagram (v1.7.3), ComplexHeatmap (v2.16.0), scales (v1.3.0), RColorBrewer (v1.1.3) and viridis (v0.6.5) for plotting along with utils (v4.3.1), tidyverse (v2.0.0), tidyr (v1.3.1), readxl (v1.4.3), and dplyr (v1.1.4), fgsea (v1.26.0), tibble (v3.2.1) ineq (v0.2.13), purrr (v1.0.4) and reshape2 (v1.4.4) for data wrangling and visualization. Fold change data for *T. gondii* sexual stage transcriptomes were obtained from ToxoDB (www.toxodb.org) using the “Fold Change” option without applying any significance or fold change thresholds. Fold change data for *T. gondii* extracellular transcriptomes were obtained from Hassan et al., 2017^80^. Chromatin profiling sequencing reads for *Tg*MORC were obtained from Farhat et al., 2020^7^ (GSE136060). The analysis pipelines supporting the current study are available from the corresponding author upon request.

## ACKNOWLEDGEMENTS

We thank Thomas Schwartz and Cesar Dominguez for providing access to the microfluidizer, and Dylan Valleau, Fabian Schulte, and Brook Linnehan for their valuable support with mass spectrometry. We are grateful to Aditi Shukla for assistance with RNA-seq analysis, and to Mary Gehring, Steve Bell, and George Bell for their insightful discussions and scientific guidance. We also thank Faye Harling, Emily Shortt, Adel Misherghi, and Jake Altomare for their dedicated technical support. Finally, we acknowledge VEuPathDB as an essential resource for this work and thank all its contributors. This work was supported by NIH grants AI144369 and AI158501, and by the Burroughs Wellcome Fund (grant 1021330) awarded to S.L.; by the NIH training grant 5T32GM007287 to the Department of Biology at MIT; and by the American Heart Association predoctoral fellowship 898634 awarded to D.S.

## EXTENDED DATA FIGURES & LEGENDS

**Extended Data Figure 1.**
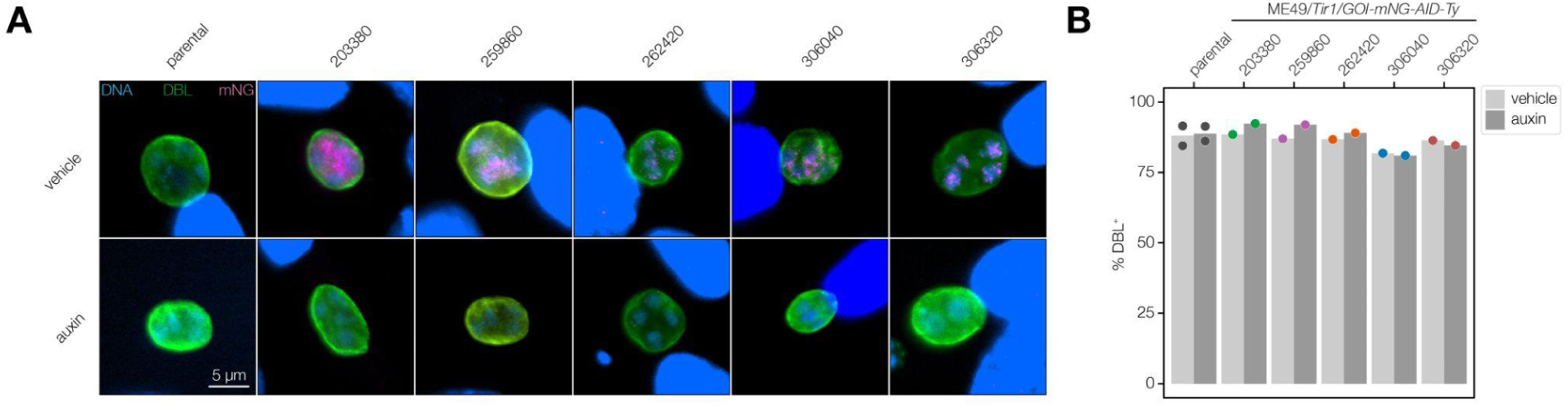
Analysis of Myb proteins without evident lytic cycle defects for potential function in chronic stage differentiation. **A.** Representative images of differentiation assays performed on all Myb proteins that did not exhibit fitness defects in the lytic cycle. Fixed parasites were probed with antibodies against mNG (magenta) as well as DBL-575 (green) and Hoechst (blue) to label the parasite’s cyst wall or DNA, respectively. **B.** Quantification of DBL positive vacuoles. Vacuoles were quantified based on immunofluorescence analysis (A).

**Extended Data Figure 2.**
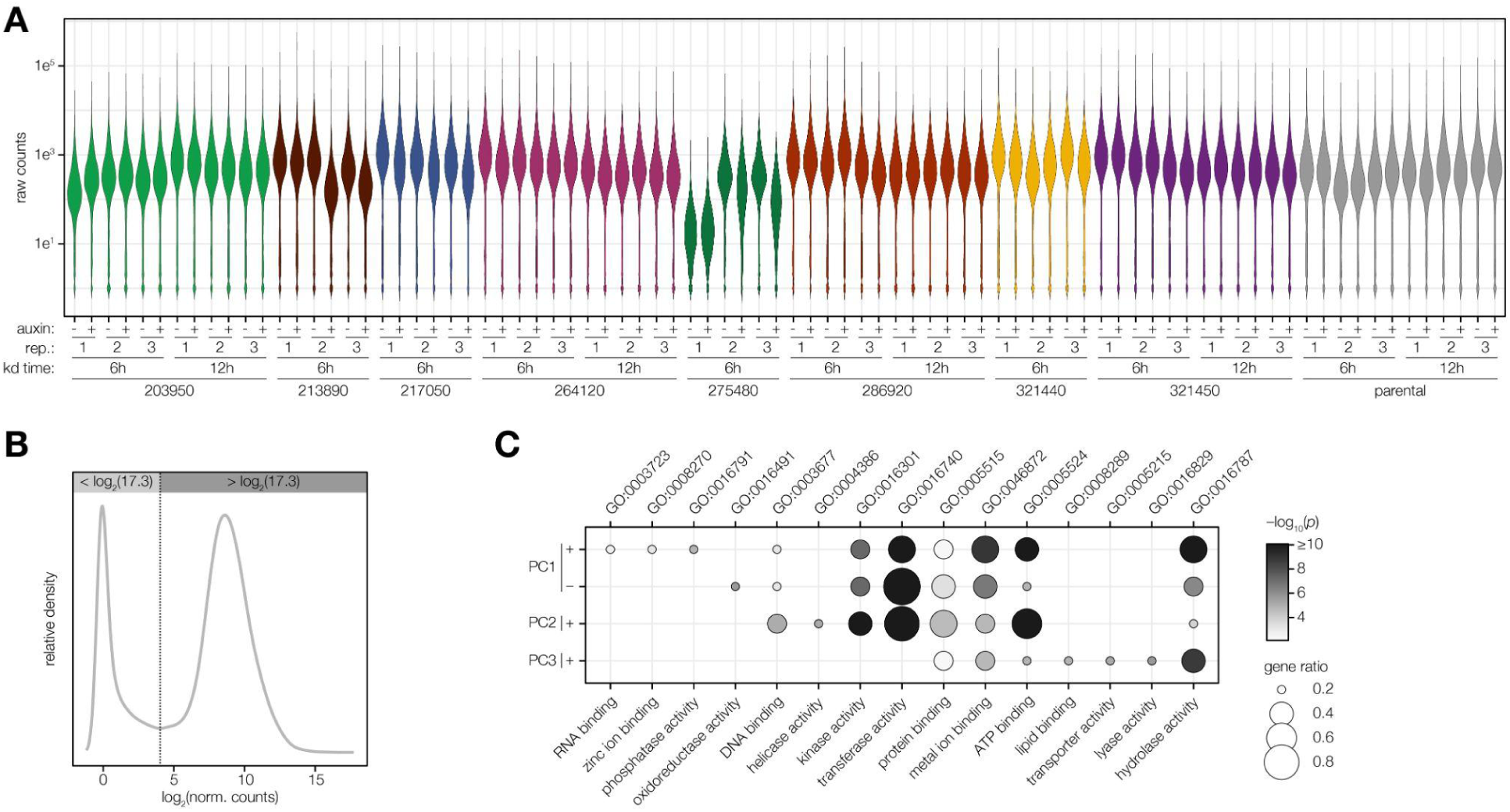
RNA-seq quality assessment and thresholding. **A.** Distribution of raw read counts across all RNA-seq samples relevant for Figure 2, showing consistency in overall sequencing depth and library complexity. **B.** Distribution of normalized read counts used to identify lowly expressed transcripts with high variance; these were excluded from downstream analyses to reduce noise. **C.** Gene ontology (GO) enrichment of PCA loadings corresponding to Figure 2A. The gene ratio is the proportion of genes with the indicated GO term divided by the total number of genes. The significance was determined with a hypergeometric test. Only GO terms with a *p*-value < 0.05 and a gene ratio > 0.1 are shown. Redundant GO terms were removed.

**Extended Data Figure 3.**
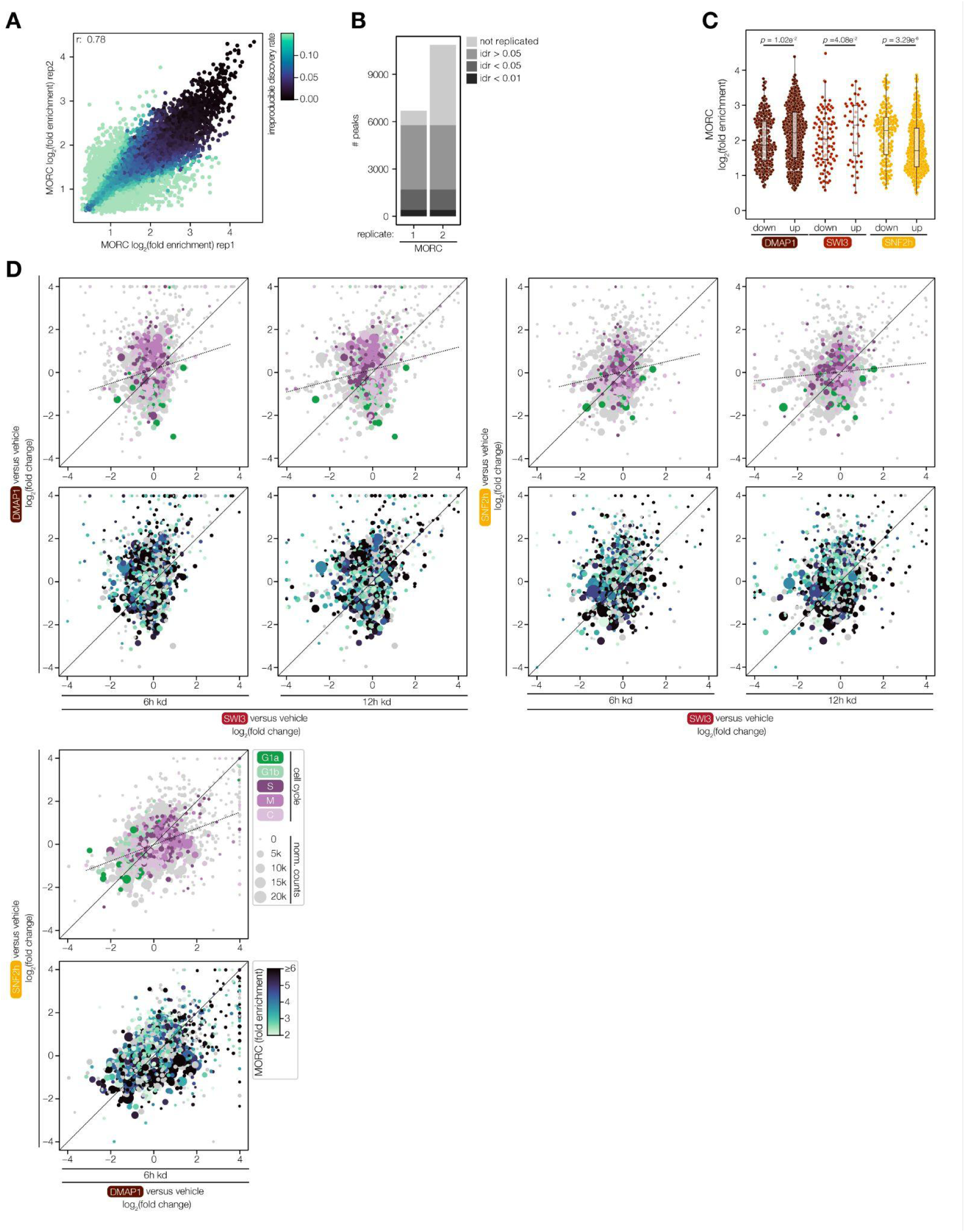
Myb protein chromatin remodelers cooperate with *Tg*MORC. **A.** Reproducibility analysis of published *Tg*MORC ChIP-seq data^7^ using the IDR framework to identify loci consistently enriched across biological replicates. Peaks are colored according to IDR-calculated *p*-values. Pearson correlation coefficient (r) is indicated. **B.** Number of peaks identified per *Tg*MORC ChIP-seq replicate. Bars represent the total number of peaks called by MACS2 in each replicate, with replicated peaks indicated in a darker shade according to IDR-calculated *p*-values. **C.** *Tg*MORC occupancy at promoter regions of genes differentially regulated after 6 hr knockdown of the indicated chromatin regulators. Statistical significance between up- and downregulated genes in *Tg*MORC occupancy was assessed using the Wilcoxon rank-sum test. **D.** Pairwise comparisons of transcriptomic changes following 6 hr and 12 hr knockdown of proteins representing components of three distinct ATP-dependent chromatin remodeling complexes. Scatter plots display log₂ fold changes from RNA-seq data of genes with an adjusted *p*-value < 0.005 in either sample. Genes are color-coded according to their cell cycle–regulated classification or by *Tg*MORC fold enrichment in the associated promoter region. Dot size corresponds to the mean RNA-seq counts in vehicle-treated samples of the corresponding strains.

**Extended Data Figure 4.**
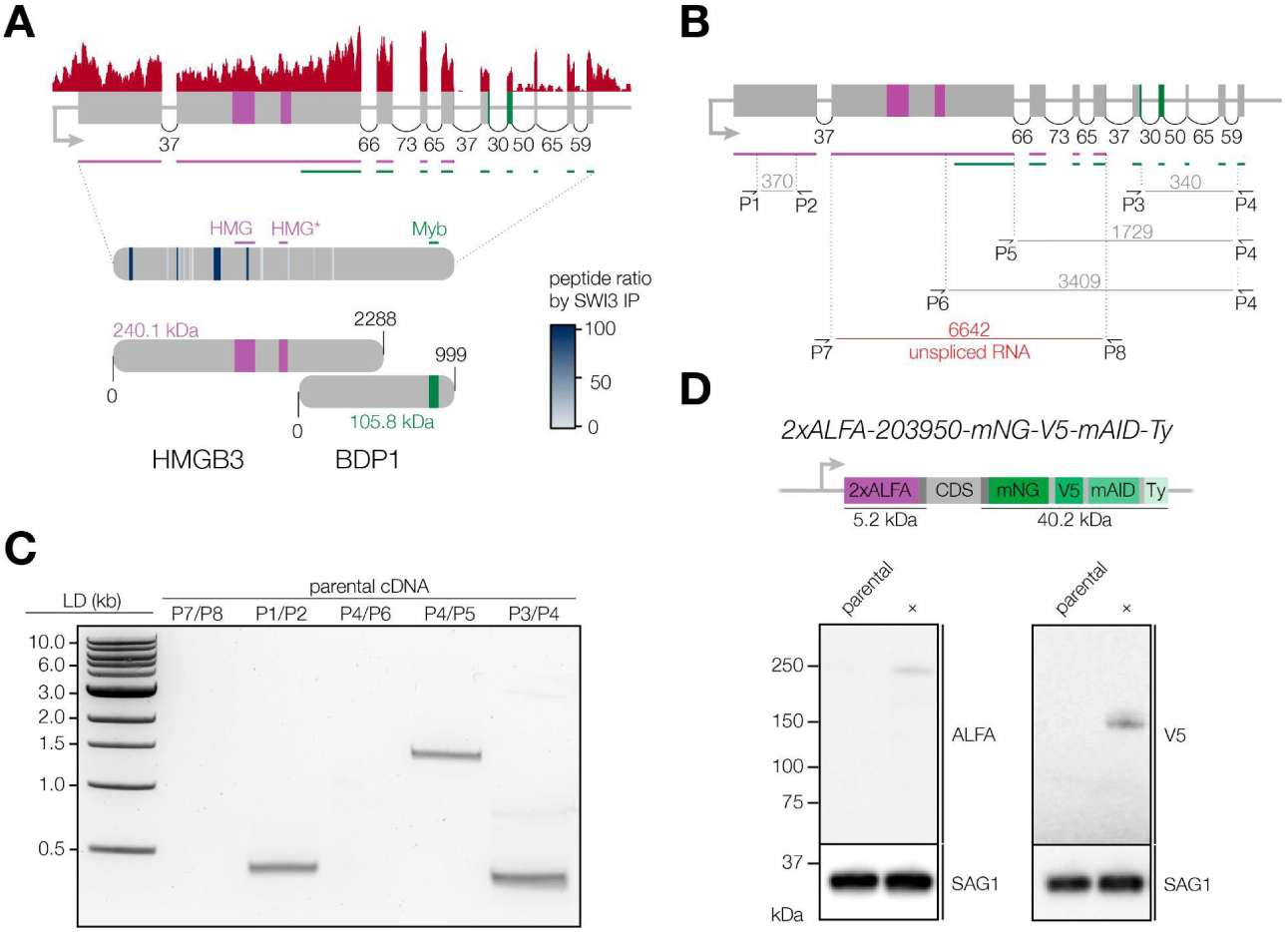
The annotated *Tg*BDP1 locus encodes two different proteins. **A.** RNA-seq read coverage (red) and annotated splicing junction spanning reads from the 6 hr parental RNA-seq sample are shown across the gene model of TGME49_203950 with HMG domains annotated in purple and the Myb domain annotated in green. Peptide coverage from the *Tg*SWI3 co-IPs is indicated in shades of blue. The peptide mapping is displayed in the context of the current gene model, and the proposed revised model, suggesting two distinct transcripts, is shown below. **B.** Schematic of primer pairs designed to test for transcript continuity across the TGME49_203950 locus by cDNA PCR. The expected amplicon sizes are indicated in bp. **C.** Agarose gel of cDNA PCR results, confirming that the N- and C-terminal regions of TGME49_203950 are expressed as separate transcripts. **D.** Schematic of the double-tagged strain with 2×ALFA and mNG-V5-mAID-Ty epitope tags at the N- and C-termini, respectively, and corresponding immunoblot analysis. Blots were probed with anti-ALFA and anti-V5 to confirm the size of the N- and C-terminal portions, and anti-SAG1 as loading control.

**Extended Data Figure 5.**
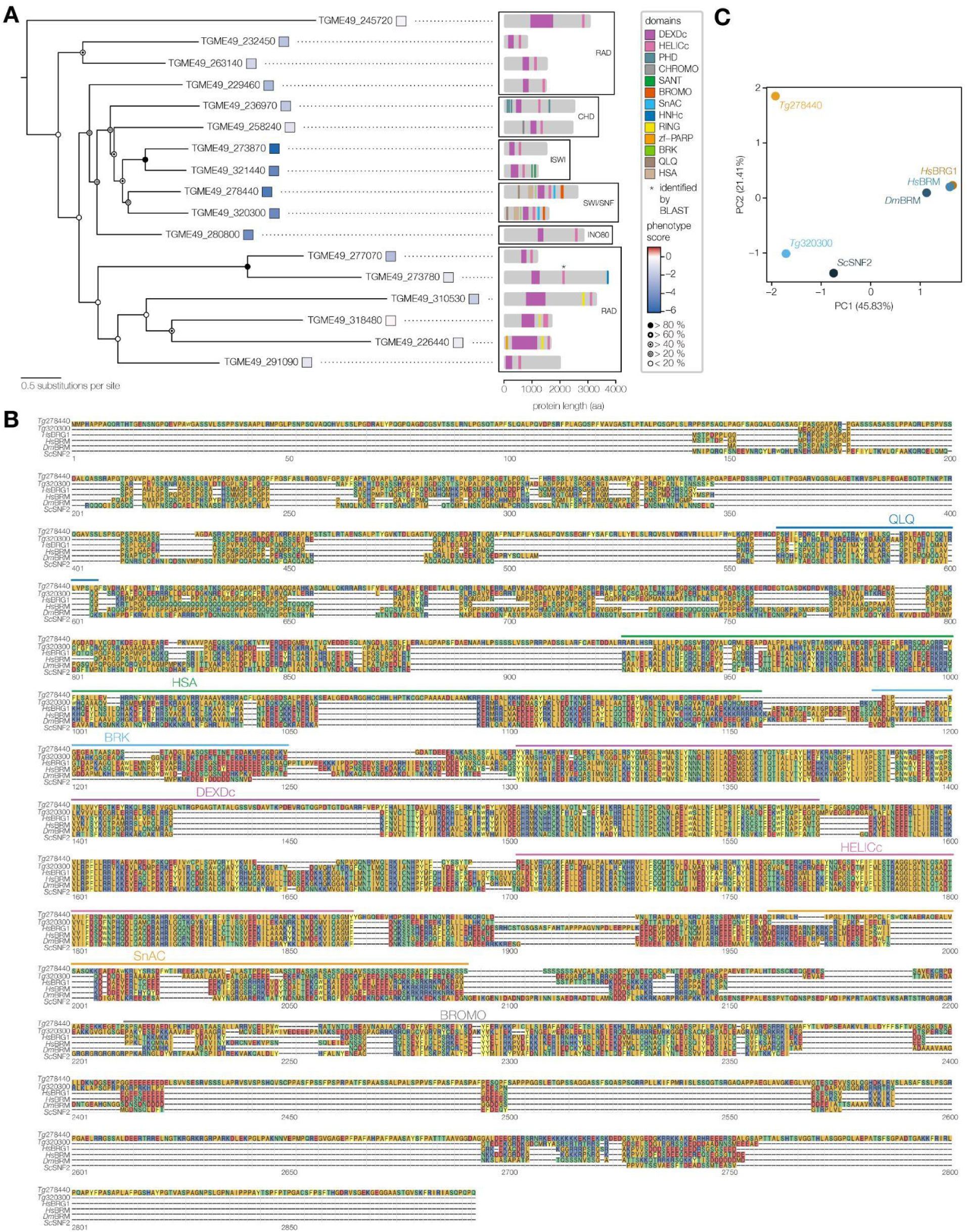
Classification of *Toxoplasma* DEXDc and HELICc domain containing proteins related to *Tg*BRG1 and *Tg*BRM. **A.** Phylogenetic tree drawn with DEXDc and HELICc domains of the 17 most closely related (e-value cutoff: 0.01) *Toxoplasma* proteins to TGME49_278440 and TGME49_320300. Phenotype scores derived from a genome-wide CRISPR screen^23^, reflecting the relative fitness contributions for each gene are indicated by colored squares. Clades as well as the domain architecture of the corresponding proteins are indicated with protein length in amino acids (aa). Bootstrap values of 1000 replicates are represented by circles. **B.** Multiple sequence alignment (MSA) of SWI/SNF helicases across species including *T. gondii* TGME49_278440 and TGME49_320300, *Homo sapiens* BRG1 (SMARCA4) and BRM (SMARCA2), *Drosophila melanogaster* BRM, and *Saccharomyces cerevisiae* SNF2. Conserved regions are highlighted. **C.** PCA of SWI/SNF helicase sequence similarity using full length pairwise alignment scores (BLOSUM62) between *T. gondii* TGME49_278440 and TGME49_320300, *H. sapiens* BRG1 and BRM, *D. melanogaster* BRM, and *S. cerevisiae* SNF2.

**Extended Data Figure 6.**
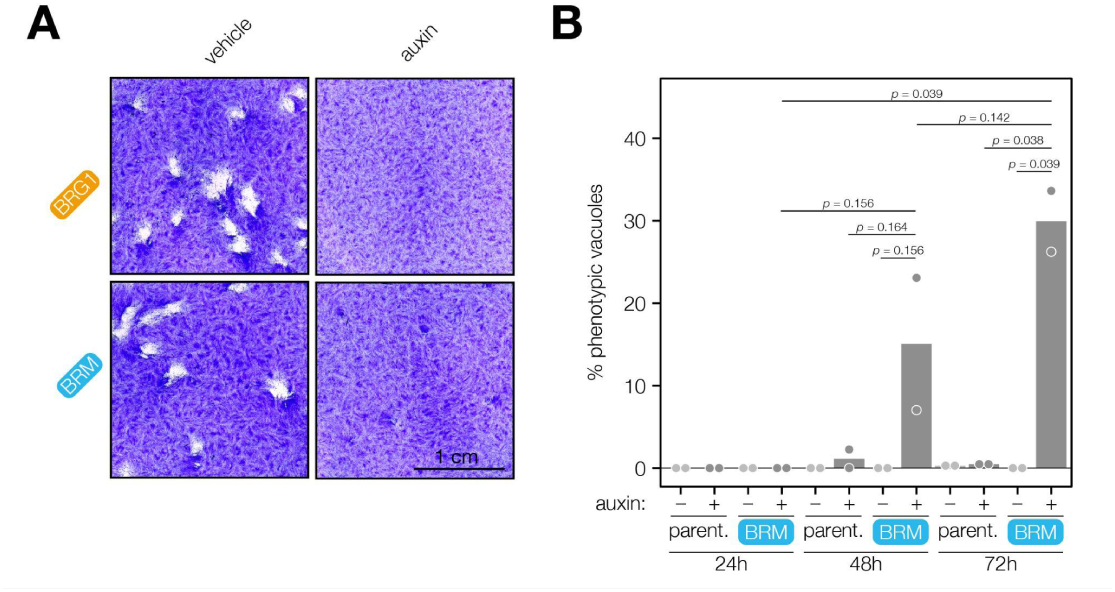
*Tg*BRM depletion results in delayed phenotypic effects, in contrast to the acute consequences of *Tg*BRG1 loss. **A.** Plaque assays of auxin induced *Tg*BRG1 or *Tg*BRM knockdown compared to vehicle-treated parasites. **B.** Quantification of morphological defects in *Tg*BRM knockdown parasites related to Figure 3F. Defects were assessed by immunofluorescence staining using antibodies against GAP45, IMC1, and mNG, with Hoechst labeling DNA. Data represent the mean from two independent blinded replicates, each analyzing at least 100 vacuoles per condition. Statistical significance was assessed using Welch’s one-tailed *t-*tests.

**Extended Data Figure 7.**
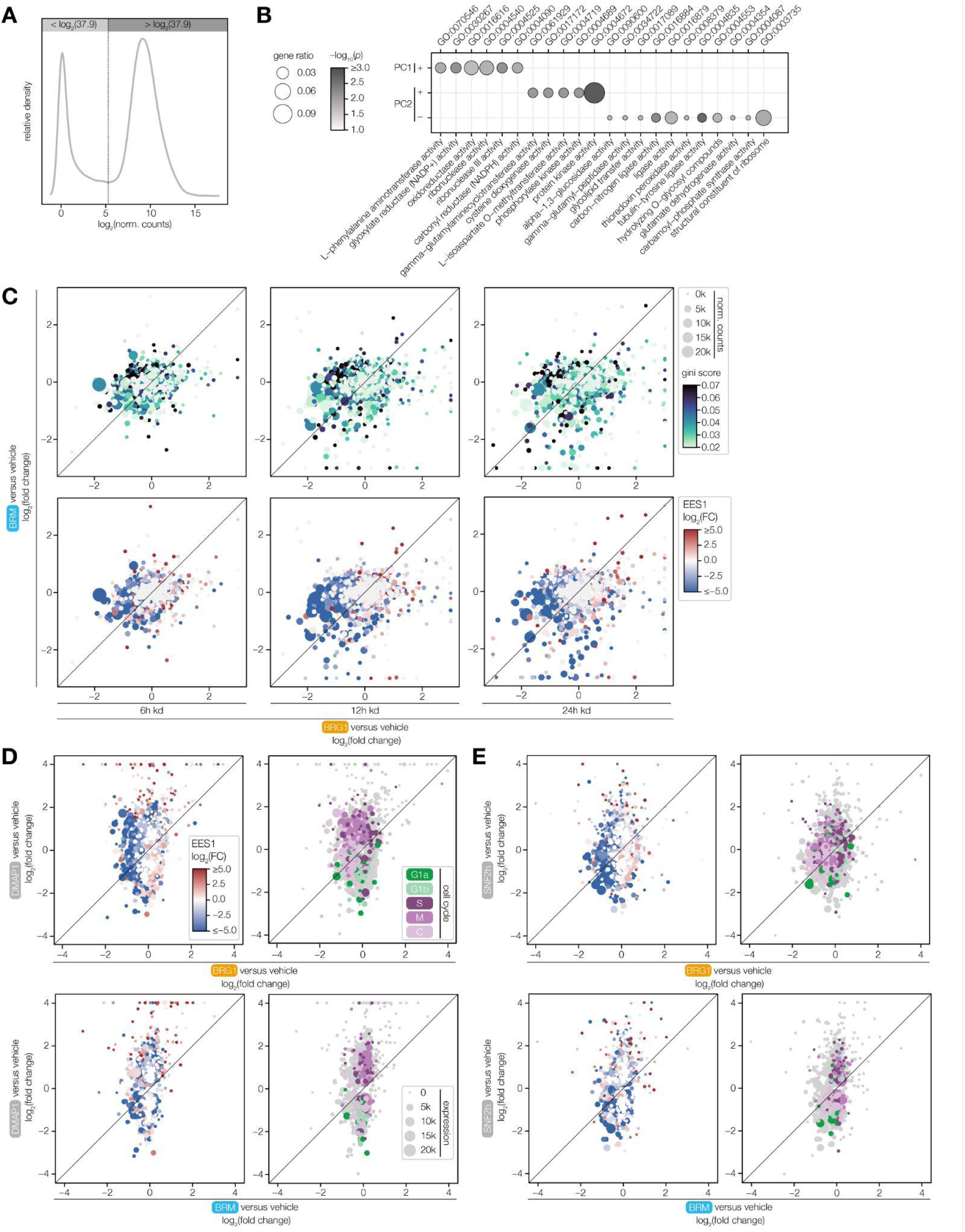
SWI/SNF helicases and other chromatin remodelers differentially regulate developmental and cell cycle–specific transcripts. **A.** Distribution of normalized read counts used to identify lowly expressed transcripts with high variance; these were excluded from downstream analyses to reduce noise. **B.** GO enrichment of PCA loadings corresponding to Figure 4B. The gene ratio is the proportion of genes with the indicated GO term divided by the total number of genes. The significance was determined with a hypergeometric test. Only GO terms with a *p*-value < 0.025 are shown. Redundant GO terms were removed. **C.** Pairwise comparisons of transcriptomic changes following 6, 12 and 24 hr knockdown of *Tg*BRG1 and *Tg*BRM. Scatter plots display log₂ fold changes from RNA-seq data. Genes are color-coded according to their Gini coefficient calculated based on transcriptome data from synchronized parasites according to Behnke et al.^39^ (top) or by their log₂ fold change in EES1-stage parasites relative to tachyzoites, as determined by Ramakrishnan et al.^60^ (bottom). Dot size corresponds to the mean RNA-seq counts in vehicle-treated samples of the corresponding strains. **D–E.** Pairwise comparisons of transcriptomic changes following 6 hr knockdown of *Tg*DMAP1 and both SWI/SNF helicases (D) and *Tg*SNF2h and SWI/SNF helicases (E). Scatter plots display log₂ fold changes from RNA-seq data of genes with an adjusted *p*-value < 0.005. Genes are color-coded according to their log₂ fold change in EES1-stage parasites relative to tachyzoites, as determined by Ramakrishnan et al.^60^ or by their cell cycle–regulated classification as determined by Xue et al.^37^. Dot size corresponds to the mean RNA-seq counts in vehicle-treated samples of the corresponding strains.

**Extended Data Figure 8.**
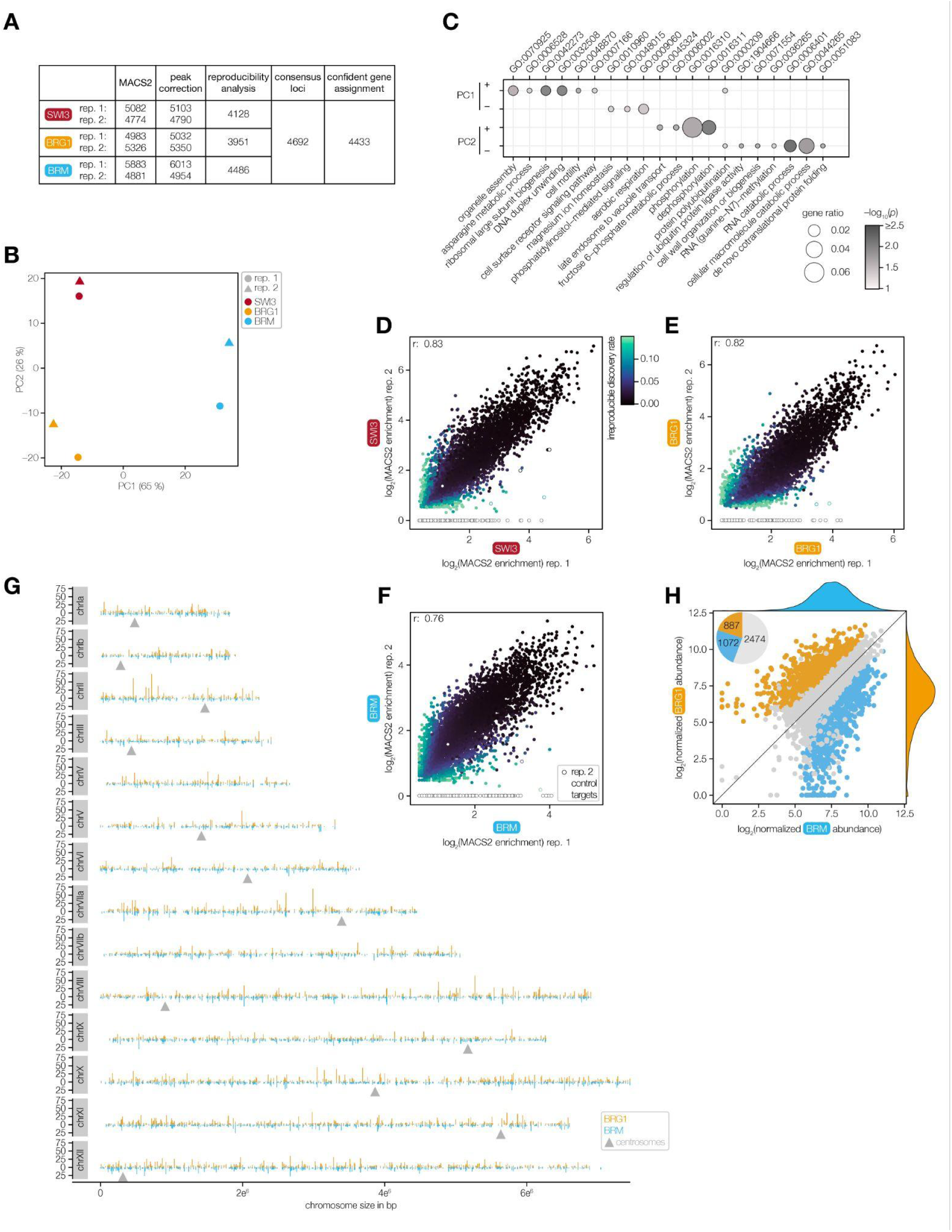
ALFA-CUT&RUN robustly identifies distinct binding profiles of SWI/SNF subunits in *Toxoplasma*. **A.** Summary table of peak counts identified for *Tg*SWI3, *Tg*BRG1, and *Tg*BRM following MACS2 peak calling, peak correction, and reproducibility analysis across individual replicates. **B.** PCA of all CUT&RUN replicates based on normalized read counts using the 1000 most variable consensus peaks across all samples. The first two principal components (PC1–PC2) are shown, representing the major axes of transcriptional variation. **C.** GO enrichment of PCA loadings corresponding to A. The gene ratio is the proportion of genes with the indicated GO term divided by the total number of genes. The significance was determined with a hypergeometric test. Only GO terms with a *p*-value < 0.05 are shown. Redundant GO terms were removed. **D–F.** Pairwise comparisons of read enrichment over the background per peak area in replicates 1 and 2 determined by MACS2 for *Tg*SWI3 (D), *Tg*BRG1 (E), and *Tg*BRM (F). Points are colored by IDR *p*-value; *Tg*BDP1 target loci are highlighted as circles and pearson correlation coefficients (r) are indicated. **G.** Genomic distribution of *Tg*BRG1 and *Tg*BRM consensus peaks across the *T. gondii* genome. Chromosomes and the location of centrosomes are indicated. **H.** Comparison of normalized log₂ read abundance between *Tg*BRG1 and *Tg*BRM at consensus peaks. Points represent single genomic consensus loci and are colored by enrichment category: *Tg*BRM-enriched loci (blue) were defined by adjusted *p*-value < 0.05 and a fold change > 2; *Tg*BRG1-enriched loci (orange) were defined by adjusted *p*-value < 0.05 and a fold change < –2. The number of loci in each category is shown in the accompanying pie chart.

**Extended Data Figure 9.**
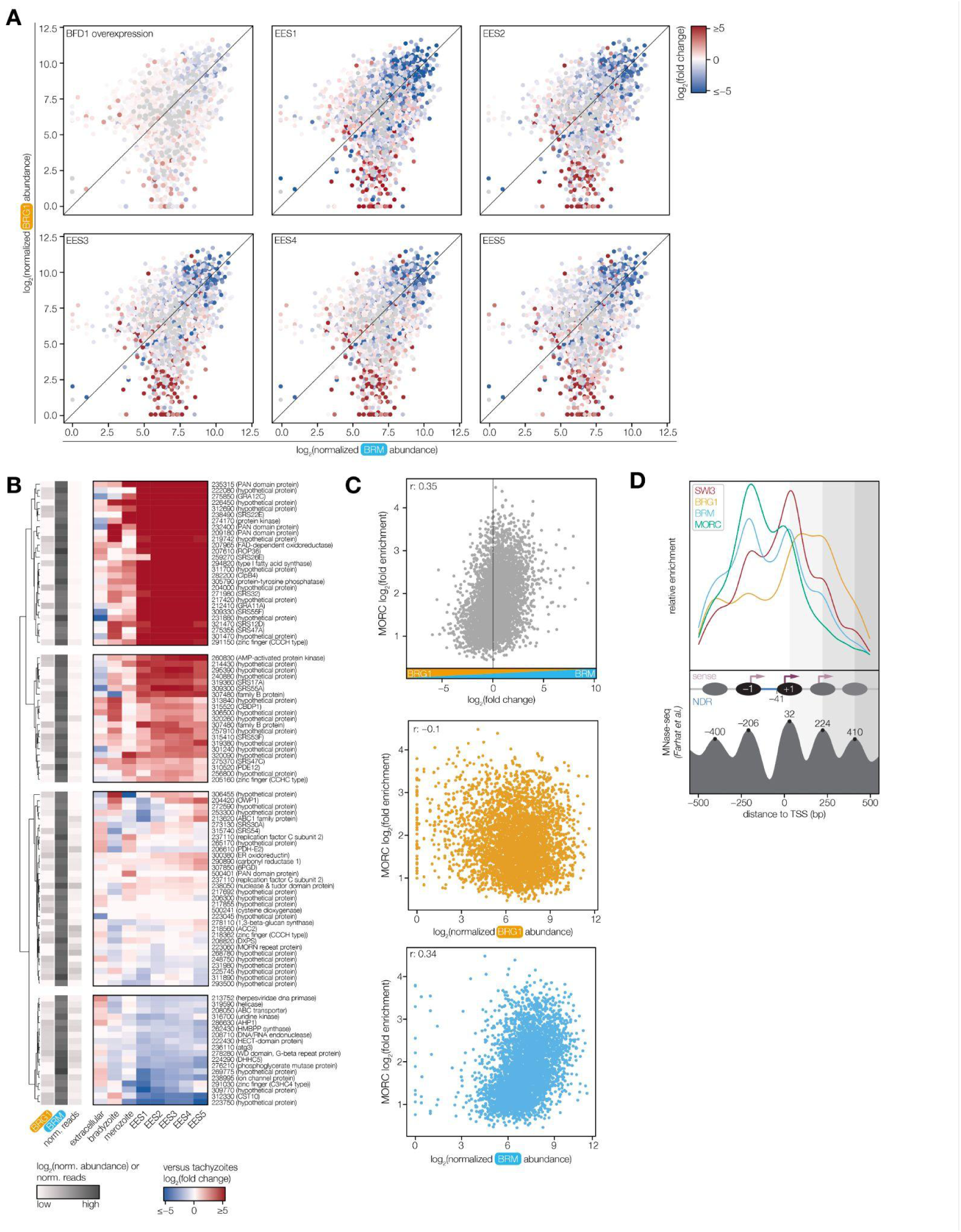
*Tg*BRM and *Tg*MORC but not *Tg*BRG1 occupy developmental loci associated with *Toxoplasma* differentiation. **A.** Comparison of *Tg*BRG1 and *Tg*BRM log₂ abundance based on normalized read counts after background signal subtraction. Points are colored according to the log₂ fold change in RNA levels of BFD1 differentiated parasites relative to tachyzoites determined by Waldman et al.^3^ and by log₂ fold changes in RNA levels of EES1–5 relative to tachyzoites as determined by Ramakrishnan et al.^60^. **B.** Heatmap showing the expression profiles of genes associated with selected loci (highlighted in Figure 5G; *Tg*BRG1 < 2 & *Tg*BRM > 5) across extracellular tachyzoites^80^, bradyzoites^60^, merozoites^85^, and EES1–5 stages^60^, each relative to intracellular tachyzoites. Rows correspond to individual genes, with *Tg*BRG1 and *Tg*BRM occupancy levels and normalized read counts indicated. **C.** Correlation between *Tg*MORC enrichment and the log₂ *Tg*BRM/*Tg*BRG1 abundance ratio across consensus loci. Pearson correlation coefficients (r) are indicated. Additional panels show the correlation of *Tg*MORC enrichment independently with *Tg*BRG1 and *Tg*BRM occupancy. **D.** Relative enrichment of *Tg*SWI3, *Tg*BRG1, *Tg*BRM and *Tg*MORC within ±500 bp of annotated TSSs. MNase-seq defined nucleosome positions from Farhat et al.^7,82^ are highlighted to indicate phased nucleosome occupancy relative to the nearest TSS.

**Extended Data Figure 10.**
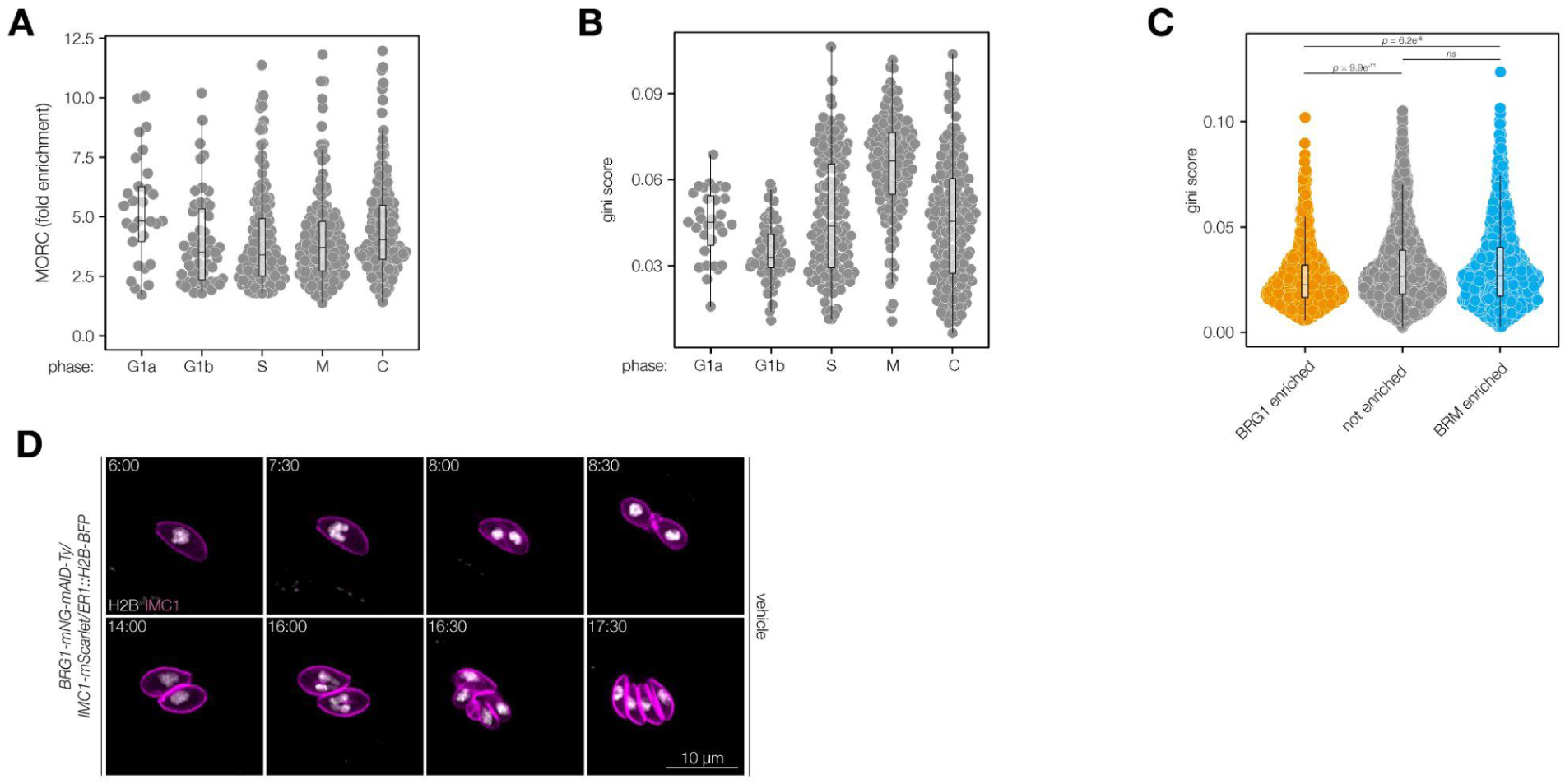
*Tg*MORC and *Tg*BRM preferentially associate with promoters of tightly regulated, short-phase cell cycle genes. **A–B.** *Tg*MORC enrichment (A) or Gini scores (B) of genes exhibiting phase-specific expression patterns, classified by single-cell RNA-sequencing^37^. *Tg*MORC enrichment is based on a reanalysis of published *Tg*MORC ChIP-seq data^7^. Gini scores were calculated from published transcriptome data obtained from synchronized parasites, with higher values indicating tightly phase-restricted transcription across the cell cycle^39^**. C.** Gini scores of genes showing significant enrichment for *Tg*BRG1, *Tg*BRM or no enrichment at consensus loci according to **Extended Data** Figure 8H. Gini scores were calculated from published transcriptome data obtained from synchronized parasites^39^. Differences between Gini scores of each category were assessed using the Wilcoxon rank-sum test. **D.** Live-cell imaging of vehicle-treated *Tg*BRG1 parasites expressing fluorescent markers for the inner membrane complex (IMC1-mScarlet, magenta) and chromatin (H2B-BFP, white). Parasites were imaged over a 24 hr period beginning 6 hr after vehicle treatment. See also **Video S3**.

**Extended Data Figure 11.**
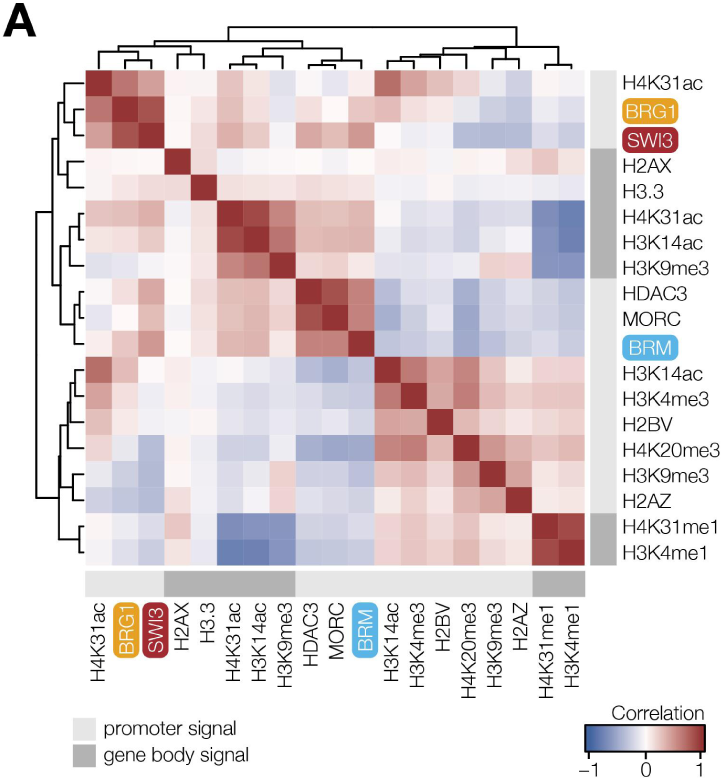
Expanded correlation map reveals two main chromatin clusters. **A.** Extended correlation heatmap of indicated chromatin-associated features across all genes associated with a consensus locus between *Tg*SWI3, *Tg*BRG1 and *Tg*BRM samples. Pairwise Pearson correlation coefficients were calculated using log_2_ transformed and mean-centered values. Light grey annotation bars indicate that the signal was measured over the promoter region within the SWI/SNF consensus locus, whereas dark grey bars denote that the signal was assessed across the gene body. Chromatin-associated features were derived from Farhat et al.^7^, Nardelli et al.^40^ and Sindikubwabo et al.^47^.

## EXTENDED DATA TABLES

**Supplementary Table 1.**

Quantification of competition assays of fitness-conferring Myb or SANT proteins comparing the relative growth of mAID-tagged strains against the parental strain expressing IMC1-tdTomato in auxin-containing media. At the indicated time points, the proportion of fluorescent parasites within each competing population was measured by flow cytometry and normalized to the vehicle control. Provided are relative abundances of each strain relative to the parental competitor strain. Related to Figure 1E.

**Supplementary Table 2.**

Quantification of centrosomes per parasite nucleus of CDC5L knockdown parasites. Centrosomes were quantified based on immunofluorescence analysis. Provided is the proportion of parasites with 0, 1, and 2 centrosomes per parasite nucleus after 24 hr of auxin-induced knockdown for ≥100 vacuoles per replicate. Related to Figure 1G.

**Supplementary Table 3.**

Quantification of DBL-positive vacuoles of BDP1 knockdown parasites. Vacuoles were quantified based on immunofluorescence analysis. Provided is the proportion of DBL-positive vacuoles quantified for ≥100 vacuoles per replicate after 48 hr of auxin-induced knockdown. Related to Figure 1I.

**Supplementary Table 4.**

Differentiation assay quantification after knockdown of Myb proteins without evident lytic cycle defects. Vacuoles were quantified based on immunofluorescence analysis. Provided is the proportion of DBL-positive vacuoles quantified for ≥100 vacuoles per replicate. Related to Extended Data Figure 1.

**Supplementary Table 5.**

Differential expression analysis of nuclear-localized and fitness-conferring Myb proteins following knockdown for 6 hr and 12 hr. The behavior of listed genes in prior transcriptional and chromatin profiling datasets is included. Related to Figure 2; Extended Data Figure 2–3, 7.

**Supplementary Table 6.**

Reproducibility analysis of published *Tg*MORC ChIP-seq data using the IDR framework to identify loci consistently enriched across biological replicates. Related to Extended Data Figure 3.

**Supplementary Table 7.**

Results from co-IP MS experiments of endogenously tagged *Tg*SWI3 with mNG and reciprocal co-IP MS of endogenously tagged *Tg*BRG1-2×ALFA and *Tg*BRM-2×ALFA. Related to Figure 3A.

**Supplementary Table 8.**

Quantification of competition assays of *Tg*SWI3, *Tg*BRG1, and *Tg*BRM mAID-tagged strains comparing the relative growth against the parental strain expressing IMC1-tdTomato in auxin-containing media. At the indicated time points, the proportion of fluorescent parasites within each competing population was measured by flow cytometry and normalized to the vehicle control. Provided are relative abundances of each strain relative to the parental competitor strain. Related to Figure 3E.

**Supplementary Table 9.**

Quantification of morphological defects in *Tg*BRM knockdown parasites. Defects were assessed by immunofluorescence staining. Provided is the proportion of phenotypic vacuoles quantified for ≥100 vacuoles per replicate. Related to Extended Data Figure 6B.

**Supplementary Table 10.**

Differential expression analysis of *Tg*BRG1, *Tg*BRM, and *Tg*SWI3 parasites following knockdown for 6, 12, and 24 hr. CUT&RUN data (columns labeled as “CR_”) as well as the behavior of listed genes in prior transcriptional and chromatin profiling datasets are included. Related to Figure 4, 7; Extended Data Figure 7.

**Supplementary Table 11.**

Reproducibility analysis of *Tg*BRG1, *Tg*BRM, or *Tg*SWI3 CUT&RUN experiments using the IDR framework to identify loci consistently enriched across biological replicates. Related to Figure 5; Extended Data Figure 8.

**Supplementary Table 12.**

Differential binding analysis of *Tg*BRG1 & *Tg*BRM at consensus loci. Related to Figure 5–7; Extended Data Figure 8, 10.

**Supplementary Table 13.**

Quantification of nuclear mNG fluorescence intensity per pixel from live-cell video microscopy of endogenously tagged *Tg*BRG1. Fluorescence intensity of *n* = 4 biological replicates was quantified using CellProfiler. Related to Figure 6D.

**Supplementary Table 14.**

Quantification of nuclear mNG fluorescence intensity per pixel from live-cell video microscopy of endogenously tagged *Tg*BRM. Fluorescence intensity of *n* = 4 biological replicates was quantified using CellProfiler. Related to Figure 6E.

**Supplementary Table 15.**

Quantification of replication assays of *Tg*BRG1- and *Tg*BRM-mAID–tagged strains and the parental line in the presence or absence of auxin. Parasites per vacuole were quantified based on immunofluorescence images across *n* = 3 biological replicates. Provided is the proportion of vacuoles with 2, 4, 8, and ≥16 parasites for ≥100 vacuoles per replicate. Related to Figure 6F.

**Supplementary Table 16.**

Quantification of DNA content staining of parental as well as *Tg*BRG1 and *Tg*BRM knockdown strains 12, 24, and 48 hr after auxin treatment. Provided is the percentage of parasites with S, M, and C phase DNA content profiles across *n* = 3 biological replicates. Related to Figure 6H.

**Supplementary Table 17.**

Quantification of published chromatin-associated features across all genes associated with a consensus locus between *Tg*SWI3, *Tg*BRG1 and *Tg*BRM samples. GEO accession numbers relevant datasets are indicated. CUT&RUN data (columns labeled as “CR_”) are included. Related to Figure 7; Extended Data Figure 11A.

**Supplementary Table 18.**

All oligos used or generated in this study.

## EXTENDED DATA VIDEOS

**Video S1. *Tr*im knockdown cell cycle reporter parasites imaged 48 hr after auxin-induced knockdown.**

Cell cycle reporter strain engineered for live-cell imaging. IMC1 was endogenously tagged at the C terminus with mScarlet-I to visualize daughter cell formation and H2B fused to BFP was ectopically expressed to visualize nuclear dynamics. Live-cell imaging of *Tg*BRM knockdown parasites imaged over a 24-hr period beginning 48 hr after auxin-induced knockdown.

**Video S2. *Tg*BRG1 knockdown cell cycle reporter parasites imaged 6 hr after auxin-induced knockdown.**

Cell cycle reporter strain engineered for live-cell imaging. IMC1 was endogenously tagged at the C terminus with mScarlet-I to visualize daughter cell formation and H2B fused to BFP was ectopically expressed to visualize nuclear dynamics. Live-cell imaging of *Tg*BRG1 knockdown parasites imaged over a 24-hr period beginning 6 hr after auxin-induced knockdown.

**Video S3. *Tg*BRG1 knockdown cell cycle reporter parasites imaged 6 hr after vehicle treatment.**

Cell cycle reporter strain engineered for live-cell imaging. IMC1 was endogenously tagged at the C terminus with mScarlet-I to visualize daughter cell formation and H2B fused to BFP was ectopically expressed to visualize nuclear dynamics. Live-cell imaging of vehicle-treated *Tg*BRG1 parasites imaged over a 24-hr period beginning 6 hr after auxin-induced knockdown.

## References

1. Painter, H. J., Carrasquilla, M. & Llinás, M. Capturing in vivo RNA transcriptional dynamics from the malaria parasite Plasmodium falciparum. Genome Res. 27, 1074–1086 (2017).

2. Lippuner, C. et al. RNA-Seq analysis during the life cycle of Cryptosporidium parvum reveals significant differential gene expression between proliferating stages in the intestine and infectious sporozoites. Int. J. Parasitol. 48, 413–422 (2018).

3. Waldman, B. S. et al. Identification of a master regulator of differentiation in Toxoplasma. Cell 180, 359–372.e16 (2020).

4. Cheeseman, K. et al. Dynamic methylation of histone H3K18 in differentiating Theileria parasites. Nat. Commun. 12, 3221 (2021).

5. Salomaki, E. D. et al. Gregarine single-cell transcriptomics reveals differential mitochondrial remodeling and adaptation in apicomplexans. BMC Biol. 19, 77 (2021).

6. Lou, J. et al. Single cell expression and chromatin accessibility of the Toxoplasma gondii lytic cycle identifies AP2XII-8 as an essential ribosome regulon driver. Nat. Commun. 15, 7419 (2024).

7. Farhat, D. C. et al. A MORC-driven transcriptional switch controls Toxoplasma developmental trajectories and sexual commitment. Nat. Microbiol. 5, 570–583 (2020).

8. Pachano, B. et al. An ISWI-related chromatin remodeller regulates stage-specific gene expression in Toxoplasma gondii. Nat. Microbiol. (2025) doi:10.1038/s41564-025-01980-2.

9. Radke, J. R. et al. Defining the cell cycle for the tachyzoite stage of Toxoplasma gondii. Mol. Biochem. Parasitol. 115, 165–175 (2001).

10. Gubbels, M.-J., Coppens, I., Zarringhalam, K., Duraisingh, M. T. & Engelberg, K. The modular circuitry of Apicomplexan cell division plasticity. Front. Cell. Infect. Microbiol. 11, 670049 (2021).

11. Padilla, L. F. A., Murray, J. M. & Hu, K. The initiation and early development of the tubulin-containing cytoskeleton in the human parasite Toxoplasma gondii. Mol. Biol. Cell 35, ar37 (2024).

12. Alonso, A. M., Corvi, M. M. & Diambra, L. Gene target discovery with network analysis in Toxoplasma gondii. Sci. Rep. 9, 646 (2019).

13. Jeninga, M. D., Quinn, J. E. & Petter, M. ApiAP2 transcription factors in Apicomplexan parasites. Pathogens 8, 47 (2019).

14. Chen, Y. et al. Chromatin accessibility: biological functions, molecular mechanisms and therapeutic application. Signal Transduct. Target. Ther. 9, 340 (2024).

15. Saraf, A. et al. Dynamic and combinatorial landscape of histone modifications during the intraerythrocytic developmental cycle of the malaria parasite. J. Proteome Res. 15, 2787–2801 (2016).

16. Hollin, T. & Le Roch, K. G. From genes to transcripts, a tightly regulated journey in Plasmodium. Front. Cell. Infect. Microbiol. 10, 618454 (2020).

17. Zhu, Y. et al. Regulation of the developmental programs in Toxoplasma by a novel SNF2L-containing chromatin remodeling complex. Nat. Commun. 16, 5757 (2025).

18. Schwarz, D. & Lourido, S. The multifaceted roles of Myb domain-containing proteins in apicomplexan parasites. Curr. Opin. Microbiol. 76, 102395 (2023).

19. Gissot, M., Briquet, S., Refour, P., Boschet, C. & Vaquero, C. PfMyb1, a Plasmodium falciparum transcription factor, is required for intra-erythrocytic growth and controls key genes for cell cycle regulation. J. Mol. Biol. 346, 29–42 (2005).

20. Walzer, K. A. et al. Transcriptional control of the Cryptosporidium life cycle. Nature 630, 174–180 (2024).

21. Bieluszewski, T., Prakash, S., Roulé, T. & Wagner, D. The role and activity of SWI/SNF chromatin remodelers. Annu. Rev. Plant Biol. 74, 139–163 (2023).

22. Smith, T. A., Lopez-Perez, G. S., Herneisen, A. L., Shortt, E. & Lourido, S. Screening the Toxoplasma kinome with high-throughput tagging identifies a regulator of invasion and egress. Nat. Microbiol. 7, 868–881 (2022).

23. Sidik, S. M. et al. A genome-wide CRISPR screen in Toxoplasma identifies essential Apicomplexan genes. Cell 166, 1423–1435.e12 (2016).

24. Giannone, R. J. et al. The protein network surrounding the human telomere repeat binding factors TRF1, TRF2, and POT1. PLoS One 5, e12407 (2010).

25. Rosenzweig, R., Nillegoda, N. B., Mayer, M. P. & Bukau, B. The Hsp70 chaperone network. Nat. Rev. Mol. Cell Biol. 20, 665–680 (2019).

26. Zhang, S., Xie, M., Ren, G. & Yu, B. CDC5, a DNA binding protein, positively regulates posttranscriptional processing and/or transcription of primary microRNA transcripts. Proc. Natl. Acad. Sci. U. S. A. 110, 17588–17593 (2013).

27. Kastner, B., Will, C. L., Stark, H. & Lührmann, R. Structural insights into nuclear pre-mRNA splicing in higher eukaryotes. Cold Spring Harb. Perspect. Biol. 11, a032417 (2019).

28. Kashyap, P., Aswale, K. R. & Deshmukh, A. S. Deletion of splicing factor Cdc5 in Toxoplasma disrupts transcriptome integrity, induces abortive bradyzoite formation, and prevents acute infection in mice. Nat. Commun. 16, 3769 (2025).

29. Toenhake, C. G. et al. Chromatin accessibility-based characterization of the gene regulatory network underlying Plasmodium falciparum blood-stage development. Cell Host Microbe 23, 557–569.e9 (2018).

30. Hargreaves, D. C. & Crabtree, G. R. ATP-dependent chromatin remodeling: genetics, genomics and mechanisms. Cell Res. 21, 396–420 (2011).

31. He, S. et al. Structure of nucleosome-bound human BAF complex. Science 367, 875–881 (2020).

32. Gouge, J. et al. Molecular mechanisms of Bdp1 in TFIIIB assembly and RNA polymerase III transcription initiation. Nat. Commun. 8, 130 (2017).

33. Abramson, J. et al. Accurate structure prediction of biomolecular interactions with AlphaFold 3. Nature 630, 493–500 (2024).

34. Mashtalir, N. et al. Modular organization and assembly of SWI/SNF family chromatin remodeling complexes. Cell 175, 1272–1288.e20 (2018).

35. Loyola, A. C. et al. Streptonigrin at low concentration promotes heterochromatin formation. Sci. Rep. 10, 3478 (2020).

36. White, M. W. & Suvorova, E. S. Apicomplexa cell cycles: Something old, borrowed, lost, and new. Trends Parasitol. 34, 759–771 (2018).

37. Xue, Y. et al. A single-parasite transcriptional atlas of Toxoplasma Gondii reveals novel control of antigen expression. Elife 9, (2020).

38. Ceriani, L. & Verme, P. The origins of the Gini index: extracts from Variabilità e Mutabilità (1912) by Corrado Gini. J. Econ. Inequal. 10, 421–443 (2012).

39. Behnke, M. S. et al. Coordinated progression through two subtranscriptomes underlies the tachyzoite cycle of Toxoplasma gondii. PLoS One 5, e12354 (2010).

40. Nardelli, S. C. et al. Genome-wide localization of histone variants in Toxoplasma gondii implicates variant exchange in stage-specific gene expression. BMC Genomics 23, 128 (2022).

41. Bhaskaran, M., Mudiyam, V., Mouveaux, T., Roger, E. & Gissot, M. Cascading expression of ApiAP2 transcription factors controls daughter cell assembly in Toxoplasma gondii. PLoS Pathog. 20, e1012810 (2024).

42. Hainer, S. J. & Fazzio, T. G. High-resolution chromatin profiling using CUT&RUN. Curr. Protoc. Mol. Biol. 126, e85 (2019).

43. Meers, M. P., Bryson, T. D., Henikoff, J. G. & Henikoff, S. Improved CUT&RUN chromatin profiling tools. Elife 8, (2019).

44. Götzke, H. et al. The ALFA-tag is a highly versatile tool for nanobody-based bioscience applications. Nat. Commun. 10, 4403 (2019).

45. Ishiguro, A., Kassavetis, G. A. & Geiduschek, E. P. Essential roles of Bdp1, a subunit of RNA polymerase III initiation factor TFIIIB, in transcription and tRNA processing. Mol. Cell. Biol. 22, 3264–3275 (2002).

46. Li, Q., Brown, J. B., Huang, H. & Bickel, P. J. Measuring reproducibility of high-throughput experiments. aoas 5, 1752–1779 (2011).

47. Sindikubwabo, F. et al. Modifications at K31 on the lateral surface of histone H4 contribute to genome structure and expression in apicomplexan parasites. Elife 6, (2017).

48. Fraschka, S. A.-K., Henderson, R. W. M. & Bártfai, R. H3.3 demarcates GC-rich coding and subtelomeric regions and serves as potential memory mark for virulence gene expression in Plasmodium falciparum. Sci. Rep. 6, 31965 (2016).

49. Dalmasso, M. C., Onyango, D. O., Naguleswaran, A., Sullivan, W. J., Jr & Angel, S. O. Toxoplasma H2A variants reveal novel insights into nucleosome composition and functions for this histone family. J. Mol. Biol. 392, 33–47 (2009).

50. Purman, C. E. et al. Regional gene repression by DNA double-strand breaks in G1 phase cells. Mol. Cell. Biol. 39, 1–16 (2019).

51. Park, J.-H. et al. Mammalian SWI/SNF complexes facilitate DNA double-strand break repair by promoting gamma-H2AX induction. EMBO J. 25, 3986–3997 (2006).

52. Lee, H.-S., Park, J.-H., Kim, S.-J., Kwon, S.-J. & Kwon, J. A cooperative activation loop among SWI/SNF, gamma-H2AX and H3 acetylation for DNA double-strand break repair. EMBO J. 29, 1434–1445 (2010).

53. Solovjeva, L. V., Svetlova, M. P., Chagin, V. O. & Tomilin, N. V. Inhibition of transcription at radiation-induced nuclear foci of phosphorylated histone H2AX in mammalian cells. Chromosome Res. 15, 787–797 (2007).

54. Rountree, M. R., Bachman, K. E. & Baylin, S. B. DNMT1 binds HDAC2 and a new co-repressor, DMAP1, to form a complex at replication foci. Nat. Genet. 25, 269–277 (2000).

55. Gissot, M., Choi, S.-W., Thompson, R. F., Greally, J. M. & Kim, K. Toxoplasma gondii and Cryptosporidium parvum lack detectable DNA cytosine methylation. Eukaryot. Cell 7, 537–540 (2008).

56. Wei, H. et al. Characterization of Cytosine Methylation and the DNA Methyltransferases of Toxoplasma gondii. Int. J. Biol. Sci. 13, 458–470 (2017).

57. Yu, J. et al. Structural insights into histone exchange by human SRCAP complex. Cell Discov. 10, 15 (2024).

58. Wiechens, N. et al. The chromatin remodelling enzymes SNF2H and SNF2L position nucleosomes adjacent to CTCF and other transcription factors. PLoS Genet. 12, e1005940 (2016).

59. Robertson, A. K., Geiman, T. M., Sankpal, U. T., Hager, G. L. & Robertson, K. D. Effects of chromatin structure on the enzymatic and DNA binding functions of DNA methyltransferases DNMT1 and Dnmt3a in vitro. Biochem. Biophys. Res. Commun. 322, 110–118 (2004).

60. Ramakrishnan, C. et al. An experimental genetically attenuated live vaccine to prevent transmission of Toxoplasma gondii by cats. Sci. Rep. 9, 1474 (2019).

61. Raab, J. R., Runge, J. S., Spear, C. C. & Magnuson, T. Co-regulation of transcription by BRG1 and BRM, two mutually exclusive SWI/SNF ATPase subunits. Epigenetics Chromatin 10, 62 (2017).

62. Fernando, T. M. et al. Functional characterization of SMARCA4 variants identified by targeted exome-sequencing of 131,668 cancer patients. Nat. Commun. 11, 5551 (2020).

63. Nishi, T., Kaneko, I., Iwanaga, S. & Yuda, M. PbARID-associated chromatin remodeling events are essential for gametocyte development in Plasmodium. Nucleic Acids Res. 52, 5624–5642 (2024).

64. Kumar, S. et al. PfARID regulates P. falciparum malaria parasite male gametogenesis and female fertility and is critical for parasite transmission to the mosquito vector. MBio 13, e0057822 (2022).

65. Zhu, Z. et al. Mitotic bookmarking by SWI/SNF subunits. Nature 618, 180–187 (2023).

66. Hawkins, L. M. et al. The Crk4-Cyc4 complex regulates G2/M transition in Toxoplasma gondii. EMBO J. 43, 2094–2126 (2024).

67. Barisic, D., Stadler, M. B., Iurlaro, M. & Schübeler, D. Mammalian ISWI and SWI/SNF selectively mediate binding of distinct transcription factors. Nature 569, 136–140 (2019).

68. Bindels, D. S. et al. mScarlet: a bright monomeric red fluorescent protein for cellular imaging. Nat. Methods 14, 53–56 (2017).

69. Heim, R. & Tsien, R. Y. Engineering green fluorescent protein for improved brightness, longer wavelengths and fluorescence resonance energy transfer. Curr. Biol. 6, 178–182 (1996).

70. Markus, B. M., Bell, G. W., Lorenzi, H. A. & Lourido, S. Optimizing Systems for Cas9 Expression in Toxoplasma gondii. mSphere 4, (2019).

71. Herneisen, A. L., Peters, M. L., Smith, T. A., Shortt, E. & Lourido, S. SPARK regulates AGC kinases central to the Toxoplasma gondii asexual cycle. Elife 13, (2024).

72. Brown, K. M., Long, S. & Sibley, L. D. Plasma membrane association by N-acylation governs PKG function in Toxoplasma gondii. MBio 8, (2017).

73. Licon, M. H. et al. A positive feedback loop controls Toxoplasma chronic differentiation. Nat. Microbiol. 8, 889–904 (2023).

74. Plattner, F. et al. Toxoplasma profilin is essential for host cell invasion and TLR11-dependent induction of an interleukin-12 response. Cell Host Microbe 3, 77–87 (2008).

75. Wichroski, M. J., Melton, J. A., Donahue, C. G., Tweten, R. K. & Ward, G. E. *Clostridium septicum*alpha-toxin is active against the parasitic protozoan*Toxoplasma gondii*and targets members of the SAG family of glycosylphosphatidylinositol-anchored surface proteins. Infect. Immun. 70, 4353–4361 (2002).

76. Schindelin, J., et al. Fiji: an open-source platform for biological-image analysis. Nat. Methods 9, 676–682 (2012).

77. Love, M. I., Huber, W. & Anders, S. Moderated estimation of fold change and dispersion for RNA-seq data with DESeq2. Genome Biol. 15, 550 (2014).

78. Altschul, S. F. et al. Gapped BLAST and PSI-BLAST: a new generation of protein database search programs. Nucleic Acids Res. 25, 3389–3402 (1997).

79. Schäffer, A. A., et al. Improving the accuracy of PSI-BLAST protein database searches with composition-based statistics and other refinements. Nucleic Acids Res. 29, 2994–3005 (2001).

80. Hassan, M. A., Vasquez, J. J., Guo-Liang, C., Meissner, M. & Nicolai Siegel, T. Comparative ribosome profiling uncovers a dominant role for translational control in Toxoplasma gondii. BMC Genomics 18, 961 (2017).

81. Letunic, I. & Bork, P. 20 years of the SMART protein domain annotation resource. Nucleic Acids Res. 46, D493–D496 (2018).

82. Markus, B. M., Waldman, B. S., Lorenzi, H. A. & Lourido, S. High-resolution mapping of transcription initiation in the asexual stages of Toxoplasma gondii. Front. Cell. Infect. Microbiol. 10, 617998 (2020).

83. Korotkevich, G. et al. Fast gene set enrichment analysis. bioRxiv 060012 (2016) doi:10.1101/060012.

84. Carpenter, A. E. et al. CellProfiler: image analysis software for identifying and quantifying cell phenotypes. Genome Biol. 7, R100 (2006).

85. Behnke, M. S., Zhang, T. P., Dubey, J. P. & Sibley, L. D. Toxoplasma gondii merozoite gene expression analysis with comparison to the life cycle discloses a unique expression state during enteric development. BMC Genomics 15, 350 (2014).

